# The conformational and mutational landscape of the ubiquitin-like marker for the autophagosome formation in cancer

**DOI:** 10.1101/635284

**Authors:** Burcu Aykac Fas, Mukesh Kumar, Valentina Sora, Maliha Mashkoor, Matteo Lambrughi, Matteo Tiberti, Elena Papaleo

## Abstract

Autophagy is a cellular process to recycle damaged cellular components and its modulation can be exploited for disease treatments. A key autophagy player is a ubiquitin-like protein, LC3B. Compelling evidence attests the role of autophagy and LC3B in different cancer types. Many LC3B structures have been solved, but a comprehensive study, including dynamics, has not been yet undertaken. To address this knowledge gap, we assessed ten physical models for molecular dynamics for their capabilities to describe the structural ensemble of LC3B in solution using different metrics and comparison with NMR data. With the resulting LC3B ensembles, we characterized the impact of 26 missense mutations from Pan-Cancer studies with different approaches. Our findings shed light on driver or neutral mutations in LC3B, providing an atlas of its modifications in cancer. Our framework could be used to assess the pathogenicity of mutations by accounting for the different aspects of protein structure and function altered by mutational events.

## Introduction

Autophagy is a highly conserved pathway in eukaryotes that allows the recycling of multiple cellular components during physiological conditions and in response to different types of stress such as starvation (Galluzzi et al., 2017; Gatica et al., 2018). Autophagy requires the sequestration of cytoplasm within double-membrane vesicles, known as autophagosomes, which then mature and fuse to lysosomes, leading to the degradation of their cargo (Parzych and Klionsky, 2013). This last step allows for the production of molecular building blocks that can be recycled and reused by the cell (He and Klionsky, 2009).

Autophagy selectively mediates the removal of damaged or unwanted cellular components (Fimia et al., 2013; Gatica et al., 2018; Stolz et al., 2014; Zaffagnini and Martens, 2016) and involves the binding of cargo receptors to the core autophagy machinery (Fimia et al., 2013; Gatica et al., 2018; Rogov et al., 2014; Stolz et al., 2014). Selective autophagy occurs in diverse forms depending on the target organelles (Kubli and Gustafsson, 2012; Zaffagnini and Martens, 2016) or cellular components. A common trait among different types of selective autophagy can be identified; a receptor protein needs to bind the cargo and to link it to the LC3/GABARAP family of Atg8 mammalian homologs (Maruyama et al., 2014; Schaaf et al., 2016) through the interaction with a scaffold protein (Gatica et al., 2018). An autophagy receptor is, by definition, a protein that can bridge the cargo to the autophagosomal membrane, thus promoting the engulfment of the cargo by the autophagosome (Johansen and Lamark, 2011; Stolz et al., 2014). Atg8 family members can also interact with another class of proteins, the autophagy adaptors. The adaptor proteins do not guide the engulfment and degradation of the cargo, but they work as anchor points or scaffolds for the autophagy machinery (Johansen and Lamark, 2011; Stolz et al., 2014).

One of the most studied LC3/GABARAP family proteins is LC3B, which is used as a marker for the assessment of autophagy activity in cellular assays (Kabeya, 2004; Klionsky et al., 2016; Mizushima et al., 2010; Tanida et al., 2008). The first study focused on the function of LC3B in autophagy was published in 2000 (Kabeya et al., 2000) and has gained more than 5000 citations ever since. LC3B structure is characterized by two α-helices at the N-terminal of the protein and a ubiquitin (Ub)-like core (Figure 1A and B) (Mohan and Wollert, 2018; Sugawara et al., 2004). LC3B is a versatile protein that serves as a platform for protein-protein interactions (Nakatogawa et al., 2007; Wild et al., 2014). LC3B and Atg8 proteins, more broadly, recruit the autophagy receptors through the binding of a specific short linear motif known as the LC3-interacting region (LIRs) in eukaryotes (Figure 1C) (Alemu et al., 2012; Kim et al., 2016; Klionsky and Schulman, 2014; Noda et al., 2010, 2008; Slobodkin and Elazar, 2013). The first LIR was discovered in the mammalian SQSTM1 (also known as p62) autophagy receptor in 2007 (Pankiv et al., 2007). The consensus sequence of the LIR motif includes an aromatic residue (W/F/Y) and a hydrophobic one (L/I/V), separated by two other residues (Alemu et al., 2012; Birgisdottir et al., 2013; Noda et al., 2008). Non-canonical LIR motifs have also been identified (Di Rita et al., 2018; Kuang et al., 2016a; von Muhlinen et al., 2012). The binding of the LIR to the LC3 proteins occurs at the interface between the N-terminal helical region and the Ub-like subdomain (Figure 1C). Interactions of biological partners with LC3/GABARAP can also be mediated at interfaces distinct from the LIR motif (Marshall et al., 2019; Wild et al., 2014). The LC3/GABARAP family members are not necessarily redundant in terms of functions, and each of them can be involved in different autophagy pathways (Lee and Lee, 2016; Schaaf et al., 2016), as illustrated in Figure 1D.

**Figure 1.**
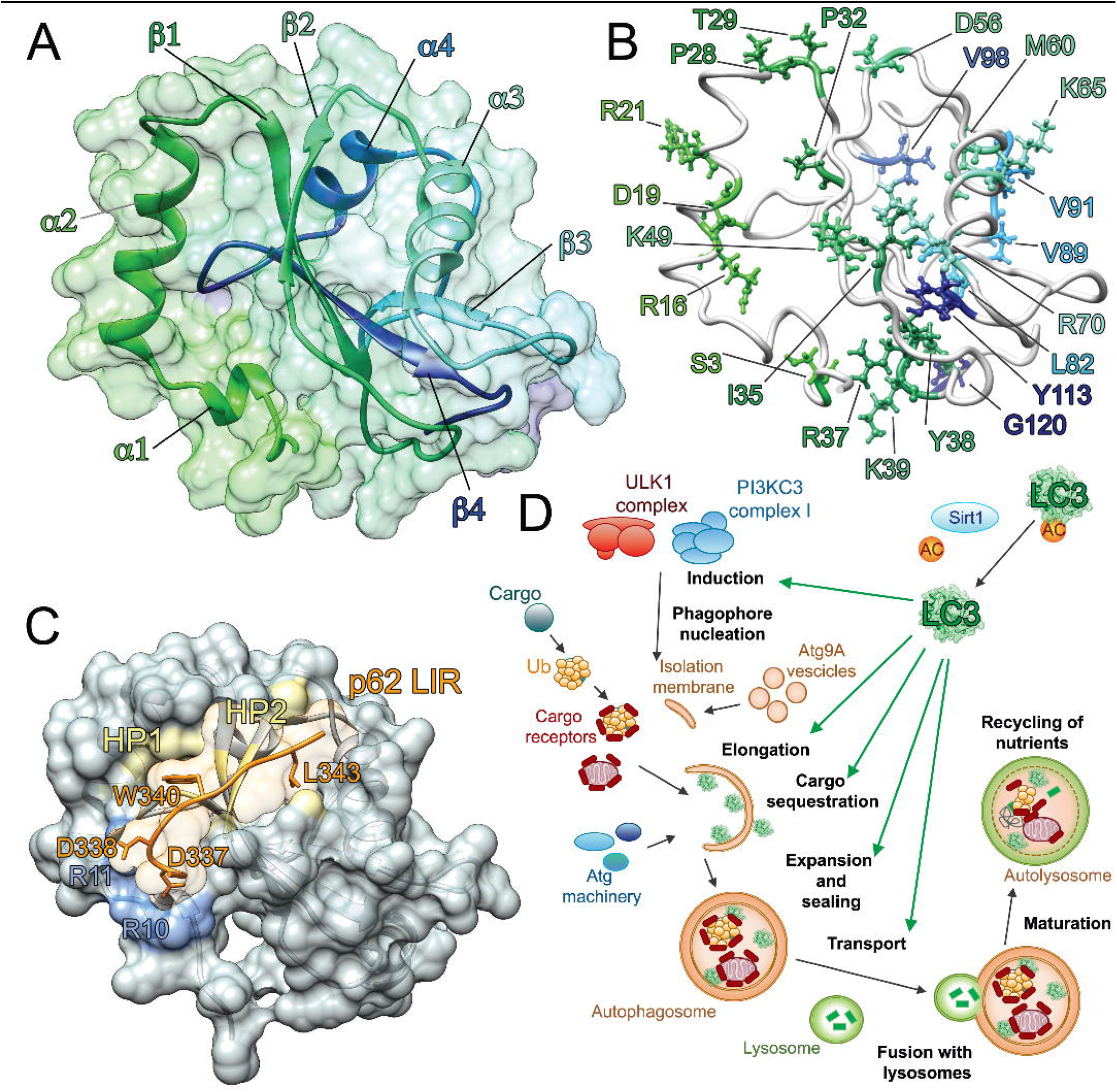
Structural properties and functional roles of the key marker of autophagy LC3B. A) LC3B is an ubiquitin-like protein, characterized by two α-helices at the N-terminal followed by an ubiquitin (Ub)-like core. The structure of LC3B (PDB entry: 1V49) is shown as cartoon and surface, using a color gradient from the N-terminal (green) to the C-terminal (dark blue). B) Localization in the structure of LC3B of the 28 residues identified as targets of missense mutations from cancer genomic studies. LC3B is shown as white cartoon while the target residues are indicated as colored stick, using same color gradient as panel A. C) LC3B mediates protein-protein interactions and recruits the autophagy receptors through the binding of a short linear motif, called LC3-interacting region (LIRs). The complex of LC3B (grey cartoon and surface) with the LIR motif of mammalian p62 (orange cartoon) is shown (PDB entry: 2ZJD). The important residues for the binding of p62 LIR motif to LC3B are indicated as sticks. The two hydrophobic pockets (HP1 and HP2) and the R10 and R11 in the binding interface of LC3B are indicated in yellow and light blue, respectively. D) LC3B is involved in different functions at several steps of the selective autophagy pathway.

Selective autophagy pathways are known to positively or negatively affect a large range of diseases including cancer (Mah and Ryan, 2012; Mancias and Kimmelman, 2016; Mizushima, 2018; Rybstein et al., 2018). These findings have attracted the attention of pharmaceutical and medical researchers as modulation of autophagy could provide means for new disease treatments and the identification of prognostic factors and disease-related markers (Thorburn et al., 2014; Towers and Thorburn, 2016). For example, in colorectal cancer, high LC3B expression levels correlate with improved survival after surgery (Wu et al., 2015), whereas in clear cell renal cell carcinomas, without the loss of the tumor suppressor gene *VHL*, high levels of LC3 proteins have been linked to tumor progression (Cowey and Rathmell, 2009; Mikhaylova et al., 2012) and the *VHL*-regulated miRNA-204 can suppress tumor growth inhibiting LC3B (Mikhaylova et al., 2012). In another study, the low protein expression of Beclin1 and LC3A/B contributed to an aggressive cancer phenotype in metastatic colorectal carcinoma (H. Zhao et al., 2017). A systematic review of cancer data found that high expression of LC3B predicts adverse prognosis in breast cancer (He et al., 2014) and oral squamous carcinoma (Tang et al., 2013). In a Pan-Cancer study on 20 cancer types, LC3B exhibited a common expression pattern in malignancy, whereby high LC3B expression was associated with proliferation, invasion, metastasis, high grades and an adverse outcome (Lazova et al., 2012). Similar expression patterns for LC3B have also been found in other independent studies on triple negative breast cancer (Zhao et al., 2013) and hepatocellular carcinoma (Wu et al., 2014). In summary, LC3B-associated autophagy is likely to promote or repress tumorigenic events depending on the context and tumor site (Schaaf et al., 2016). Autophagy can have protective functions against cancer development but can contribute to cancer progression and resistance to treatment. In this way, autophagy is often described as a double-edged sword in cancer (Kimmelman, 2011; Singh et al., 2018; Thorburn, 2014; White, 2012; White and DiPaola, 2009; Yao et al., 2016).

Apart from evidence regarding changes in the expression level of LC3B in different cancer types, genomic alterations of LC3/GABARAP family members and their relation to cancer have been poorly explored (Nuta et al., 2018). We thus decided to curate an atlas of missense mutations found in cancer samples for the LC3B autophagy markers and characterize them with a combination of structural and systems biology approaches. We aim at providing a framework that can be possible to extend to other Atg8 proteins and other cancer targets in general.

Several three-dimensional (3D) structures of LC3B have been solved by X-ray or NMR in the free state (Kouno et al., 2005; Rogov et al., 2013) or in complex with different biological partners (Ichimura et al., 2008; Jemal et al., 2011; Kuang et al., 2016b; Kwon et al., 2017; Lv et al., 2016; McEwan et al., 2015; Olsvik et al., 2015; Qiu et al., 2017; Rogov et al., 2017, 2013; Stadel et al., 2015; Suzuki et al., 2014; Yang et al., 2017). This provides a valuable source of information for investigations using biomolecular simulations and other structural computational methods of both the LC3B wild-type and its mutated variant. LC3 proteins have not been thoroughly explored by systematic biomolecular simulations or other computational structural methods. Among the simulations present in the literature, each one focused on specific aspects of LC3B dynamics using all-atom molecular dynamics (MD) simulations with a single force field (Jatana et al., 2019; Liu et al., 2013; Nuta et al., 2018; L. Zhao et al., 2017) or coarse-grained approaches (Thukral et al., 2015).

Here we aim to apply methods which combine conformational ensembles derived by molecular dynamics (MD) simulations and analysis with methods inspired by graph theory (Nygaard et al., 2016; Papaleo, 2015; Papaleo et al., 2016), to study the LC3B structure-dynamics-function relationship. We also aim to investigate the effects of missense mutations found in different cancer types to facilitate the classification of cancer driver and passenger mutations. As a first step, we aimed to choose the best physical description for LC3B by assessing ten different state-of-the-art physical models (i.e., force fields) for MD. The choice of the force field can affect the quality of the sampling in a system-dependent manner, and it is crucial to evaluate them when a new protein is under investigation (Dror et al., 2012; Klepeis et al., 2009). Most of the current force fields indeed differ for subtle substitutions in the torsional potential of the protein backbone and side chains, which can have a significant impact on the simulated structure and dynamics (Guvench and MacKerell, 2008; Lange et al., 2010; Lindorff-Larsen et al., 2012; Martín-García et al., 2015; Tiberti et al., 2015; Unan et al., 2015). This assessment has been extensively carried out for ubiquitin with different strategies (Bowman, 2016; F et al., 2012; Huang et al., 2017; Lange et al., 2010; Li and Brüschweiler, 2010a; Lindorff-Larsen et al., 2016, 2012; Long et al., 2011; Maragakis et al., 2008; Martín-García et al., 2015; Showalter and Brüschweiler, 2007; Sultan et al., 2018) but not for the Atg8 proteins.

With meaningful conformational ensembles in hand for LC3B, we turned our attention to the study of the impact of mutations identified in genomics Pan-Cancer studies, focusing on those which are unlikely to be natural polymorphisms in the healthy population. In our study, we accounted for the different aspects that a mutation could alter (Figure 2): i) protein stability, ii) interaction with the biological partners, iii) long-range communication between sites distant from the functional ones (which is often at the base of allostery), and iv) the interplay with post-translational modifications. We thus integrated the analysis of the MD ensembles with a range of other bioinformatic, network-based and structural approaches to achieve a comprehensive classification of the missense mutations in the LC3B coding region.

**Figure 2.**
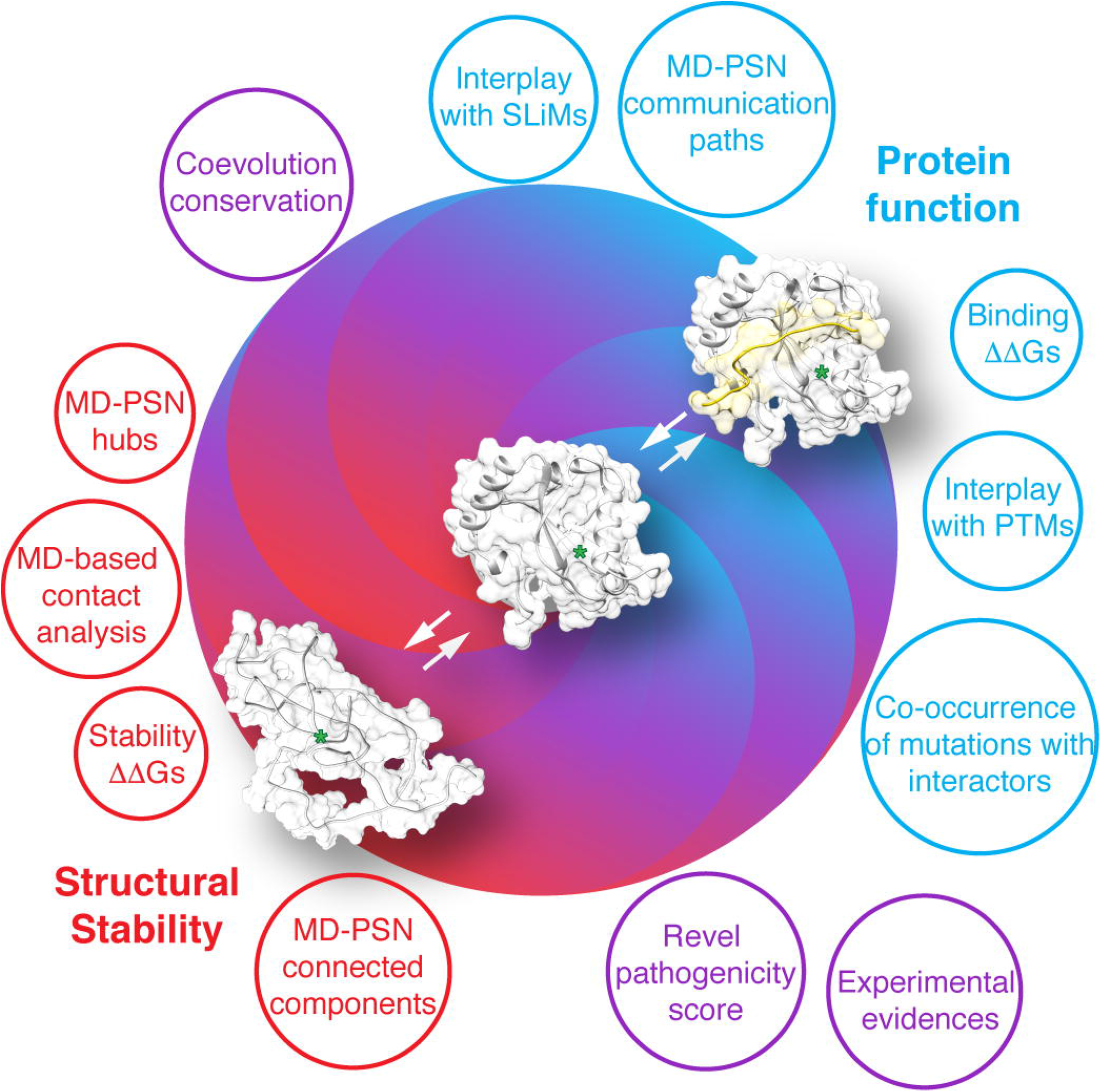
Schematic illustration of the methods and properties used for the assessment of the LC3B missense mutations from cancer genomic studies. We analyzed LC3B sequence, available structures of LC3B and its complexes with interactors, and LC3B MD-derived structural ensembles by using a combination of different approaches to account for the different layers of alterations that a mutation can induce at the level of: i) protein structural stability (i.e., contact analysis, stability ΔΔGs, shown in red); ii) protein function (i.e., binding ΔΔGs, interplay with SLiMs and PTMs, co-occurrence of mutations with interactors, shown in blue), iii) or implicitely accounting for both of them (i.e., Revel pathogenicity score, coevolution conservation, shown in purple).

## Results and Discussion

### State-of-the-art force fields for all-atom molecular dynamics consistently describe microsecond dynamics of LC3B

To compare the quality and sampling of the MD simulations of LC3B carried out with the ten state-of-the-art force fields selected for this study, we integrated different and complementary metrics.

At first, we estimated the atomic resolution (R) for each of the MD ensembles, as we recently did for another cancer-related protein simulated with different physical models (Nygaard et al., 2016). The predicted R-value allows for the collective quantification of different parameters for structural quality (Berjanskii et al., 2012) such as the population of side-chain dihedrals, rotamers that deviate from the “penultimate rotamer library” (Lovell et al., 2000), deviation from the allowed regions of the Ramachandran plot, packing of the protein core, hydrogen-bond networks and atomic clashes. After discarding the flexible N- (1-6) and C-terminal (117-120) tails from the analysis, the median R values for the different MD ensembles of LC3B_7-116_ were in reasonable agreement with each other, and all the ensembles featured R values mostly below the of resolution (Figure 3A). Notably, all the MD simulations provided conformational ensembles with R values better than the applied MD algorithm are refining the structure to a level close to the X-ray structure deposited in the entry 3VTU (R= 1.6, Å, (Rogov et al., 2013)). We performed a statistical pairwise Wilcoxon test to verify which of the MD ensembles provided a significantly different resolution distribution (see **Table S1**). Out of the force fields tested, RSFF1 had the highest median in proximity of the threshold, and ff99SB*-ILDN had lowest median values with most R values below the threshold. ff99SBnmr1, a99SB-disp, ff99SB*-ILDN-Q had the highest R values and were significantly different from the MD ensembles generated with the CHARMM force fields or ff99SB*-ILDN. LC3B has been studied by NMR spectroscopy, providing a number of probes of protein dynamics in solution, such as the assignment of the backbone chemical shifts (Kouno et al., 2005). Chemical shifts provide information about motions occurring on a heterogeneous range of time scales (Case, 2013; Li and Brüschweiler, 2010a; Palmer, 2015; Robustelli et al., 2012) and can be calculated from a MD ensemble of structures (Kohlhoff et al., 2009; Li and Brüschweiler, 2015, 2012). As chemical shifts are useful for experimental validation of the MD ensembles produced using different force fields (Beauchamp et al., 2012; Best et al., 2012b; Guvench and MacKerell, 2008; Henriques et al., 2015; Lange et al., 2010; Lindorff-Larsen et al., 2012; Martín-García et al., 2015; Nygaard et al., 2016; Papaleo et al., 2018, 2014b; Piana et al., 2014; Unan et al., 2015), we calculated the backbone chemical shifts from each MD ensemble and compared them to the experimental values. The calculated chemical shifts from our simulations are overall in fair agreement with the experimental data (**Table S2**) with an RMSE value below or close to the error of the prediction associated with each atom type (Li and Brüschweiler, 2015). The only relevant exceptions are RSFF1 and RSFF2 with RMSE values higher than 1.20 ppm for the Cα atoms. For LC3B in its unbound, we did not find available Cβ or side-chain methyl chemical shifts, which might have been more sensitive to changes in the intramolecular interactions described by the different force fields (Papaleo et al., 2018).

**Figure 3.**
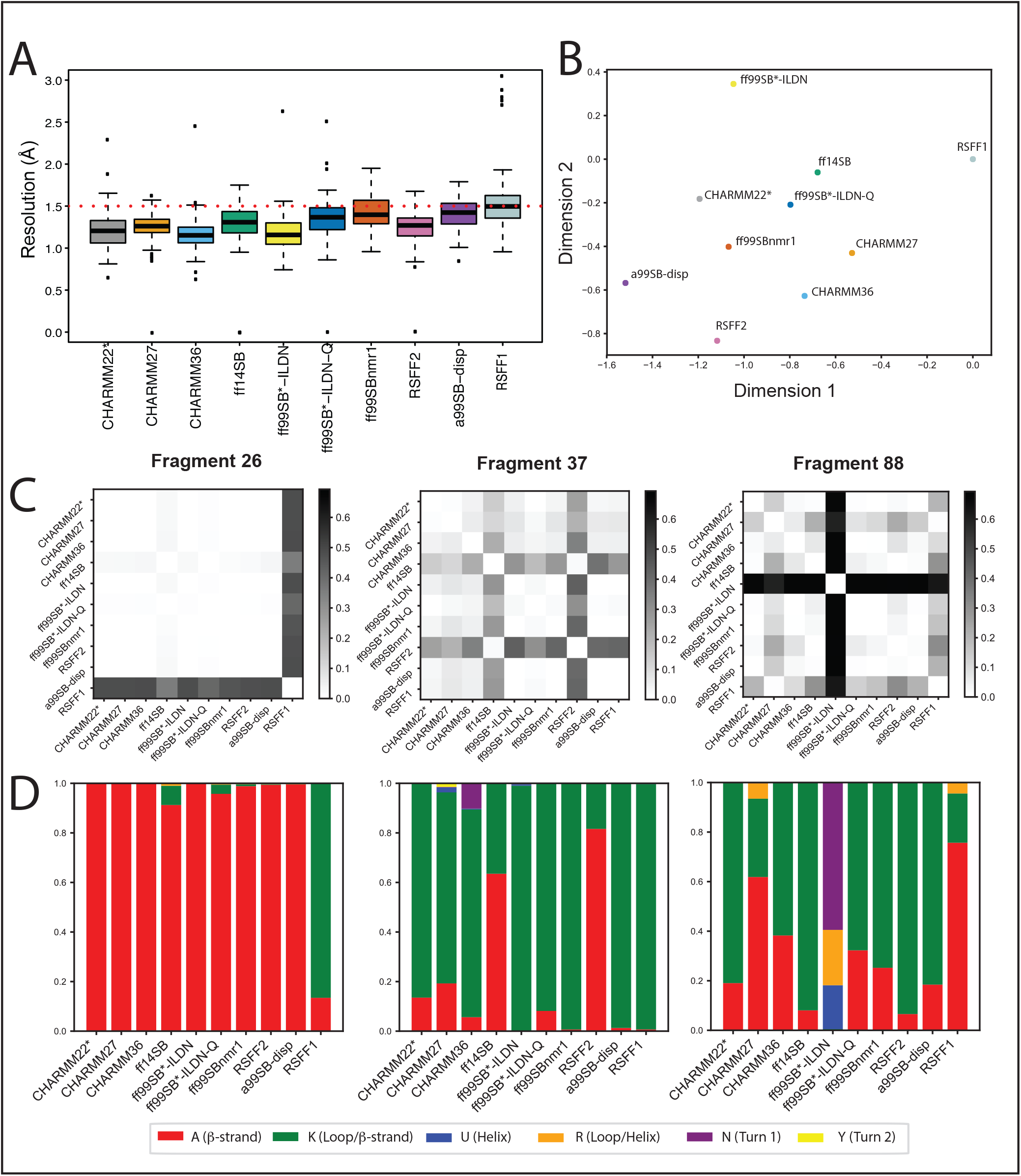
Comparison of MD ensembles of LC3B. A) Prediction of structural resolution values for different MD ensembles of LC3B generated using different force fields. All the MD force fields generally refine the structure to a resolution close to an X-ray structure deposited in the entry 3VTU and are in reasonable agreement with each other according to this parameter. The only (not pronounced) differences are observed for RSFF1, ff99SBnmr1, a99SB-disp and ff99SB*-ILDN, according to the statistical test reported in Table S1. B) Comparison of MD ensembles from different force fields using the clustering-based ensemble similarity (CES) method implemented in *ENCORE*. The two-dimensional plot for the clustering of the of the ten force field ensembles is based on the pairwise similarities calculated by CES using the tree preserving embedding method. The higher the distance between two force-fields is in the plot, the more different the sampling was between the respective simulations. The corresponding heatmap is provided in Supplementary File S2. The analysis shows that RSFF1, ff99SB*-ILDN, RSFF2 and a99SB-disp were the ensembles more distant from the others. C-D) These figures showcase the results obtained with the reduced structural alphabet (SA) method for three different SA fragments as representative example (i.e., fragments 26,37 and 88). The full datasets are available as Supplementary Files S3 and S4. Here, independently for each fragment, we have calculated the frequency of each letter in each force-field simulation, as shown in panel D. The frequencies of the samples have been used as an estimate of the underlying probability distributions and compared using the Jensen-Shannon divergence (dJS) between each pair of simulations, as shown in panel C. Higher values in this plot mean a more different sampling of SA letters between a given pair of force fields for a specific fragment.

As a complementary approach to evaluate the impact of different force fields on the description of the protein, the structural ensembles can also be compared in terms of the overlap of the conformational space sampled by the different simulations. Dimensionality reduction techniques can be used as part of this approach (Lange et al., 2010; Martín-García et al., 2015; Tiberti et al., 2015). At first, we used Principal Component Analysis (PCA) and focused on the projection along the first three principal components (PCs) (**Supplementary File S1)**. We found that in most of the simulations there is a region of general overlap but that the sampling also deviates from the starting structure (marked as a *). This is probably due to the low resolution of the initial structure deposited in PDB which the MD force fields refined, as confirmed by the lower R values and agreement with the chemical shifts.

Since the first three PCs explained only 35.5% of the total variance for the concatenated trajectory, we turned our attention to more quantitative metrics for comparison of the distribution underlying structural ensembles. We decided to use a structural clustering method (Clustering-based Ensemble Similarity, CES) implemented in *ENCORE* (Tiberti et al., 2015) to achieve a complementary assessment of the similarities between force fields and the sampled conformational subspaces. As revealed by the projections in Figure 3B and the heatmaps (**Supplementary File S2**), RSFF1, ff99SB*-ILDN, RSFF2 and a99SB-disp are the ensembles more distant from the others.

We then wondered if these differences could be due to local conformational variabilities in the structures sampled by each simulation. To this aim, we exploited the structural alphabet (SA) paradigm (Craveur et al., 2015) to estimate the sampling of local states in the simulations, relying on a reduced version of the M32K25 structural alphabet (Pandini et al., 2010) (A, K, U, R, N and Y, as detailed in Methods and Figure 3C). At first, we estimated the discrete probability distribution of the states for each fragment of the protein in the different MD ensembles. We used the Jensen-Shannon divergence (jsD) as a measure to compare the calculated probability distributions to have an estimate of the differences in the conformations of the MD ensembles (Figure 3C **and Supplementary File S3**). The SA analysis confirmed the diversity of the structures explored by RSFF1 in different areas of the protein, including the LIR-binding interface (see **Supplementary File S3**). The analysis also highlighted the different local structures sampled by ff99SB*-ILDN in the C-terminal subdomain of the protein (residues 82-105). Moreover, we observed local differences for other force fields, particularly CHARMM36, CHARMM27, ff14SB and RSFF2 (Figure 3D **as an example and Supplementary File S4**).

In summary, all the force fields provide a structural ensemble of reasonable quality for LC3B. However, using methods for comparison of ensembles of structures and their local states, we have been able to identify important local differences. Overall, CHARMM22* seems to be among the most robust force fields in the description of LC3B according to the properties here analysed. We thus selected it for further analyses to link the MD ensemble to functional properties and to study the mutation sites of LC3B found in genomic cancer studies.

### An atlas of LC3B missense mutations in cancer and their interplay with post-translational modifications and functional motifs

We retrieved 28 missense mutations identified in LC3B from cancer genomic studies (Figure 1B) and verified if there were possible natural polymorphisms in healthy individuals. In the *ExAC* database (Kobayashi et al., 2017), we identified only three mutations within our dataset (R37Q, K65E, and R70H) that had a very low frequency in the population (< 1/10000). We thus retained all 28 mutations in 23 residue sites found in 13 different cancer types (see **Supplementary File S5**). We analyzed the mutation sites in the context of their interplay with PTMs, overlap with functional short linear motifs (SLiMs), coupling and conservation upon coevolution-based analysis, and their potential to be pathogenic according to the *REVEL* score (Nilah M Ioannidis et al., 2016; Li et al., 2018). We also verified that the corresponding mutated transcript was expressed in the samples, through analysis of the corresponding entry in *Cbioportal* (Cerami et al., 2012). Most of the mutant LC3B variants are expressed with the exception of M60I and R70C, for which we cannot exclude additional effects due a marked low expression of the gene product. We verified (for those samples with information available) that the mutation was the only one targeting the LC3B gene in that specific sample. For most of the mutation sites, we also found an overlap with predicted SLiMs associated to different functions, regulatory modifications or interactors (Figure 4A).

**Figure 4.**
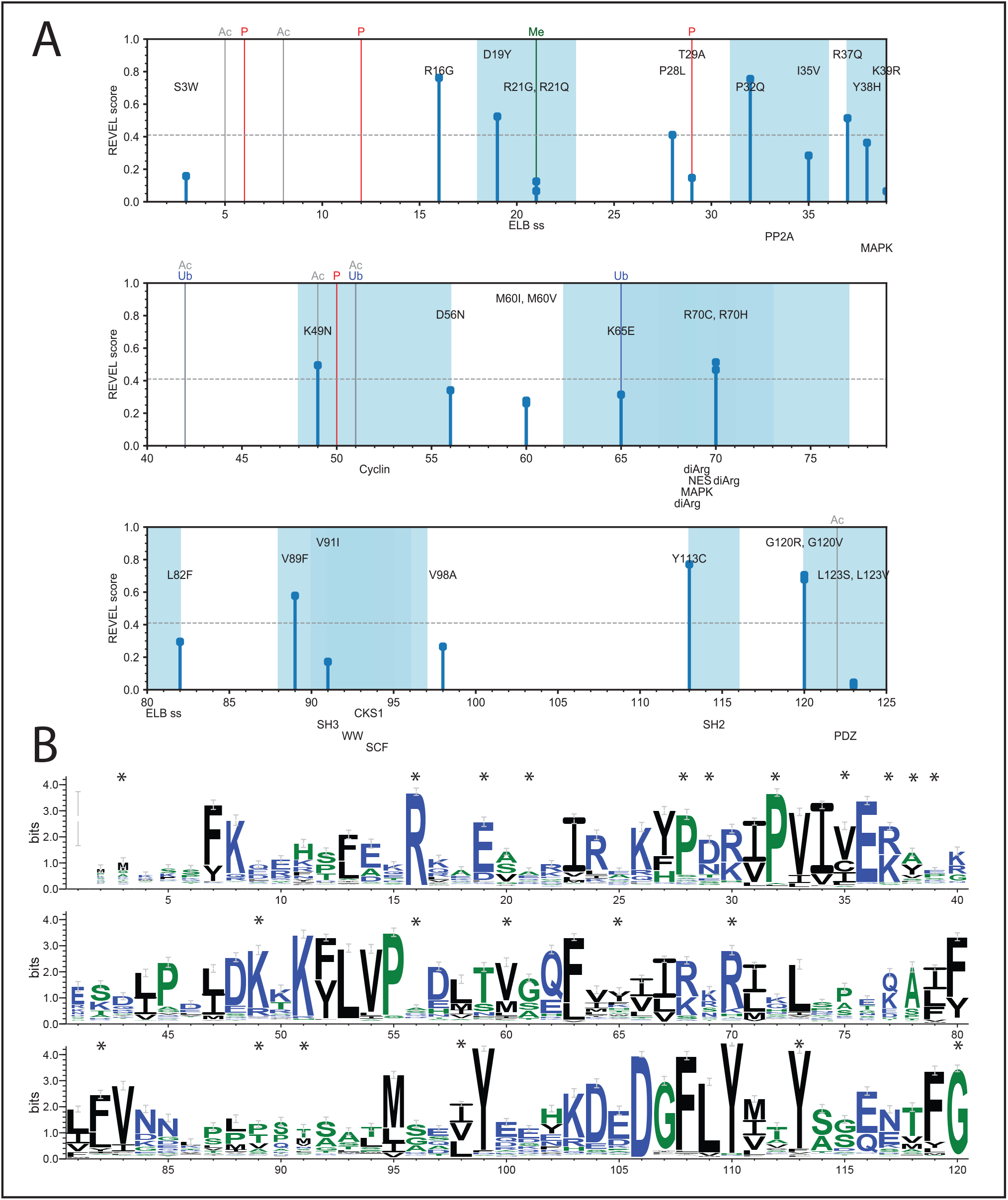
Sequence-based assessment of the LC3B mutations sites found in cancer genomic studies. A) Schematic representation of the identified cancer-related mutations and the analysis of: i) *REVEL* score, ii) overlap with PTMs and, iii) overlap with Short Linear motifs (SLiMs). In this plot, the amino acid sequence of LC3B, according to the main Uniprot isoform of the protein, runs on the X-axis. The missense mutations are shown as sticks, whose height is proportional to the associated *REVEL* pathogenicity score. PTMs from literature annotations are shown as vertical lines spanning the height of the plot (P for phosphorylation, Ubq for ubiquitination, Me for methylation, Ac for acetylation). SLiMs predicted for the protein sequence are shown below the plot, and the associated residue intervals colored in blue shades. Only the SLiMs overlapping with mutation sites are shown. The full set of data and annotation is provided as Supplementary File S5. We observed a substantial overlap with PTMs and functional motifs for the LC3B mutations under investigation and 11 mutations with a high pathogenicity score. B) The logo plot obtained by the multiple sequence alignment calculated by *Gremlin* is shown to evaluate the conservation and tolerated substitutions for the mutation sites of LC3B, which are marked by a *.

We carried out a first annotation of the potential pathogenic impact for each mutation using *REVEL*. We identified 11 likely pathogenic variants: R16G, D19Y, P28L, P32Q, R37Q, K49N, R70C/H, V89F, Y113C, and G120 R/V. We evaluated if the mutations could have a functional impact abolishing any of the SLiMs or PTMs, along with the likelihood of mutant variants harboring new PTM sites (Figure 4A). The mutation T29A is expected to abolish a phosphorylation site specific for the protein kinase C (Jiang et al., 2010). The analysis of the multiple sequence alignment generated by a coevolution method (Ovchinnikov et al., 2014) and the associated scores show that at this position negatively charged residues or other phosphorylatable residues (i.e., serine) are favoured with high conservation scores, emphasizing the functional importance of a negative charge at this position (Figure 4B). S3W, P32Q, and K49N are in the flanking region of T29 or other two phosphorylation sites (i.e. T6, (Jiang et al., 2010) and T50 (Wilkinson et al., 2015; Wilkinson and Hansen, 2015) and could impair the binding of the kinases/phosphatases. The R21G/Q, K49N and K65E mutations are predicted to abolish methylation (Figure 4A), acetylation (Huang et al., 2015) and ubiquitination sites (Wagner et al., 2011), respectively (Figure 4A). In particular, acetylation of K49 on the cytoplasmic form of LC3B is important for nuclear transport and the maintenance of the LC3B reservoir, deacetylation is, on the contrary, functional to the translocation to cytoplasm for interaction with the autophagy machinery (Huang et al., 2015; Huang and Liu, 2015). Deacetylation/acetylation cycles are important to maintain the proper pools of LC3B in the cell and a mutation impairing this modification, such as K49 could have detrimental effects increasing the cytoplasmic pool of LC3B and an uncontrolled autophagy.

D19Y is predicted to introduce a phosphorylatable residue for the TK and EGFR families of kinases by both *GPS* (Xue et al., 2008) and *NetPhos* (Blom et al., 2004). The residue is exposed on the protein surface and its substitution to tyrosine could make it available for post-translational modification, introducing a new level of regulation absent in the wild-type variant.

D19 is also tightly coupled to R16 according to the coevolution analysis (**Supplementary File S6**) and the only substitutions which are conserved are to glutamate and lysine (Figure 4B), respectively, suggesting an important contribution by charged residues at these positions, which might be compromised by the somatic mutation of arginine to glycine and of aspartate to tyrosine.

The position 35, where an isoleucine is located in the wild-type LC3B, shows propensity to allocate any hydrophobic aliphatic residue and, as a such, the I35V mutations are likely to provide the same features of the wild type variant (Figure 4B). A similar scenario occurs for the M60I and L82F mutations (Figure 3B). The arginine at the position 70 shows a lower conservation score than the histidine residue to which it is mutated in cancer (**Supplementary File S6**).

Most of the positions corresponding to the LC3B mutation sites are conserved in the other LC3 proteins (LC3A, LC3C and GABARAP) or with very conservative substitutions, especially when LC3A and LC3B are compared. The only marked differences are in R21 (highly variable site), T29 (where only LC3A retain the phospho-site), Y38 replaced by an alanine in GABARAP, K39 and D56 substituted by prolines or polar residues in LC3C and GABARAP, and K65 which is replaced by a serine and a phenylalanine in LC3C and GABARAP, respectively. We thus evaluated if LC3A, LC3C and GABARAP featured mutations similar to LC3B in cancer samples, using a similar approach as the one illustrated in Figure 4A. The mutation sites which featured the highest overlap in the four proteins were D19, P28, T29, P32, Y38, R70, and V89. Other sites were sensitive to mutations only in LC3B and A but not in GABARAP or LC3C. Interestingly, the R70C and H mutations are found in cancer samples in the corresponding regions of the other LC3 proteins, whereas the P28L and P32Q are found as somatic mutations in other members of the family, suggesting a sensitive hotspot for these regions. The surroundings of P32 and R70 have been also shown as important for the protein activity since the corresponding mutant variants in yeast were autophagy-defective (Nakatogawa et al., 2007).

The mutations D19Y and R21Q/G are also likely to impair the signal motif (i.e. Endosome-Lysosome-Basolateral sorting signals, ELB) to direct the protein to the endosome and lysosome compartments with a potential major impact on the autophagy pathway (Figure 4A). Several LC3B mutations could affect docking or recognition motifs for phosphatases or kinases, thus impacting on the regulation of LC3B activity and stability by its upstream regulators (**see Supplementary File S5**). Mutations at the residue G120 could abolish the cleavage site by ATG4B, impairing the activity of LC3B, as confirmed by experiments on the G120A mutant variant of LC3B (Kouno et al., 2005; Satoo et al., 2009).

### Assessment of the impact on protein stability upon LC3B missense mutations

A deeper understanding of the impact of the missense mutations on the protein can be achieved using structural methods (Nielsen et al., 2017; Nygaard et al., 2016; Riera et al., 2014; Scheller et al., 2019; Stein et al., 2019). A deleterious effect of a mutation could, for example, be to alter the protein structural stability, causing local misfolding and a higher propensity of the protein to be degraded in the cell. We thus estimated the changes in free energy associated to protein stability for each of the LC3B somatic mutations using an empirical energy function (Guerois et al., 2002; Schymkowitz et al., 2005). We implemented this procedure in a high-throughput manner (Nygaard et al., 2016; Papaleo et al., 2014a) so that all the possible mutations at each LC3B site can be assessed. Such high-throughput approach allowed us to evaluate the impact of all possible mutations in the protein without being limited to the mutations found in the cancer genomic studies. Indeed, we investigated, more broadly, if the cancer mutation sites are sensitive hotspots to substitutions and to predict if other sites of the protein when mutated could impact on its stability, providing groundwork for the prediction of LC3B somatic mutations which might arise from other cancer genomic studies or from the profiling of more cancer samples (Figure 5A). Our mutational scan shows that R16, P32, I35 and Y113 are sensitive hotspots for protein stability and they cannot tolerate most of the amino-acid substitutions. With regards to the atlas of somatic mutations found for LC3B, twelve of them are clearly destabilizing for the structural stability of the protein (R16G, R21G, P28L, P32Q, I35V, Y38H, R70C, L82F, and Y113C). Other mutations have mild or neutral effects with the exception of V89F which has been found with a stabilizing effect, likely to be due to an improved packing of the protein core (Figure 5B).

**Figure 5.**
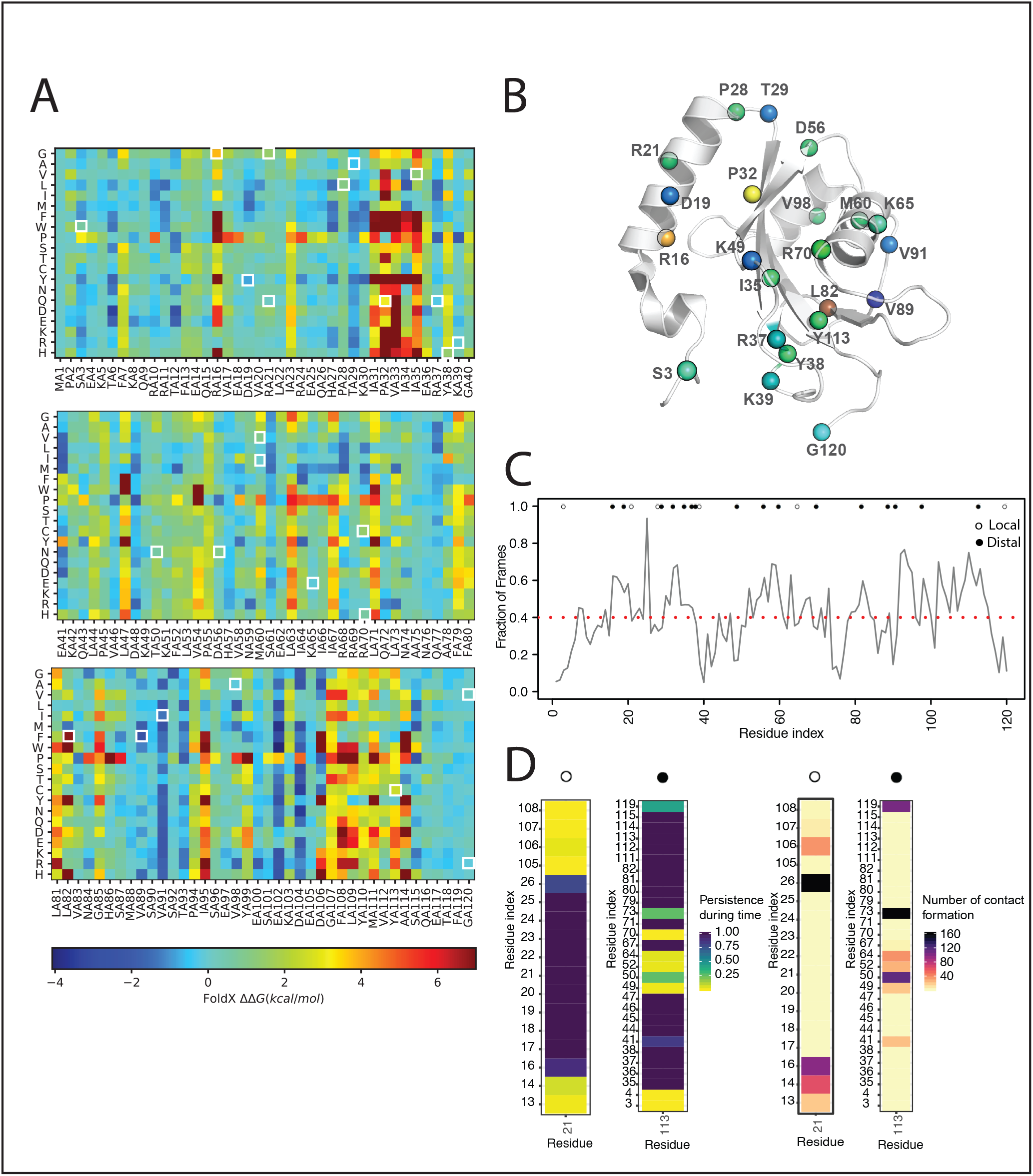
Assessment of the impact on stability upon missense mutations of LC3B found in cancer genomic studies. (A) The figure illustrated the heatmap of the saturation mutational scan performed using the 1V49 PDB entry of LC3B. Values of ΔΔG higher than 0 indicates destabilizing mutations. (B) Mapping on the 3D structure of LC3B (PDB entry 1V49) of the ΔΔG associated to protein stability of the 26 missense mutations of LC3B under investigation. In case of different mutations at the same site the highest ΔΔG is shown. The the mutations wer color-coded according to the corresponding heatmap color for sake of clarity. C-D) The contact-based analyses on the LC3B MD-derived ensemble with the CHARMM22* force field is illustrated in panels C and D and Supplementary File S13. The analyses were carried out with *CONAN*. C) The average local interaction time (avLIT) profile of the LC3B points out several mutation sites (R16, D19, P32, L82, V98 and Y113), with avLIT values higher than the mean of the distribution (0.4 fraction of the simulation frames), as possible sensitive hotspots for protein stability. D) Persistence and number of encounters in the CHARMM22* MD ensemble for each interaction of the mutation site R21 and Y113, as examples. The persistence of each contact is normalized on the total time length of the simulation and represented with a color gradient, from low (yellow) to high (blue) persistence. The number of formation events for each contact is represented with a color gradient, from low (light brown) to high (black). Analogous plots for each residue in the LC3B are reported in Supplementary File S13. We identified two classes of mutation sites: local and distal, indicated with white and black dots, respectively. The “local” class accounts for mutation sites (e.g. R21) forming strong atomic contacts with only residues contiguous in the sequence and they are mostly predicted to be neutral for stability. The “distal” class accounts for mutation sites (e.g. Y113) forming strong atomic contacts with also residues distant in the sequence and they are mostly predicted to be relevant for stability.

To achieve a better understanding of the role of these residues in the structural stability of LC3B, we also employed a method to estimate atomic contacts (Mercadante et al., 2018) and their lifetime in the CHARMM22* MD ensemble (see Materials and Methods). We calculated the average local interaction time (avLIT) for each residue of the protein during the simulation (Figure 5C). The mean of the distribution of avLIT values is of 0.4 (fraction of frames) and the somatic mutations sites with avLIT values higher than this threshold were R16, D19, P32, L82, V98 and Y113, reflecting the results of the mutational scan above. We then estimate the strength and location of the interactions for each mutation site over time, as well as the associated number of encounters (Figure 5D). Two macro-groups can be identified, i.e. mutation sites with only atomic contacts with residues contiguous in the sequence and mutation sites involved also in strong or more dynamic contacts with residues distant in the sequence. In the first class, we found mostly the mutation sites predicted neutral for stability, whereas the second group account for residues such as D19, P32, L82, and Y113.

### Local effects of the mutations on binding of LIR motifs and other interactors

LC3B interactome is large (Wild et al., 2014) and the main function of LC3B is to recruit many different proteins to the autophagosome. Thus, to better appreciate the effect of the mutations identified in the cancer samples, one should also consider in the same samples the spectrum of alteration of the LC3B interactors. To this goal, we retrieved LC3B interactors mining the *IID* protein-protein interactions database (Kotlyar et al., 2019) and integrated them with interactors reported in a recent publication (Wild et al., 2014). We identified overall 95 LC3B partners and for each of them we verified if a LIR motif was reported in the literature as experimentally validated. For cases where no information was available on the mode of interaction, we predicted putative LIRs with *iLIR* (Jacomin et al., 2016).

We identified 70 interactors as either experimentally validated LIR-containing proteins or having a predicted LIR motif with a significant score by the predictor. We then retrieved the mutational status of the LIR-containing interactors in the same samples where the LC3B missense mutations were identified to explore the possibility of co-occurrence of mutations. For each of them, we evaluated if the mutation was in the proximity of the experimentally validated or predicted LIRs (Figure 6A-B). We found 39 mutations in 27 LIR-containing interactors occurring in samples where LC3B was mutated (**highlighted in red in** Figure 6A). In particular, 17 of these mutations were truncation with the potential of abolishing all or most of the LIR motifs in the interactors (Figure 6B). The remaining mutations were located in the core motif or in its proximity, along with in the N- and C-terminal regions (Figure 6B). We noticed that the mutations were in the proximity of the LIR region or in the core motifs affected mostly charged residues, which are likely to stabilize the binding of the wild-type complexes through changes in the electrostatic interactions.

**Figure 6.**
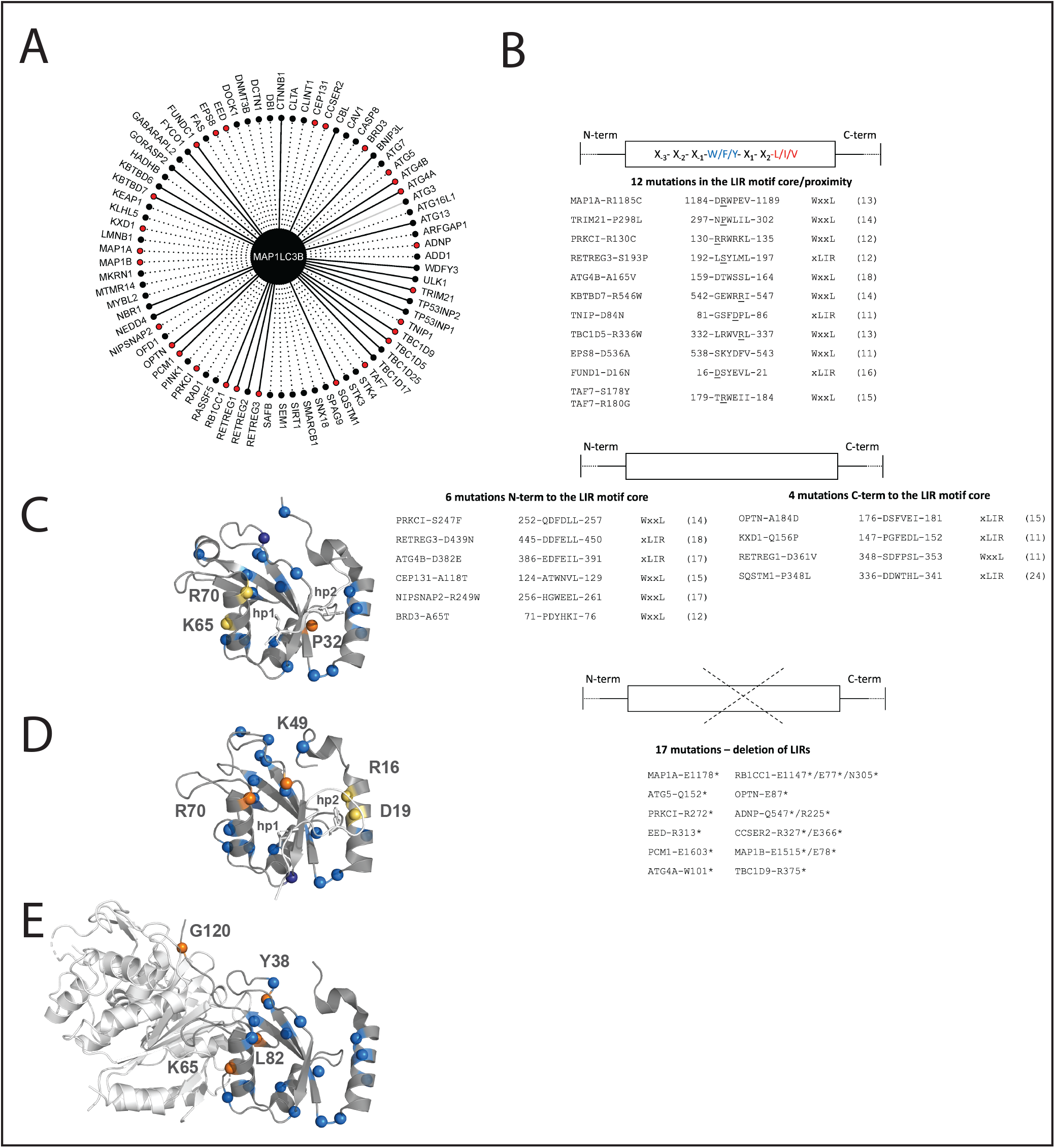
Co-occurrence of mutations in LC3B and LIR-containing interactors and local effects of mutations on intermolecular interactions. A) Network of the 70 LIR-containing proteins that interact with LC3B. The plot shows all the LC3B interactors which harbor an experimentally validated (solid line) or predicted LIR with PSSM score > 11 (dashed line). The 39 mutations found in 27 LIR-containing proteins in the proximity or within the LIR motif and co-occurring with LC3B mutations are highlighted in red. Only in our case, we identified an experimentally validated interactions for which no LIR motif could be predicted within the significance threshold used for the prediction, i.e. ATG3, which is showed as a grey edge. B) Among the 39 co-occurring mutations, 12 were located in the LIR core motif or its proximity (top figure), 6 and 4 were N- and C-terminal to the LIR motif core (middle figure), and 17 resulted in truncations abolishing all or most of the LIR motifs of the interactors (bottom figure). C-E) We estimated the local changes induced on the binding between LC3B and its interactors using a mutational scan based on an empirical energy function. We estimated the changes on binding free energies using, as references, two different LIR-binding modes (C, LC3B-p62 complex, PDB entry 2ZJD and D, LC3B-FUNDC1, PDB entry 2N9X) and a non-LIR interactor (E, LC3B-ATG4B complex, PBD entry 2ZZP). The mutation sites are showed as spheres which are colored according to the ΔΔG value.

For two of the interactors (OPTN and FUNCD1), we used the known 3D structures of the complexes with LC3B to estimate the changes in binding free energy associated to the cancer mutations of the LIR in co-occurrence with LC3B mutations: A184D of OPTN and R37Q of LC3B, along with D16N of FUNDC1 and R70C of LC3B. We identified a slightly negative binding ΔΔG (average ΔΔG =-0.37 kcal/mol) due to the combination of A184D (OPTN) and R37Q (LC3B) and a destabilization (average ΔΔG =1.41 kcal/mol) induced by D16N(FUNDC1) and R70C (LC3B), respectively. In parallel, we estimate the changes in binding free energies also with another protocol based on *Rosetta* (Barlow et al., 2018) for a reciprocal control of the calculation results, as recently suggested for similar applications (Buß et al., 2018). The calculation suggests that only D16N(FUNDC1)-R70C(LC3B) has a destabilizing effect on binding affinity, whereas A184D(OPTN)-R37Q(LC3B) has neutral effects according to *Rosetta* scan.

Moreover, we estimated local effects induced by the LC3B mutations on the binding to their partners of interaction calculating the changes in binding free energies for the complexes of LC3B-p62 (Ichimura et al., 2008) and LC3B-FUNDC1 (Kuang et al., 2016a), as prototypes of two different binding modes (Figure 6C-D). Most of the mutations have neutral effects on the local binding. We observed LIR-specific effects for the remaining mutations, i.e., P32Q and R70C as destabilizing mutations for the p62-LC3B complex, whereas K49N and R70C/H affected the LC3B binding with the FUNDC1 LIR. These results are also confirmed by the mutational scan with another method based on the *Rosetta* energy function (**Supplementary File S8**). The results were in agreement with the decreased binding of LC3B to p62 and FYCO LIRs upon mutations of LC3B R70 to alanine (Ichimura et al., 2008; Olsvik et al., 2015). D19N was recently characterized in the context of the binding of FUNDC1 with LC3B (Kuang et al., 2016a). The minor effect that we observed for this mutation on the unphosphorylated FUNDC1-LC3B is in agreement with the fact that D19 is located in proximity of the LIR phospho-site Y18 of FUNDC1 LIR and it is responsible for the stabilization of the phosphorylated state of FUNDC1, a function that is likely abolished by this cancer mutation, but that does not have a marked effect on the binding of the unphosphorylated LIRs. We thus assessed the impact of this mutation (D19N) on the structure of the phosphorylated FUNDC1-LC3B (Kuang et al., 2016a) complex and we estimated a higher destabilizing effect with a ΔΔG of approximately 1.14-1.27 kcal/mol.

K49N is also predicted to have destabilizing effects on the binding of the FUNDC1 LIR with LC3B (Figure 6D). K49 is especially important since it is a gatekeeper that regulates the binding of the LIR to the LC3/GABARAP pocket and undergoes conformational changes upon binding (Suzuki et al., 2014).

LC3B doesn’t interact with LIR motifs exclusively, but also with other proteins of the core autophagy machinery, such as Atg4B (Satoo et al., 2009). We estimated the changes in binding free energy upon mutations also for this protein complex (Figure 6E). A group of mutations (Y38H, K65E, L82F and G120V/R) specifically altered the interaction with Atg4B and did not affect the ones with the LIR motifs, as also shown by *Rosetta Flex ddG* (**Supplementary File S8**). Interestingly, L82 has been shown to be an essential residue for the C-terminal cleavage of LC3B (Fass et al., 2007), supporting our predictions. Y113C was also showed experimentally with a functional impact on the interaction with Atg4B (Nuta et al., 2018), but it is not identified as destabilizing for the interaction by our local mutation scan. According to our mutational scan on protein stability, this could be due to the fact that Y113C has an impact on structural stability (Figure 5B). Thus, a Y113C LC3B variant could result in protein variant more prone to proteasomal degradation. The altered biological readout might thus be a consequence of lower levels of this protein variant in the cell.

### LC3B ensemble under the lens of protein structure network: hubs for stability and long-range induced effects

The mutational scan described in the previous section only captures local effects for mutations in residues in the proximity of the interface. We thus used the Protein Structure Network (PSN) framework combined to MD simulations (Papaleo, 2015) to assess more distal effects. PSN-based methods provide is a solid framework to study protein structures (Angelova et al., 2011; Di Paola and Giuliani, 2015; Ghosh and Vishveshwara, 2008; Mariani et al., 2013; Verkhivker, 2019; Vuillon and Lesieur, 2015). We recently applied it to the study of other cancer-related proteins, allowing to shed light on the underlying structural communication among distal sites (Invernizzi et al., 2014; Lambrughi et al., 2016; Papaleo et al., 2012a; Sora and Papaleo, 2019) or to help in the classification of potential cancer passenger and driver missense mutations (Nygaard et al., 2016).

In details, the PSN employs the graph formalism to identify a network of interacting residues in a given protein from the number of non-covalent contacts in the protein or other intramolecular interactions (Tiberti et al., 2014; Viloria et al., 2017). Two main properties of a PSN are the hub residues, i.e., residues that are highly connected within the network and the connected components, i.e., clusters of residues which are inter-connected but do not interact with residues in other clusters. Hubs in a PSN could have both the role of shortening the communication between distal residues or they can have a structural role thanks to their contribution to the robustness of the network. Indeed, substitutions occurring on the nodes with small degree are likely not to have a large effect on the network (and thus the structure) integrity. On the contrary, if hubs are altered, the network integrity can be compromised. We calculated three PSNs, based on sidechain contacts (Tiberti et al., 2014; Viloria et al., 2017), hydrogen bonds (Tiberti et al., 2014) and salt-bridges (Jónsdóttir et al., 2014; Tiberti et al., 2014), respectively.

We then calculated the hubs and connected components from the contact-based and hydrogen-bond based PSNs from the CHARMM22* simulation of LC3B. Among the LC3B cancer mutation sites, I35, Y38, and V89 have a hub behaviour in the contact-based PSN (Figure 7A). These residues also belong to the second connect component of the contact-based network together with other hydrophobic residues, highlighting their importance for protein stability (Figure 7B). Moreover, many mutation sites are located at hub positions in the hydrogen bond network (i.e., R16, D19, R21, Y38, M60, K65, R70, V98 and Y113) (Figure 7C-D). Overall, due to their pivotal role to mediate different classes of intramolecular interactions, and the introduction of substitutions which would not conserve these interactions, the mutations R16G, D19Y, R21G, Y38H, R70C, Y113C are likely to impact on the structural stability of LC3B.

**Figure 7.**
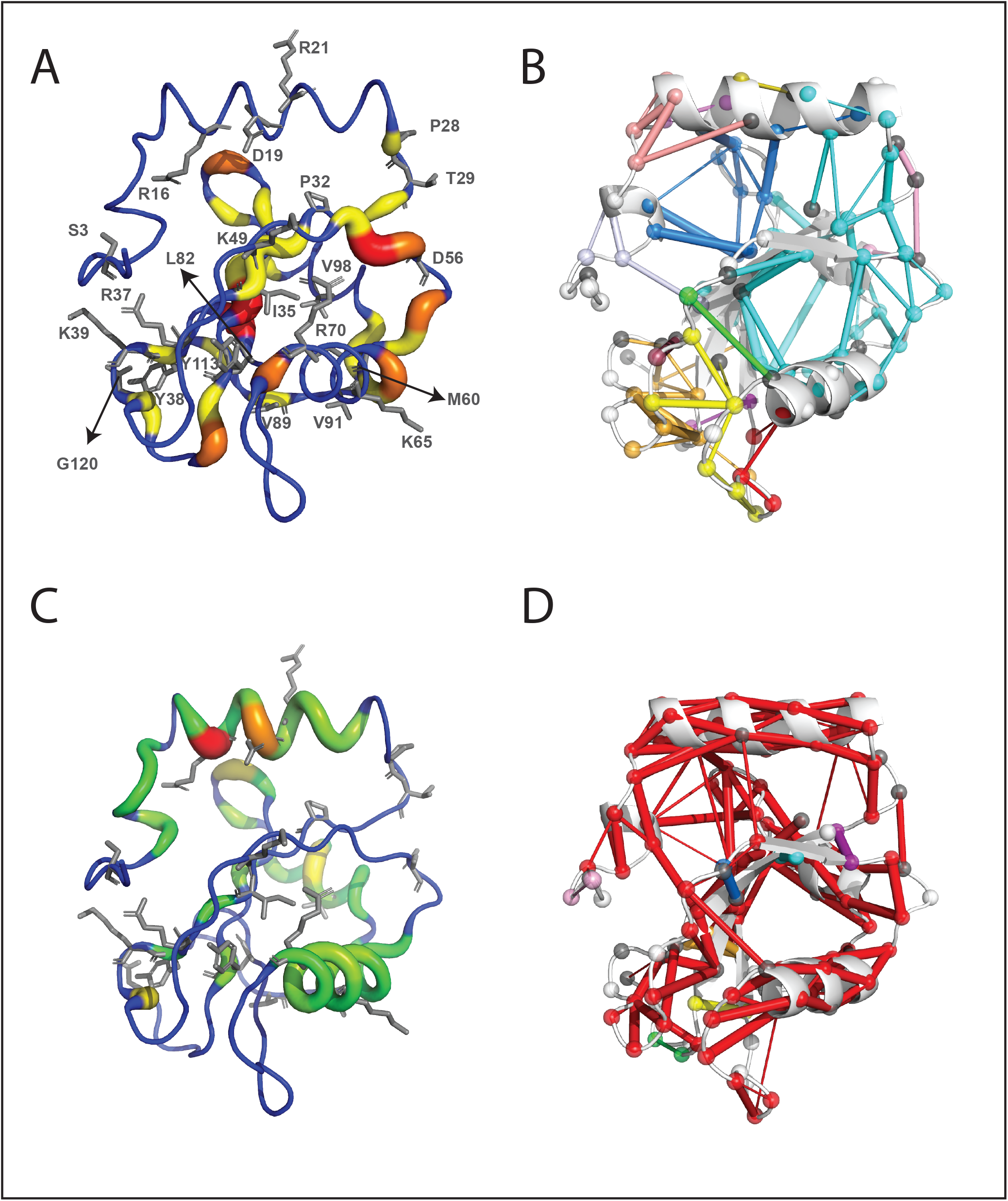
Assessment of long-range effects induced by LC3B missense mutations found in cancer genomic studies. Schematic representation of hubs and connected components for the PSN based on side-chain contacts (A-B) and hydrogen bonds (C-D), calculated from the CHARMM22* MD ensemble of LC3B. A-C) Hubs are residues that are highly connected within a protein structure network (PSN). The hubs showed in the figure are color-coded with green (node degree = 3), yellow (4), orange (5), dark orange (6) and red (7), the ribbon thickness indicates the node degree of each hub. B-D) Connected components are clusters of linked nodes with no edges in common with the nodes belonging to other clusters of the PSN. The results showed in panels A and B refer the contact-based PSN calculations based on a 5.125 Å cut-off.

Since 11 mutations where in charged residues of LC3B, we also calculated the network of electrostatic interactions between positively (arginine and lysine) and negatively charged (aspartate and glutamate) residues in the MD ensemble (**Supplementary File S9**). D19 is central to a small network of salt bridges with K51 and R11 (which is important for the LIR binding), R16 is on one side of the four-residue network with D106, K8 and D104, which constraint a loop of the protein. K65 and R21 are only involved in local intra-helical salt bridges, whereas R70 shows a persistent interaction with D48. The other charged mutation sites have either low persistent electrostatic interactions or they are not involved in salt bridges and in solvent exposed positions.

To predict effects promoted from distal sites to the LIR binding region, we also calculated the shortest paths of communication from the contact-based PSN between each of the mutation sites and the LIR binding interface, i.e., R10, R11 (Ichimura et al., 2008; Kirkin et al., 2009), K49, K51, L53, H57 and R70 (Ichimura et al., 2008; Kirkin et al., 2009; Olsvik et al., 2015), which could be disrupted or weakened by the mutations (Table 1). We identified long-range communication between the LC3B mutation sites only for the interface residues K49, H57 and L53. We observed that I35 is crucial for the communication from the core of the protein to the LIR binding interface at several different sites, spanning different areas of the LIR binding groove. I35 is also often intermediate residue in other paths mediated by different mutation sites (Table 1). In particular, we identified a strong short path of communication between the gatekeeper residue 49K at the LIR binding site to the mutation site 35I (average weight = 64.7) which might be weakened when I35 is mutated to a shorter side-chain as valine. On the opposite site of the binding pocket, I35 can, acting through a longer path, trigger conformational changes to the L53 and H57 binding site residues (**Table 1**). K49, apart from playing an important local role in mediating the LIR binding, can also communicate long range with H57 on the LIR binding groove important for the binding of the C-terminal part of the LIR. A similar behaviour is observed for M60 and K65 which are pivotal for long range communication to all the three LIR binding sites (K49, L53 and H57, Table 1). In addition, K65 and I35 are part of the same long-range communication spine from the surface of the protein to the LIR binding interface, suggesting that two site communication can occur between the Atg4B binding site to which K65 belongs and the LIR binding site. Other potential long-range effects can be exerted by P32 to H57 (passing also through L53), along with the three valine residues at positions 89, 91 and 98, which seems to act more as intermediate nodes of more complex paths (Table 1).

**Table 1.** Shortest paths of communication between the mutation sites and the residues at the interaction interface between the LIR and LC3B in the MD ensembles derived by the CHARMM22* force field. Only communication to K49, L53 and H57 was identified for the mutation sites: P32, I35, K49, M60, K65, V89, V91 and V98. * This position is both a LIR-interacting site and a mutation site.

| MUTATION SITE | END NODE: K49 | END NODE: L53 | END NODE: H57 |
| --- | --- | --- | --- |
| P32 | L53-I23-P32 (av.weight = 41.2) |  | H57-P55-K30-L53-I23-P32 (av.weight=43.8) |
| I35 | K49-F52-I35 (av.weight= 64.7 ) | L53-K30-P55-V54-V33-M111-I35 (av.weight= 44) | H57-V58-V54-V33-M111-I35 (av.weight=41.1) |
| K49* |  |  | H57-V58-V54-V33-M111-I35-F52-K49 (av.weight=47.8) |
| M60 | K49-F52-I35-M111-V83-V89-M60 (av.weight = 53.1) | L53-K30-P55-V54-V33-M111-V83-V89-M60 (av.weight=51.6) | H57-V58-V54-V33-M111-V83-V89-M60 (av.weight=50.7) |
| K65 | K49-F52-I35-M111-V33-V54-V58-E62-K65 (av. weight= 45.9) | L53-K30-P55-V58-E62-K65 (av. weight=45.5) | H57-V58-E62-K65 (av.weight=38.7) |
| V89 | K49-F52-I35-M111-V83-V89 (av.weight =56.3) | L53-K30-P55-V54-V33-M111-V83-V89 (av.weight=53.7) | H57-V58-V54-V33-M111-V83-V89 (av.weight=52.9) |
| V91 | K49-F52-I35-M111-V33-V54-V58-E62-S61-V91 (av.weight=48.3) | L53-K30-P55-V58-E62-S61-V91 (av.weight=49.3) | H57-V58-E62-S61-V91 (av.weight=46.0) |
| V98 | K49-F52-I35-M111-V83-V89-V98 (av.weight=60.6) | L53-K30-P55-V54-V33-M111-V83-V89-V98 (av.weight=57.2) | H57-V58-V54-V33-M111-V83-V89-V98 (av.wegth=57.8) |

For all these residues that we found critical in distal communication, we speculate that their mutations could alter the native structural communication of LC3B protein from distal sites to the binding sites for different interactors.

### Classification and impact of LC3B missense mutations

We integrated all the data collected in this study to map the different effects that the cancer-related mutations of LC3B could exert, providing a comprehensive view of the many aspects that they can alter and that are ultimately linked to protein function at the cellular level, i.e. protein stability, regulation, abolishment/formation of PTMs or functional motifs for protein-protein interactions, local and distal effects influencing the binding to the partner of interactions, along which co-occurrence of mutations in the same cancer samples for the protein and its ligands (Figure 8A). We then ranked the mutations according to the properties that they alter to help in the classification of potential damaging and neutral ones (Figure 8B). The ranking allowed the selection of mutations to prioritize for validation of their “driver” or “passenger” potential, along with planning the proper experimental readout for the validation. For example, if a mutation is predicted damaging in relation to stability, experiments tailored to estimate the cellular protein level and half-life could be used as we recently did for other disease-related proteins (Nielsen et al., 2017; Scheller et al., 2019). On the other side, if the impact is more related to the protein activity or introduction/abolishment of functional motifs, binding assays for example based on peptide arrays, isothermal titration calorimetry, NMR spectroscopy, co-immunoprecipitation can be used together with assays of the related biological readouts in cellular models as for example we recently combined to the study of a LIR-containing scaffolding protein (Di Rita et al., 2018). Assays with upstream modifiers, such as ubiquitination or phosphorylation assays can instead be used to validate the interplay with PTMs, both in the direction of introduction of new layers of regulation upon mutation or their abolishment.

**Figure 8.**
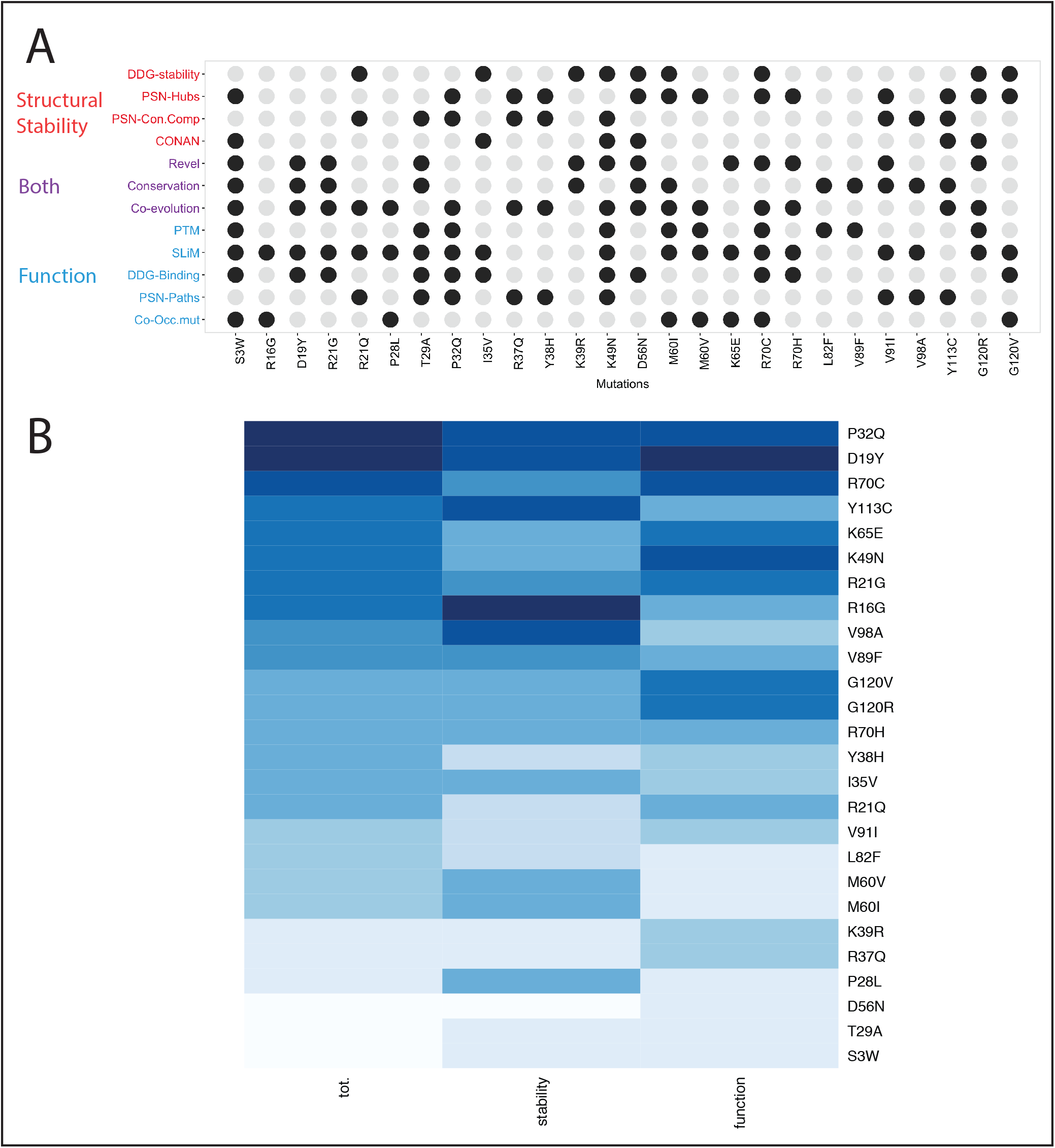
Classification of LC3B mutations found in cancer genomic studies. A) The different analyses carried out in our study have been aggregated to associate the potential of damaging (driver) or neutral (passenger) effect of each mutation. We used descriptors that account for protein stability (red), function (blue) or implicitly for both (purple). Mutations altering one of these properties are highlighted as black dots. The diagram in panel A allow to link each mutation to a specific effect which could guide the selection of the best set up for experimental validation and further studies. B) The heatmap is the results of a ranking on a collective score of damaging potential for the mutation (first column). The results for the ranking only according stability and function is showed as a reference in the second and third column of the heatmap. Darker the color more damaging the mutation is.

In our case study, we identified three potential classes of driver mutations: i) mutations that alter both stability and activity (R16G, D19Y, R70C, P32Q, Y113C, and R21G); ii) detrimental mutations for protein stability (R16G, V98A, V89F); iii) mutations neutral for the stability but altering the activity (G120R, G120V, and K49N).

We then searched in literature for experimental data that could validate our predictions, and we found results in agreement with the functional impact for mutations at R70, D19, G120, and K49, supporting our results. Mutations at R70 showed no accumulation of the pro-forms for LC3B (Liu et al., 2013), slower kinetics for Atg4B-mediated cleavage (Costa et al., 2016) and reduced binding for more that 20 interactors (Behrends et al., 2010; Kraft et al., 2014; Olsvik et al., 2015). G120 is fundamental for a proper C-terminal cleavage, which is impaired when this glycine is mutated to alanine (Tanida et al., 2004) and also G120 substitution with alanine has been shown to impair the binding of lamin B1 with LC3B (Dou et al., 2015). Y113C was recently shown to inhibit the enzymatic activity of Atg7 (E1-like enzyme) but not the E2-like activity (Nuta et al., 2018). Mutations at K49 alter the binding to the phosphorylated variants of the LIR-containing LC3B interactor, FUNDC1 (Lv et al., 2016), whereas if this residue is mutated to alanine can increase the binding of another LIR-containing protein, i.e. Nix (Rogov et al., 2017). D19N altered the selectivity for phosphorylation and unphosphorylated variants of FUNDC1 (Kuang et al., 2016a).

## Materials and Methods

### Molecular Dynamics Simulations

The molecular dynamics (MD) simulations for the human LC3B protein were performed in explicit solvent using *GROMACS* version 4.6 (Hess et al., 2008). One- μs MD simulations of LC3B monomer were collected starting from the free state NMR structure with PDB entry 1V49 (Kouno et al., 2005). We have employed ten force fields (ff14SB (Maier et al., 2015), ff99SBnmr1 (Li and Brüschweiler, 2010b), ff99SB*-ILDN (Best and Hummer, 2009; Lindorff-Larsen et al., 2010), ff99SB*-ILDN-Q (Best et al., 2012a; Best and Hummer, 2009; Lindorff-Larsen et al., 2010), a99SB-disp (Robustelli et al., 2018), RSFF2 (Li and Elcock, 2015; Zhou et al., 2015), CHARMM22* (Piana et al., 2011), CHARMM27 (Bjelkmar et al., 2010), CHARMM36 (Huang et al., 2017) and RSFF1 (Jiang et al., 2014).

We used as the solvent model the TIP3P adjusted for CHARMM force fields (MacKerell, et al., 1998), TIP3P (Jorgensen et al., 1983) for AMBER force fields and TIP4P-Ew (Horn et al., 2004) water model for the RSFF1 force field. We used a dodecahedral box with a minimum distance between protein and box edges of 12 Å applying periodic boundary conditions and a concentration of NaCl of 150mM, neutralizing the charges of the system. In the simulations, we used the Nε2-H tautomer for all the histidine residues. The system was equilibrated according to a protocol previously applied to other cases of study (Papaleo et al., 2014b). We carried out productive MD simulations in the canonical ensemble at 300 K and a time-step of 2 fs. We calculated long-range electrostatic interactions using the Particle-mesh Ewald summation scheme, whereas we truncated Van der Waals and Coulomb interactions at 10 Å.

### Structural assessment of the MD ensembles

We selected a subset of the MD ensembles, selecting 100 frames (equally spaced in time) from each of the ten different simulations as representative of each trajectory to be used in the prediction of chemical shifts and atomic resolution.

In particular, we calculated the backbone and proton side-chain chemical shift values for each MD ensemble with *PPM_One* (Li and Brüschweiler, 2015). We calculated the Root Mean Square Error (RMSE) and the Mean Average Error (MAE) between the predicted chemical shift values (from the simulations) and the experimentally measured chemical shifts from Biological Magnetic Resonance Bank (BMRB entry 5958) (Kouno et al., 2005) to assess the ability of the force fields in describing a conformational ensemble close to the experimental one. We used for the comparison 119 Cα, 116 C, 119 Hα, 113 H and 114 N chemical shifts, respectively.

We predicted the atomic resolutions with *ResProx* (Berjanskii et al., 2012) to assess the structural quality of the MD conformational ensembles representing each trajectory. *ResProx* uses machine learning techniques to estimate the atomic resolution of a protein structure from features that are derived from its atomic coordinates. We also verified that the *ResProx* results and their distribution were not affected by the approach used to select the subset of 100 MD frames. To this aim, we carried out the *ResProx* calculations on a set of equally-spaced 1000 frames selected from the one-μs CHARMM36 trajectory. Moreover, we also carried out structural clustering on the 1000-frame ensemble of CHARMM36 trajectory using the GROMOS algorithm for clustering. In the clustering, we used a mainchain Root Mean Square Deviation (RMSD) cutoff of 2.4 Å. We retained only the most populated cluster (which accounts for 888 structures) and estimated the atomic resolution of each structure of the cluster and the corresponding distribution. The two approaches gave results similar to calculations performed on the 100-frame ensembles, featuring similar distributions and median values.

To assess the statistical difference of the *ResProx* data for the different force field simulations, we calculated the distribution of the atomic resolution data for each MD ensemble and performed the Shapiro-Wilk test to evaluate if our data distribution can be approximated to a normal distribution. According to this test, most of the *ResProx* data distributions do not come from a normally-distributed population. In fact, the distribution of the sampled R values were either left- or right-skewed, due to outlier structures. The only exceptions are the data of the MD ensembles obtained with ff99SBnmr1 and a99SB-disp. We performed a pairwise Wilcoxon (Mann-Whitney U) rank sum test with continuity correction adjusting the p-value with the Holm-Bonferroni method on the R sets obtained from each force field pair to test whether the samples were selected from populations having the same distributions. The samples of R values from different force field simulation pairs that have p-values higher than 0.05 are not significantly different from each other under the Holm-Bonferroni correction.

### Principal Component Analysis of the MD trajectories

Principal component analysis (PCA) is used to extract the essential motions relevant to the function of the protein through the eigenvectors (principal components, PCs) of the covariance matrix of the positional fluctuations observed in MD trajectories, leaving out the irrelevant physically constrained local fluctuations of the protein (Amadei et al., 1993). We have performed all-atom PCA on a concatenated trajectory (including all the trajectories for the ten force fields) superposing the protein using the Cɑ coordinates to compare them in the same subspace. Prior the calculation, we discarded the six N-terminal and four C-terminal residues from our analyses, to prevent their motion to mask the important motion in the remainder of the protein. The first three PCs account for 35% of the fluctuations of the protein, which is not surprising since the trajectory used for the analyses is a concatenated one including all the force field simulations.

### MD ensemble comparison with ENCORE

We have used the *ENCORE* (Tiberti et al., 2015) as implemented in the *MDAnalysis* package (Michaud-Agrawal et al., 2011) to calculate ensemble similarity scores between each pair of ensembles. *ENCORE* estimates the probability distributions of the conformations that underlie each ensemble and calculates a probability similarity measures between each pair of them. To compare the LC3B simulations, we used the clustering ensemble similarity (CES) approach available in *ENCORE*. The method calculates the ensemble similarity as the Jensen-Shannon divergence between the estimated probability densities. CES partitions the whole conformational space in clusters and uses the relative populations of different ensembles in the clusters as an estimate of probability density. The CES values range between 0 and ln(2), where 0 indicates completely superimposable ensembles and ln(2) means non-overlapping ensembles. The clustering process is carried out using the affinity propagation method (Frey and Dueck, 2007) The calculation of the similarity score was carried out using 1000 frames for each simulation, on Cα only and excluding the flexible N- and C-terminal tails, as done in the PCA. The pairwise divergence values were visualised in heat maps and rendered as scatter plots using the tree preserving embedding method (Shieh et al., 2011).

### Structural alphabets

We have compared differences in the sampling of local conformations by analysing the trajectories using the M32K25 structural alphabet (SA for sake of clarity) (Pandini et al., 2010). This particular alphabet describes the local conformation of the protein by means of unique fragments made of Cαs of four consecutive residues, which were originally described by means of three angles. The 25 conformations or letters of the SA represent a set of canonical states describing the most probable local conformations (i.e. conformational attractors) in a set of experimentally derived protein structures. For every simulation, we have used *GSATools* (Pandini et al., 2013) to encode the conformation of each frame into a SA string, composed of 117 letters for our 120-residue protein. We then transformed the SA representation to that of a reduced structural alphabet (rSA), according to the mapping defined between these two alphabets (Pandini and Fornili, 2016). The rSA is a reduced representation of the original alphabet in which each letter corresponds to a macro-region of the density space. As we will be comparing distributions derived from the structural alphabet (see below), this ensures that the observed differences depend on significant differences between different states rather than the minor differences existing between letters of the original SA.

We then devised a fragment-wise measure to estimate the difference in sampling between different simulations. The following procedure has been carried out independently for each fragment. For each simulation, we calculated the frequency of each letter over the frames and used this as an estimate of the discrete probability distribution of the letters for that fragment and simulation. We then pairwise compared these distributions using the Jensen-Shannon divergence. In this way we obtained a JSd value for every pair of simulations, which accounts for the difference in sampling of different letters.

### Network Analyses of the MD Ensembles

We applied protein structure networks analysis (PSNs) to the MD ensemble as implemented in *Pyinteraph* (Tiberti et al., 2014). We defined as hubs those residues of the network with at least three edges, as commonly done for networks of protein structures (Papaleo, 2015). We used the node inter-connectivity to calculate the connected components, which are clusters of connected residues in the graph. For the contact-based PSN, we tested four different distance cutoffs to define the existence of a link between the nodes (i.e., 5, 5.125, 5.25 and 5.5 Å). Then we selected the cutoff of 5.125 Å as the best compromise between an entirely connected and a sparse network, according to our recent work on PSN cutoffs (Viloria et al., 2017). The distance was estimated between the center of mass of the residues side chains (expect glycines). Since MD force fields are known to have different mass definitions, we thus used the *PyInteraph* mass databases for each of the MD ensembles.

We also calculated other two PSNs to reflect other classes of intramolecular interactions, i.e., hydrogen bonds and salt bridges. For salt bridges, all of the distances between atom pairs belonging to charged moieties of two oppositely charged residues were calculated, and the charged moieties were considered as interacting if at least one pair of atoms was found at a distance shorter than 4.5 Å. In the case of aspartate and glutamate residues, the atoms forming the carboxylic group were considered. The NH3- and the guanidinium groups were employed for lysine and arginine, respectively. A hydrogen bond was identified when the distance between the acceptor atom and the hydrogen atom was lower than 3.5 Å and the donor-hydrogen-acceptor atom angle was greater than 120°.

To obtain contact, salt bridges or hydrogen bond-based PSNs for each MD ensemble, we retained only those edges which were present in at least 20% of the simulation frames (*pcrit* = 20%), as previously applied to other proteins (Papaleo et al., 2012b; Tiberti et al., 2014). We applied a variant of the depth-first search algorithm to identify the shortest path of communication. We defined the shortest path as the path in which the two residues were non-covalently connected by the smallest number of intermediate nodes. All the PSN calculations have been carried out using the *PyInteraph* suite of tools (Tiberti et al., 2014), whereas we used the *xPyder* plugin for *Pymol* (Pasi et al., 2012) the mapping of the connected components on the 3D structure.

### Contact analysis with CONAN

We performed the analysis of intramolecular contacts using the program *CONtact ANalysis* (*CONAN*) (Mercadante et al., 2018). *CONAN* is a powerful tool that implements the statistical and dynamical analysis of contacts in proteins and investigate how they evolve during time along MD simulations trajectories. The program is open-source and it performs the calculations of contacts during the trajectory times using the *mdmat* tool available in *GROMACS* version 5.1. We used the *CONAN* executable conan.py to perform the analysis over all the different force field simulations of LC3B. Inter-residue contacts in *CONAN* are defined by different cutoff of distances, that can be defined by the user.

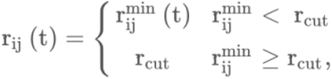

r_cut_ is the main cutoff and any pair of residues that doesn’t have at least an atom within this range are not considered. R_inter_ is the cutoff under which a contact is formed between a pair of residues. R_ij_^min^(t) is the minimum distance between atoms of residue i and j for a time period t. R_high-inter_ is the cutoff over which interactions are broken. We set up rcut at 10 Å, while r_inter_ and r_high-inter_ to 5 Å in agreement with the values used in the reference paper to study simulations of ubiquitin (Mercadante et al., 2018). We used a number of discretization levels to analyze contacts formation between 0 and r_cut_ of 1001, obtaining a resolution of 0.01 Å. We performed the calculation every one ns of simulations, considering all the atoms of the protein. We performed additional analysis on data on output from *CONAN* using the persistence value of each contact between pair of residues during simulation time, calculated as the frame in which the contact is identified divided by the total number of frames in the trajectory and the number of encounters, as the number that a contact is formed and broken during the trajectory. We used an in-house R script to produce the plot heatmaps in Figure 5D. We included in the supplementary materials the movie of contact map evolution during the trajectory created by *CONAN,* generated through mencoder, part of the *MPlayer* software package.

### Identification of cancer missense mutations and their annotation

We collected and aggregated a subset of cancer-related missense somatic mutations found in LC3B from *cBioPortal* (Cerami et al., 2012), considering all studies available on the 12th of October 2018 and *COSMIC* version 86 (Tate et al., 2019), considering all cancer types and excluding those mutations classified as natural polymorphisms. Moreover, we collected annotations on post-translational modification at the mutation sites using a local version of the *PhosphoSitePlus* database (Hornbeck et al., 2015), downloaded on 04/05/2018. Additional PTMs have been manually annotated through a survey of the literature on LC3B. We collected short linear motifs (SLiMs) located in proximity of the identified mutations using predictions from the Eukaryotic Linear Motif (*ELM*) server (Van Roey et al., 2013). Those SLiMs for which an interaction is not compatible with their localization on the LC3B structure have been discarded by further analyses, such as a PP2A docking site. Moreover, for each cancer samples where the mutation was identified we verified (when the information was available): i) the expression level of the LC3B gene; ii) if other mutations were occurring in the LC3B gene; and iii) if any of the interactors (see below) was mutated in the same sample. We also verified that any of the mutations was reported as possible natural polymorphism in the health population, through search in the *ExAC* database (Kobayashi et al., 2017). We predicted if each of the mutant variant could harbor new SLiMs querying *ELM* with the sequence of each mutant variant. We annotated each variant with the *REVEL* score (Nilah M. Ioannidis et al., 2016), as available on the *MyVariant.info* web resource (Xin et al., 2016). *REVEL* is an ensemble method for predicting the pathogenicity of missense variants from the scores generated by other individual prediction tools, which was found to be among the top performing pathogenicity predictors in a recent benchmarking effort (Li et al., 2018). The *REVEL* score can range from 0 to 1, with higher values indicating a stronger indication of pathogenicity. As done in the benchmarking study, we classified as pathogenic those variants having a score >= 0.4, which represents the best trade-off between sensitivity and specificity

### Coevolution analysis

We used two different parameters estimated by *Gremlin* (Ovchinnikov et al., 2014) to analyse the mutation sites. In particular, we employed: i) the conservation score estimated by the coupling matrix for the wild-type and the mutated residue at a certain position; ii) the residues that are coupled to the wild-type residue with a scaled score higher than one. We also derived a logo plot from the *Gremlin* sequence alignment with *Weblogo* (Crooks et al., 2004).

### LC3B interaction network and identification of LIR-containing candidates

We retrieved the known LC3B interactors through the Integrated Interaction Database (*IID*) version 2018-05 (Kotlyar et al., 2019). We retained only those interactions identified by at least two of the studies annotated in the database. We then predicted LIR motifs for each of the interactors through the *iLIR* database (Jacomin et al., 2016) and retained only those with a score higher than 11. This threshold was selected as it allows for a higher sensitivity (92%) at price of slightly lower specificity (Jacomin et al., 2016). We also verified through literature search if any of the interactors include one or more already experimentally verified LIR motifs (see Supplementary File 14). For each interactor with at least one LIR motif, we annotated the occurrence of cancer mutations in the same samples where a mutation of LC3B was found. We then retained only the interactors for which this mutation was abolishing a LIR motif or has mutations in its proximity. The resulting LC3B interactors were displayed as a network plot, using the *igraph* R package and in-house developed code.

### Structure-Based Prediction of Impact on Protein Stability and Binding Free Energies

We employed the *FoldX* energy function (Guerois et al., 2002; Schymkowitz et al., 2005) to perform *in silico* saturation mutagenesis using a *Python* wrapper that we recently developed. We used the same protocol that we recently applied to another protein (Nygaard et al., 2016). Calculations with the wrapper resulted in an average ΔΔG (differences in ΔG between mutant and wild-type variant) for each mutation over five independent runs performed using the NMR structure of LC3B (PDB entry 1V49, (Kouno et al., 2005)) and the X-ray crystallographic structure 3VTU (Rogov et al., 2013). We used the same pipeline to estimate the effect of mutations on the interaction free energy with the LIR domains using as a reference the structure of LC3B in complex with p62, FUNDC1 and ATG4B. This was performed by using the *AnalyseComplex* FoldX command on the mutant variant and corresponding wild-type conformation, and calculating the difference between their interaction energies.

Moreover, we also accounted for a correction to the *FoldX* energy values related to protein stability, as defined by Tawfik’s group (Tokuriki et al., 2007) to make the ΔΔG *FoldX* values more comparable with the expected experimental values, as previously described (Nygaard et al., 2016). In our calculations, we cannot use an experimental value of unfolding ΔG for the wild-type variant of LC3B domain since there are no experimental data available in the literature at the best of our knowledge. Nevertheless, values in the range of 5–15 kcal/mol are generally obtained for the net free energy of unfolding of proteins (Fersht and Serrano, 1993; Privalov, 1979). LC3B has a ubiquitin-like fold and we can refer to the free energy of unfolding experimentally measured for ubiquitin (Khorasanizadeh et al., 1996), which is 7.2 kcal/mol, which we used as reference value to estimate the ΔG of the mutant variant.

In parallel, we collected ΔΔG predictions upon mutation using the *Rosetta-*based *Flex ddG* protocol (Barlow et al., 2018), which couples standard side-chains repacking and minimization with a “backrub” approach to produce an ensemble of structures sampling backbone degrees of freedom and a generalized additive model applied to the *Rosetta* energy function. *Flex ddG* returns ΔΔG scores in *Rosetta* Energy Units (REUs), which are not directly convertible to kcal/mol but have been shown to correlate with experimental values (Barlow et al., 2018). We ran the protocol for each point mutation setting 35000 backrub trials, 5000 maximum iterations per minimization and an absolute score threshold for minimization convergence of 1.0 REUs. We generated ensembles of 35 different structures for each mutant and calculate the ΔΔGs for each structure and derived an average score. Destabilizing and stabilizing mutations are predicted using as empirical cutoffs values higher than 1 or lower than −1 REUs, respectively. We ran the protocol using Rosetta v2019.12.60667.

## Conclusions

We unveiled the effects exerted by missense mutations found in cancer genomic studies for the key autophagy ubiquitin-like protein, LC3B, providing a solid computational framework that allows to assess in parallel the impact on the most important properties that define its function and stability. We identified as driver sites for LC3B function four mutation sites that were already experimentally proved to alter the protein activity, supporting our approach. Moreover, we suggested new mutations for experimental studies, such as on D19, R70 and Y113C. For these variants, it would be useful to evaluate if the effect on the activity is dominated by a pronounced effect on protein stability which alters the turnover of LC3B mutated variants. R16G is also of interest, which has not been studied so far and it seems to play a critical role in modulating the protein stability. Moreover, our data, thanks to the collection of MD simulations with ten different force fields, can also guide the selection of physical models for MD simulations for the conformational ensemble and structure-dynamics-function relationships of the proteins of the Atg8 family, here illustrated on LC3B as a prototype of this family.

Our framework provides the groundwork to better understand the impact of mutations found in high-throughput cancer genomics data on a group of proteins that are key players of the autophagy pathway. More in general, it can be applied to the study of cancer proteins to prioritize candidates for experimental validation of their potential passenger and driver effects. In such cases it is also able to suggest the most convenient experimental methodologies for the validation, depending if the impact of the mutations is likely to be on the protein structural stability or its activity or even more specific aspects such as changes in post-translational regulation or allosteric mechanisms.

## Acknowledgments

This project was supported by an exploratory LEO foundation grant (LF17006), Carlsberg Foundation Distinguished Fellowship (CF18-0314), The Danish Council for Independent Research, Natural Science, Project 1 (102517), Danmarks Grundforskningsfond (DNRF125) to EP group. Moreover, the project has been supported by a KBVU pre-graduate fellowship and a Netaji Subhash ICAR international fellowship, Govt.of India to MaK and MuK to work in EP group, respectively. The calculations described in this paper were performed using the DeiC National Life Science Supercomputer Computerome at DTU (Denmark), a DECI-PRACE 14th HPC Grant for calculations on Archer (UK), and ISCRA-CINECA HP10C0T58M. The authors would like to thank Lisa Cantwell for the professional scientific proofreading of the manuscript and Emiliano Maiani for fruitful comments and discussion.

## References

Alemu EA, Lamark T, Torgersen KM, Birgisdottir AB, Larsen KB, Jain A, Olsvik H, Øvervatn A, Kirkin V, Johansen T. 2012. ATG8 family proteins act as scaffolds for assembly of the ULK complex: Sequence requirements for LC3-interacting region (LIR) motifs. J Biol Chem 287:39275–39290. doi:10.1074/jbc.M112.378109

Amadei A, Linssen AB, Berendsen HJ. 1993. Essential dynamics of proteins. Proteins 17:412–25. doi:10.1002/prot.340170408

Angelova K, Felline A, Lee M, Patel M, Puett D, Fanelli F. 2011. Conserved amino acids participate in the structure networks deputed to intramolecular communication in the lutropin receptor. Cell Mol Life Sci 68:1227–39. doi:10.1007/s00018-010-0519-z

Barlow KA, Ó Conchúir S, Thompson S, Suresh P, Lucas JE, Heinonen M, Kortemme T. 2018. Flex ddG: Rosetta Ensemble-Based Estimation of Changes in Protein-Protein Binding Affinity upon Mutation. J Phys Chem B 122:5389–5399. doi:10.1021/acs.jpcb.7b11367

Beauchamp KA, Lin Y, Das R, Pande VS. 2012. Are Protein Force Fields Getting Better ? A Systematic Benchmark on 524 Diverse NMR Measurements.

Behrends C, Sowa ME, Gygi SP, Harper JW. 2010. Network organization of the human autophagy system. Nature 466:68–76. doi:10.1038/nature09204

Berjanskii M, Zhou J, Liang Y, Lin G, Wishart DS. 2012. Resolution-by-proxy: a simple measure for assessing and comparing the overall quality of NMR protein structures. J Biomol NMR 53:167– 80. doi:10.1007/s10858-012-9637-2

Best RB, De Sancho D, Mittal J. 2012a. Residue-specific α-helix propensities from molecular simulation. Biophys J 102:1462–1467. doi:10.1016/j.bpj.2012.02.024

Best RB, Hummer G. 2009. Optimized molecular dynamics force fields applied to the helix-coil transition of polypeptides. J Phys Chem B 113:9004–15. doi:10.1021/jp901540t

Best RB, Zhu X, Shim J, Lopes PEM, Mittal J, Feig M, Alexander D. MacKerell J. 2012b. Optimization of the Additive CHARMM All-Atom Protein Force Field Targeting Improved Sampling of the Backbone ϕ, ψ and Side-Chain χ1 and χ2 Dihedral Angles. J Chem Theory Comput 8:3257– 3273.

Birgisdottir AB, Lamark T, Johansen T. 2013. The LIR motif - crucial for selective autophagy. J Cell Sci 126:3552–62. doi:10.1242/jcs.120477

Bjelkmar P, Larsson P, Cuendet MA, Hess B, Lindahl E. 2010. Implementation of the CHARMM Force Field in GROMACS: Analysis of Protein Stability Effects from Correction Maps, Virtual Interaction Sites, and Water Models. J Chem Theory Comput 6:459–66. doi:10.1021/ct900549r

Blom N, Sicheritz-Pontén T, Gupta R, Gammeltoft S, Brunak S. 2004. Prediction of post-translational glycosylation and phosphorylation of proteins from the amino acid sequence. Proteomics 4:1633–1649. doi:10.1002/pmic.200300771

Bowman GR. 2016. Accurately modeling nanosecond protein dynamics requires at least microseconds of simulation. J Comput Chem 37:558–566. doi:10.1002/jcc.23973

Buß O, Rudat J, Ochsenreither K. 2018. FoldX as Protein Engineering Tool: Better Than Random Based Approaches? Comput Struct Biotechnol J 16:25–33. doi:10.1016/j.csbj.2018.01.002

Case D a. 2013. Chemical shifts in biomolecules. Curr Opin Struct Biol 23:172–179. doi:10.1016/j.sbi.2013.01.007

Cerami E, Gao J, Dogrusoz U, Gross BE, Sumer SO, Aksoy BA, Jacobsen A, Byrne CJ, Heuer ML, Larsson E, Antipin Y, Reva B, Goldberg AP, Sander C, Schultz N. 2012. The cBio Cancer Genomics Portal: An open platform for exploring multidimensional cancer genomics data. Cancer Discov 2:401–404. doi:10.1158/2159-8290.CD-12-0095

Costa JR, Prak K, Aldous S, Gewinner CA, Ketteler R. 2016. Autophagy gene expression profiling identifies a defective microtubule-associated protein light chain 3A mutant in cancer. Oncotarget 7. doi:10.18632/oncotarget.9754

Cowey CL, Rathmell WK. 2009. VHL gene mutations in renal cell carcinoma: role as a biomarker of disease outcome and drug efficacy. Curr Oncol Rep 11:94–101.

Craveur P, Joseph AP, Esque J, Narwani TJ, NoÃ«l F, Shinada N, Goguet M, Leonard S, Poulain P, Bertrand O, Faure G, Rebehmed J, Ghozlane A, Swapna LS, Bhaskara RM, Barnoud J, TÃ©letchÃ©a S, Jallu V, Cerny J, Schneider B, Etchebest C, Srinivasan N, Gelly J-C, de Brevern AG. 2015. Protein flexibility in the light of structural alphabets. Front Mol Biosci 2:1–20. doi:10.3389/fmolb.2015.00020

Crooks G, Hon G, Chandonia J, Brenner S. 2004. WebLogo: a sequence logo generator. Genome Res 14:1188–1190. doi:10.1101/gr.849004.1

Di Paola L, Giuliani A. 2015. Protein contact network topology: a natural language for allostery. Curr Opin Struct Biol 31:43–48. doi:10.1016/j.sbi.2015.03.001

Di Rita A, Peschiaroli A, D′Acunzo P, Strobbe D, Hu Z, Gruber J, Nygaard M, Lambrughi M, Melino G, Papaleo E, Dengjel J, El Alaoui S, Campanella M, Dötsch V, Rogov V V., Strappazzon F, Cecconi F. 2018. HUWE1 E3 ligase promotes PINK1/PARKIN-independent mitophagy by regulating AMBRA1 activation via IKKα. Nat Commun 9:3755. doi:10.1038/s41467-018-05722-3

Dou Z, Xu C, Donahue G, Shimi T, Pan JA, Zhu J, Ivanov A, Capell BC, Drake AM, Shah PP, Catanzaro JM, Ricketts MD, Lamark T, Adam SA, Marmorstein R, Zong WX, Johansen T, Goldman RD, Adams PD, Berger SL. 2015. Autophagy mediates degradation of nuclear lamina. Nature 527:105–109. doi:10.1038/nature15548

Dror RO, Dirks RM, Grossman JP, Xu H, Shaw DE. 2012. Biomolecular simulation: a computational microscope for molecular biology. Annu Rev Biophys 41:429–52. doi:10.1146/annurev-biophys-042910-155245

F Jelc, Diagne F, Fiorello M, Janani M, Kargbo P, Kilby A, León G, Polley T, Knight B, Laitin D, Leamer E, Munshi K, Olken B, Rao B, Ray D. 2012. Beyond the histone take: HP1alpha deregulation in breast cancer epigenetics. doi:10.1080/02626667.2015.1006226

Fass E, Amar N, Elazar Z. 2007. Identification of essential residues for the C-terminal cleavage of the mammalian LC3: a lesson from yeast Atg8. Autophagy 3:48–50.

Fersht AR, Serrano L. 1993. Principles of protein stability derived from protein engineering experiments. Curr Opin Struct Biol 3:75–83. doi:10.1016/0959-440X(93)90205-Y

Fimia GM, Kroemer G, Piacentini M. 2013. Molecular mechanisms of selective autophagy. Cell Death Differ 20:1–2. doi:10.1038/cdd.2012.97

Frey BJ, Dueck D. 2007. Clustering by passing messages between data points. Science (80-) 1–15. doi:10.1126/science.1136800

Galluzzi L, Baehrecke EH, Ballabio A, Boya P, Bravo San Pedro JM, Cecconi F, Choi AM, Chu CT, Codogno P, Colombo MI, Cuervo AM, Debnath J, Deretic V, Dikic I, Eskelinen E, Fimia GM, Fulda S, Gewirtz DA, Green DR, Hansen M, Harper JW, Jäättelä M, Johansen T, Juhasz G, Kimmelman AC, Kraft C, Ktistakis NT, Kumar S, Levine B, Lopez-Otin C, Madeo F, Martens S, Martinez J, Melendez A, Mizushima N, Münz C, Murphy LO, Penninger JM, Piacentini M, Reggiori F, Rubinsztein DC, Ryan KM, Santambrogio L, Scorrano L, Simon AK, Simon H, Simonsen A, Tavernarakis N, Tooze SA, Yoshimori T, Yuan J, Yue Z, Zhong Q, Kroemer G. 2017. Molecular definitions of autophagy and related processes. EMBO J 36:1811–1836. doi:10.15252/embj.201796697

Gatica D, Lahiri V, Klionsky DJ. 2018. Cargo recognition and degradation by selective autophagy. Nat Cell Biol 20:233–242. doi:10.1038/s41556-018-0037-z

Ghosh A, Vishveshwara S. 2008. Variations in clique and community patterns in protein structures during allosteric communication: Investigation of dynamically equilibrated structures of methionyl tRNA synthetase complexes. Biochemistry 47:11398–11407. doi:10.1021/bi8007559

Guerois R, Nielsen JE, Serrano L. 2002. Predicting changes in the stability of proteins and protein complexes: A study of more than 1000 mutations. J Mol Biol 320:369–387. doi:10.1016/S0022-2836(02)00442-4

Guvench O, MacKerell AD. 2008. Comparison of Protein Force Fields for Molecular Dynamics Simulations. pp. 63–88. doi:10.1007/978-1-59745-177-2_4

He C, Klionsky DJ. 2009. Regulation Mechanisms and Signaling Pathways of Autophagy. Annu Rev Genet 43:67–93. doi:10.1093/nq/s2-V.108.66

He Y, Zhao X, Subahan NR, Fan L, Gao J, Chen H. 2014. The prognostic value of autophagy-related markers beclin-1 and microtubule-associated protein light chain 3B in cancers: a systematic review and meta-analysis. Tumour Biol 35:7317–26. doi:10.1007/s13277-014-2060-4

Henriques J, Cragnell C, Skepö M. 2015. Molecular dynamics simulations of intrinsically disordered proteins: force field evaluation and comparison with experiment. J Chem Theory Comput 11:3420–3431. doi:10.1021/ct501178z

Hess B, Kutzner C, van der Spoel D, Lindahl E. 2008. GROMACS 4: Algorithms for Highly Efficient, Load-Balanced, and Scalable Molecular Simulation. J Chem Theory Comput 4:435–447. doi:10.1021/ct700301q

Horn HW, Swope WC, Pitera JW, Madura JD, Dick TJ, Hura GL, Head-Gordon T. 2004. Development of an improved four-site water model for biomolecular simulations: TIP4P-Ew. J Chem Phys 120:9665–78. doi:10.1063/1.1683075

Hornbeck P V., Zhang B, Murray B, Kornhauser JM, Latham V, Skrzypek E. 2015. PhosphoSitePlus, 2014: mutations, PTMs and recalibrations. Nucleic Acids Res 43:D512–D520. doi:10.1093/nar/gku1267

Huang J, Rauscher S, Nawrocki G, Ran T, Feig M, de Groot BL, Grubmüller H, MacKerell AD. 2017. CHARMM36m: an improved force field for folded and intrinsically disordered proteins. Nat Methods 14:71–73. doi:10.1038/nmeth.4067

Huang R, Liu W. 2015. Identifying an essential role of nuclear LC3 for autophagy. Autophagy 11:852–853. doi:10.1080/15548627.2015.1038016

Huang R, Xu Y, Lippincott-schwartz J, Liu W, Huang R, Xu Y, Wan W, Shou X, Qian J, You Z, Liu B, Chang C, Zhou T. 2015. Deacetylation of Nuclear LC3 Drives Autophagy Initiation under Starvation Article Deacetylation of Nuclear LC3 Drives Autophagy Initiation under Starvation. Mol Cell 57:456–466. doi:10.1016/j.molcel.2014.12.013

Ichimura Y, Kumanomidou T, Sou YS, Mizushima T, Ezaki J, Ueno T, Kominami E, Yamane T, Tanaka K, Komatsu M. 2008. Structural basis for sorting mechanism of p62 in selective autophagy. J Biol Chem 283:22847–22857. doi:10.1074/jbc.M802182200

Invernizzi G, Tiberti M, Lambrughi M, Lindorff-Larsen K, Papaleo E. 2014. Communication Routes in ARID Domains between Distal Residues in Helix 5 and the DNA-Binding Loops. PLoS Comput Biol 10:e1003744. doi:10.1371/journal.pcbi.1003744

Ioannidis NM, Rothstein JH, Pejaver V, Middha S, McDonnell SK, Baheti S, Musolf A, Li Q, Holzinger E, Karyadi D, Cannon-Albright LA, Teerlink CC, Stanford JL, Isaacs WB, Xu J, Cooney KA, Lange EM, Schleutker J, Carpten JD, Powell IJ, Cussenot O, Cancel-Tassin G, Giles GG, MacInnis RJ, Maier C, Hsieh C-L, Wiklund F, Catalona WJ, Foulkes WD, Mandal D, Eeles RA, Kote-Jarai Z, Bustamante CD, Schaid DJ, Hastie T, Ostrander EA, Bailey-Wilson JE, Radivojac P, Thibodeau SN, Whittemore AS, Sieh W. 2016. REVEL: An Ensemble Method for Predicting the Pathogenicity of Rare Missense Variants. Am J Hum Genet 99:877–885. doi:10.1016/j.ajhg.2016.08.016

Jacomin AC, Samavedam S, Promponas V, Nezis IP. 2016. iLIR database: A web resource for LIR motif-containing proteins in eukaryotes. Autophagy 12:1–9. doi:10.1080/15548627.2016.1207016

Jatana N, Ascher DB, Pires DEV, Gokhale RS, Thukral L. 2019. Human LC3 and GABARAP subfamily members achieve functional specificity via specific structural modulations. Autophagy 0:15548627.2019.1606636. doi:10.1080/15548627.2019.1606636

Jemal A, Bray F, Center MM, Ferlay J, Ward E, Forman D. 2011. Global Cancer Statistics: 2011. CA Cancer J Clin 61:69–90. doi:10.3322/caac.20107.Available

Jiang F, Zhou C-Y, Wu Y-D. 2014. Residue-specific force field based on the protein coil library. RSFF1: modification of OPLS-AA/L. J Phys Chem B 118:6983–98. doi:10.1021/jp5017449

Jiang H, Cheng D, Liu W, Peng J, Feng J. 2010. Protein kinase C inhibits autophagy and phosphorylates LC3. Biochem Biophys Res Commun 395:471–476. doi:10.1016/j.bbrc.2010.04.030

Johansen T, Lamark T. 2011. Selective autophagy mediated by autophagic adapter proteins. Autophagy 7:279–296. doi:10.4161/auto.7.3.14487

Jónsdóttir LB, Ellertsson B, Invernizzi G, Magnúsdóttir M, Thorbjarnardóttir SH, Papaleo E, Kristjánsson MM. 2014. The role of salt bridges on the temperature adaptation of aqualysin I, a thermostable subtilisin-like proteinase. Biochim Biophys Acta - Proteins Proteomics 1844:2174–2181. doi:10.1016/j.bbapap.2014.08.011

Jorgensen WL, Chandrasekhar J, Madura JD, Impey RW, Klein ML. 1983. Comparison of simple potential functions for simulating liquid water. J Chem Phys 79:926. doi:10.1063/1.445869

Kabeya Y. 2004. LC3, GABARAP and GATE16 localize to autophagosomal membrane depending on form-II formation. J Cell Sci 117:2805–2812. doi:10.1242/jcs.01131

Kabeya Y, Mizushima N, Ueno T, Yamamoto A, Kirisako T, Noda T, Kominami E, Ohsumi Y, Yoshimori T. 2000. LC3, a mammalian homologue of yeast Apg8p, is localized in autophagosome membranes after processing. EMBO J 19:5720–5728. doi:10.1093/emboj/19.21.5720

Khorasanizadeh S, Peters ID, Roder H. 1996. Evidence for a three-state model of protein folding from kinetic analysis of ubiquitin variants with altered core residues. Nat Struct Biol 3:193–205.

Kim BW, Kwon DH, Song HK. 2016. Structure biology of selective autophagy receptors. BMB Rep 49:73–80. doi:10.5483/BMBRep.2016.49.2.265

Kimmelman AC. 2011. The dynamic nature of autophagy in cancer. Genes Dev 25:1999–2010. doi:10.1101/gad.17558811

Kirkin V, Lamark T, Sou YS, Bjørkøy G, Nunn JL, Bruun JA, Shvets E, McEwan DG, Clausen TH, Wild P, Bilusic I, Theurillat JP, Øvervatn A, Ishii T, Elazar Z, Komatsu M, Dikic I, Johansen T. 2009. A Role for NBR1 in Autophagosomal Degradation of Ubiquitinated Substrates. Mol Cell 33:505–516. doi:10.1016/j.molcel.2009.01.020

Klepeis JL, Lindorff-Larsen K, Dror RO, Shaw DE. 2009. Long-timescale molecular dynamics simulations of protein structure and function. Curr Opin Struct Biol 19:120–7. doi:10.1016/j.sbi.2009.03.004

Klionsky DJ, Abdelmohsen K, Abe A, Abedin MJ, Abeliovich H, Arozena AA, Adachi H, Adams CM, Adams PD, Adeli K, Adhihetty PJ, Adler SG, Agam G, Agarwal R, Aghi MK, Agnello M, Agostinis P, Aguilar P V., Aguirre-Ghiso J, Airoldi EM, Ait-Si-Ali S, Akematsu T, Akporiaye ET, Al-Rubeai M, Albaiceta GM, Albanese C, Albani D, Albert ML, Aldudo J, Algül H, Alirezaei M, Alloza I, Almasan A, Almonte-Beceril M, Alnemri ES, Alonso C, Altan-Bonnet N, Altieri DC, Alvarez S, Alvarez-Erviti L, Alves S, Amadoro G, Amano A, Amantini C, Ambrosio S, Amelio I, Amer AO, Amessou M, Amon A, An Z, Anania FA, Andersen SU, Andley UP, Andreadi CK, Andrieu-Abadie N, Anel A, Ann DK, Anoopkumar-Dukie S, Antonioli M, Aoki H, Apostolova N, Aquila S, Aquilano K, Araki K, Arama E, Aranda A, Araya J, Arcaro A, Arias E, Arimoto H, Ariosa AR, Armstrong JL, Arnould T, Arsov I, Asanuma K, Askanas V, Asselin E, Atarashi R, Atherton SS, Atkin JD, Attardi LD, Auberger P, Auburger G, Aurelian L, Autelli R, Avagliano L, Avantaggiati ML, Avrahami L, Azad N, Awale S, Bachetti T, Backer JM, Bae DH, Bae JS, Bae ON, Bae SH, Baehrecke EH, Baek SH, Baghdiguian S, Bagniewska-Zadworna A, Bai H, Bai J, Bai XY, Bailly Y, Balaji KN, Balduini W, Ballabio A, Balzan R, Banerjee R, Bánhegyi G, Bao H, Barbeau B, Barrachina MD, Barreiro E, Bartel B, Bartolomé A, Bassham DC, Bassi MT, Bast RC, Basu A, Batista MT, Batoko H, Battino M, Bauckman K, Baumgarner BL, Bayer KU, Beale R, Beaulieu JF, Beck GR, Becker C, Beckham JD, Bédard PA, Bednarski PJ, Begley TJ, Behl C, Behrends C, Behrens GMN, Behrns KE, Bejarano E, Belaid A, Belleudi F, Bénard G, Berchem G, Bergamaschi D, Bergami M, Berkhout B, Berliocchi L, Bernard A, Bernard M, Bernassola F, Bertolotti A, Bess AS, Besteiro S, Bettuzzi S, Bhalla S, Bhattacharyya S, Bhutia SK, Biagosch C, Bianchi MW, Biard-Piechaczyk M, Billes V, Bincoletto C, Bingol B, Bird SW, Bitoun M, Bjedov I, Blackstone C, Blanc L, Blanco GA, Blomhoff HK, Boada-Romero E, Böckler S, Boes M, Boesze-Battaglia K, Boise LH, Bolino A, Boman A, Bonaldo P, Bordi M, Bosch J, Botana LM, Botti J, Bou G, Bouché M, Bouchecareilh M, Boucher MJ, Boulton ME, Bouret SG, Boya P, Boyer-Guittaut M, Bozhkov P V., Brady N, Braga VMM, Brancolini C, Braus GH, Bravosan-Pedro JM, Brennan LA, Bresnick EH, Brest P, Bridges D, Bringer MA, Brini M, Brito GC, Brodin B, Brookes PS, Brown EJ, Brown K, Broxmeyer HE, Bruhat A, Brum PC, Brumell JH, Brunetti-Pierri N, Bryson-Richardson RJ, Buch S, Buchan AM, Budak H, Bulavin D V., Bultman SJ, Bultynck G, Bumbasirevic V, Burelle Y, Burke RE, Burmeister M, Bütikofer P, Caberlotto L, Cadwell K, Cahova M, Cai D, Cai J, Cai Q, Calatayud S, Camougrand N, Campanella M, Campbell GR, Campbell M, Campello S, Candau R, Caniggia I, Cantoni L, Cao L, Caplan AB, Caraglia M, Cardinali C, Cardoso SM, Carew JS, Carleton LA, Carlin CR, Carloni S, Carlsson SR, Carmona-Gutierrez D, Carneiro LAM, Carnevali O, Carra S, Carrier A, Carroll B, Casas C, Casas J, Cassinelli G, Castets P, Castro-Obregon S, Cavallini G, Ceccherini I, Cecconi F, Cederbaum AI, Ceña V, Cenci S, Cerella C, Cervia D, Cetrullo S, Chaachouay H, Chae HJ, Chagin AS, Chai CY, Chakrabarti G, Chamilos G, Chan EYW, Chan MTV, Chandra D, Chandra P, Chang CP, Chang RCC, Chang TY, Chatham JC, Chatterjee S, Chauhan S, Che Y, Cheetham ME, Cheluvappa R, Chen CJ, Chen G, Chen GC, Chen G, Chen H, Chen JW, Chen JK, Chen M, Chen M, Chen P, Chen Q, Chen Q, Chen S Der, Chen S, Chen SSL, Chen W, Chen WJ, Chen WQ, Chen W, Chen X, Chen YH, Chen YG, Chen Y, Chen Y, Chen Y, Chen YJ, Chen YQ, Chen Y, Chen Z, Chen Z, Cheng A, Cheng CHK, Cheng H, Cheong H, Cherry S, Chesney J, Cheung CHA, Chevet E, Chi HC, Chi SG, Chiacchiera F, Chiang HL, Chiarelli R, Chiariello M, Chieppa M, Chin LS, Chiong M, Chiu GNC, Cho DH, Cho SG, Cho WC, Cho YY, Cho YS, Choi AMK, Choi EJ, Choi EK, Choi J, Choi ME, Choi S Il, Chou TF, Chouaib S, Choubey D, Choubey V, Chow KC, Chowdhury K, Chu CT, Chuang TH, Chun T, Chung H, Chung T, Chung YL, Chwae YJ, Cianfanelli V, Ciarcia R, Ciechomska IA, Ciriolo MR, Cirone M, Claerhout S, Clague MJ, Clària J, Clarke PGH, Clarke R, Clementi E, Cleyrat C, Cnop M, Coccia EM, Cocco T, Codogno P, Coers J, Cohen EEW, Colecchia D, Coletto L, Coll NS, Colucci-Guyon E, Comincini S, Condello M, Cook KL, Coombs GH, Cooper CD, Cooper JM, Coppens I, Corasaniti MT, Corazzari M, Corbalan R, Corcelle-Termeau E, Cordero MD, Corral-Ramos C, Corti O, Cossarizza A, Costelli P, Costes S, Cotman SL, Coto-Montes A, Cottet S, Couve E, Covey LR, Cowart LA, Cox JS, Coxon FP, Coyne CB, Cragg MS, Craven RJ, Crepaldi T, Crespo JL, Criollo A, Crippa V, Cruz MT, Cuervo AM, Cuezva JM, Cui T, Cutillas PR, Czaja MJ, Czyzyk-Krzeska MF, Dagda RK, Dahmen U, Dai C, Dai W, Dai Y, Dalby KN, Valle LD, Dalmasso G, D’amelio M, Damme M, Darfeuille-Michaud A, Dargemont C, Darley-Usmar VM, Dasarathy S, Dasgupta B, Dash S, Dass CR, Davey HM, Davids LM, Dávila D, Davis RJ, Dawson TM, Dawson VL, Daza P, de Belleroche J, de Figueiredo P, de Figueiredo RCBQ, de la Fuente J, De Martino L, De Matteis A, De Meyer GRY, De Milito A, De Santi M, de Souza W, De Tata V, De Zio D, Debnath J, Dechant R, Decuypere JP, Deegan S, Dehay B, Del Bello B, Del Re DP, Delage-Mourroux R, Delbridge LMD, Deldicque L, Delorme-Axford E, Deng Y, Dengjel J, Denizot M, Dent P, Der CJ, Deretic V, Derrien B, Deutsch E, Devarenne TP, Devenish RJ, Di Bartolomeo S, Di Daniele N, Di Domenico F, Di Nardo A, Di Paola S, Di Pietro A, Di Renzo L, Di Antonio A, Díaz-Araya G, Díaz-Laviada I, Diaz-Meco MT, Diaz-Nido J, Dickey CA, Dickson RC, Diederich M, Digard P, Dikic I, Dinesh-Kumar SP, Ding C, Ding WX, Ding Z, Dini L, Distler JHW, Diwan A, Djavaheri-Mergny M, Dmytruk K, Dobson RCJ, Doetsch V, Dokladny K, Dokudovskaya S, Donadelli M, Dong XC, Dong X, Dong Z, Donohue TM, Donohue-Jr TM, Doran KS, D’orazi G, Dorn GW, Dosenko V, Dridi S, Drucker L, Du J, Du LL, Du L, du Toit A, Dua P, Duan L, Duann P, Dubey VK, Duchen MR, Duchosal MA, Duez H, Dugail I, Dumit VI, Duncan MC, Dunlop EA, Dunn WA, Dupont N, Dupuis L, Durán R V., Durcan TM, Duvezin-Caubet S, Duvvuri U, Eapen V, Ebrahimi-Fakhari D, Echard A, Eckhart L, Edelstein CL, Edinger AL, Eichinger L, Eisenberg T, Eisenberg-Lerner A, Eissa NT, El-Deiry WS, El-Khoury V, Elazar Z, Eldar-Finkelman H, Elliott CJH, Emanuele E, Emmenegger U, Engedal N, Engelbrecht AM, Engelender S, Enserink JM, Erdmann R, Erenpreisa J, Eri R, Eriksen JL, Erman A, Escalante R, Eskelinen EL, Espert L, Esteban-Martínez L, Evans TJ, Fabri M, Fabrias G, Fabrizi C, Facchiano A, Færgeman NJ, Faggioni A, Fairlie WD, Fan C, Fan D, Fan J, Fang S, Fanto M, Fanzani A, Farkas T, Faure M, Favier FB, Fearnhead H, Federici M, Fei E, Felizardo TC, Feng H, Feng Y, Feng Y, Ferguson TA, Fernández ÁF, Fernandez-Barrena MG, Fernandez-Checa JC, Fernández-López A, Fernandez-Zapico ME, Feron O, Ferraro E, Ferreira-Halder CV, Fesus L, Feuer R, Fiesel FC, Filippi-Chiela EC, Filomeni G, Fimia GM, Fingert JH, Finkbeiner S, Finkel T, Fiorito F, Fisher PB, Flajolet M, Flamigni F, Florey O, Florio S, Floto RA, Folini M, Follo C, Fon EA, Fornai F, Fortunato F, Fraldi A, Franco R, Francois A, François A, Frankel LB, Fraser IDC, Frey N, Freyssenet DG, Frezza C, Friedman SL, Frigo DE, Fu D, Fuentes JM, Fueyo J, Fujitani Y, Fujiwara Y, Fujiya M, Fukuda M, Fulda S, Fusco C, Gabryel B, Gaestel M, Gailly P, Gajewska M, Galadari S, Galili G, Galindo I, Galindo MF, Galliciotti G, Galluzzi L, Galluzzi L, Galy V, Gammoh N, Gandy S, Ganesan AK, Ganesan S, Ganley IG, Gannagé M, Gao FB, Gao F, Gao JX, Nannig LG, Véscovi EG, Garcia-Macía M, Garcia-Ruiz C, Garg AD, Garg PK, Gargini R, Gassen NC, Gatica D, Gatti E, Gavard J, Gavathiotis E, Ge L, Ge P, Ge S, Gean PW, Gelmetti V, Genazzani AA, Geng J, Genschik P, Gerner L, Gestwicki JE, Gewirtz DA, Ghavami S, Ghigo E, Ghosh D, Giammarioli AM, Giampieri F, Giampietri C, Giatromanolaki A, Gibbings DJ, Gibellini L, Gibson SB, Ginet V, Giordano A, Giorgini F, Giovannetti E, Girardin SE, Gispert S, Giuliano S, Gladson CL, Glavic A, Gleave M, Godefroy N, Gogal RM, Gokulan K, Goldman GH, Goletti D, Goligorsky MS, Gomes A V., Gomes LC, Gomez H, Gomez-Manzano C, Gómez-Sánchez R, Gonçalves DAP, Goncu E, Gong Q, Gongora C, Gonzalez CB, Gonzalez-Alegre P, Gonzalez-Cabo P, González-Polo RA, Goping IS, Gorbea C, Gorbunov N V., Goring DR, Gorman AM, Gorski SM, Goruppi S, Goto-Yamada S, Gotor C, Gottlieb RA, Gozes I, Gozuacik D, Graba Y, Graef M, Granato GE, Grant GD, Grant S, Gravina GL, Green DR, Greenhough A, Greenwood MT, Grimaldi B, Gros F, Grose C, Groulx JF, Gruber F, Grumati P, Grune T, Guan JL, Guan KL, Guerra B, Guillen C, Gulshan K, Gunst J, Guo C, Guo L, Guo M, Guo W, Guo XG, Gust AA, Gustafsson ÅB, Gutierrez E, Gutierrez MG, Gwak HS, Haas A, Haber JE, Hadano S, Hagedorn M, Hahn DR, Halayko AJ, Hamacher-Brady A, Hamada K, Hamai A, Hamann A, Hamasaki M, Hamer I, Hamid Q, Hammond EM, Han F, Han W, Handa JT, Hanover JA, Hansen M, Harada M, Harhaji-Trajkovic L, Harper JW, Harrath AH, Harris AL, Harris J, Hasler U, Hasselblatt P, Hasui K, Hawley RG, Hawley TS, He C, He CY, He F, He G, He RR, He XH, He YW, He YY, Heath JK, Hébert MJ, Heinzen RA, Helgason GV, Hensel M, Henske EP, Her C, Herman PK, Hernández A, Hernandez C, Hernández-Tiedra S, Hetz C, Hiesinger PR, Higaki K, Hilfiker S, Hill BG, Hill JA, Hill WD, Hino K, Hofius D, Hofman P, Höglinger GU, Höhfeld J, Holz MK, Hong Y, Hood DA, Hoozemans JJM, Hoppe T, Hsu C, Hsu CY, Hsu LC, Hu D, Hu G, Hu HM, Hu H, Hu MC, Hu YC, Hu ZW, Hua F, Hua Y, Huang C, Huang HL, Huang KH, Huang KY, Huang S, Huang S, Huang WP, Huang YR, Huang Y, Huang Y, Huber TB, Huebbe P, Huh WK, Hulmi JJ, Hur GM, Hurley JH, Husak Z, Hussain SNA, Hussain S, Hwang JJ, Hwang S, Hwang TIS, Ichihara A, Imai Y, Imbriano C, Inomata M, Into T, Iovane V, Iovanna JL, Iozzo R V., Ip NY, Irazoqui JE, Iribarren P, Isaka Y, Isakovic AJ, Ischiropoulos H, Isenberg JS, Ishaq M, Ishida H, Ishii I, Ishmael JE, Isidoro C, Isobe KI, Isono E, Issazadeh-Navikas S, Itahana K, Itakura E, Ivanov AI, Iyer AK V., Izquierdo JM, Izumi Y, Izzo V, Jäättelä M, Jaber N, Jackson DJ, Jackson WT, Jacob TG, Jacques TS, Jagannath C, Jain A, Jana NR, Jang BK, Jani A, Janji B, Jannig PR, Jansson PJ, Jean S, Jendrach M, Jeon JH, Jessen N, Jeung EB, Jia K, Jia L, Jiang H, Jiang H, Jiang L, Jiang T, Jiang X, Jiang X, Jiang Y, Jiang Y, Jiménez A, Jin C, Jin H, Jin L, Jin M, Jin S, Jinwal UK, Jo EK, Johansen T, Johnson DE, Johnson GVW, Johnson JD, Jonasch E, Jones C, Joosten LAB, Jordan J, Joseph AM, Joseph B, Joubert AM, Ju D, Ju J, Juan HF, Juenemann K, Juhász G, Jung HS, Jung JU, Jung YK, Jungbluth H, Justice MJ, Jutten B, Kaakoush NO, Kaarniranta K, Kaasik A, Kabuta T, Kaeffer B, Kågedal K, Kahana A, Kajimura S, Kakhlon O, Kalia M, Kalvakolanu D V., Kamada Y, Kambas K, Kaminskyy VO, Kampinga HH, Kandouz M, Kang C, Kang R, Kang TC, Kanki T, Kanneganti TD, Kanno H, Kanthasamy AG, Kantorow M, Kaparakis-Liaskos M, Kapuy O, Karantza V, Karim MR, Karmakar P, Kaser A, Kaushik S, Kawula T, Kaynar AM, Ke PY, Ke ZJ, Kehrl JH, Keller KE, Kemper JK, Kenworthy AK, Kepp O, Kern A, Kesari S, Kessel D, Ketteler R, Kettelhut I do C, Khambu B, Khan MM, Khandelwal VKM, Khare S, Kiang JG, Kiger AA, Kihara A, Kim AL, Kim CH, Kim DR, Kim DH, Kim EK, Kim HY, Kim HR, Kim JS, Kim JH, Kim JC, Kim JH, Kim KW, Kim MD, Kim MM, Kim PK, Kim SW, Kim SY, Kim YS, Kim Y, Kimchi A, Kimmelman AC, Kimura T, King JS, Kirkegaard K, Kirkin V, Kirshenbaum LA, Kishi S, Kitajima Y, Kitamoto K, Kitaoka Y, Kitazato K, Kley RA, Klimecki WT, Klinkenberg M, Klucken J, Knævelsrud H, Knecht E, Knuppertz L, Ko JL, Kobayashi S, Koch JC, Koechlin-Ramonatxo C, Koenig U, Koh YH, Köhler K, Kohlwein SD, Koike M, Komatsu M, Kominami E, Kong D, Kong HJ, Konstantakou EG, Kopp BT, Korcsmaros T, Korhonen L, Korolchuk VI, Koshkina N V., Kou Y, Koukourakis MI, Koumenis C, Kovács AL, Kovács T, Kovacs WJ, Koya D, Kraft C, Krainc D, Kramer H, Kravic-Stevovic T, Krek W, Kretz-Remy C, Krick R, Krishnamurthy M, Kriston-Vizi J, Kroemer G, Kruer MC, Kruger R, Ktistakis NT, Kuchitsu K, Kuhn C, Kumar AP, Kumar A, Kumar A, Kumar D, Kumar D, Kumar R, Kumar S, Kundu M, Kung HJ, Kuno A, Kuo SH, Kuret J, Kurz T, Kwok T, Kwon TK, Kwon YT, Kyrmizi I, La Spada AR, Lafont F, Lahm T, Lakkaraju A, Lam T, Lamark T, Lancel S, Landowski TH, Lane DJR, Lane JD, Lanzi C, Lapaquette P, Lapierre LR, Laporte J, Laukkarinen J, Laurie GW, Lavandero S, Lavie L, Lavoie MJ, Law BYK, Law HKW, Law KB, Layfield R, Lazo PA, Le Cam L, Le Roch KG, Le Stunff H, Leardkamolkarn V, Lecuit M, Lee BH, Lee CH, Lee EF, Lee GM, Lee HJ, Lee H, Lee JK, Lee J, Lee JH, Lee JH, Lee M, Lee MS, Lee PJ, Lee SW, Lee SJ, Lee SJ, Lee SY, Lee SH, Lee SS, Lee SJ, Lee S, Lee YR, Lee YJ, Lee YH, Leeuwenburgh C, Lefort S, Legouis R, Lei J, Lei QY, Leib DA, Leibowitz G, Lekli I, Lemaire SD, Lemasters JJ, Lemberg MK, Lemoine A, Leng S, Lenz G, Lenzi P, Lerman LO, Barbato DL, Leu JIJ, Leung HY, Levine B, Lewis PA, Lezoualch F, Li C, Li F, Li FJ, Li J, Li K, Li L, Li M, Li Q, Li R, Li S, Li W, Li X, Li Y, Lian J, Liang C, Liang Q, Liao Y, Liberal J, Liberski PP, Lie P, Lieberman AP, Lie P, Lieber man AP, Lim HJ, Lim KL, Lim K, Lima RT, Lin CS, Lin CF, Lin F, Lin F, Lin FC, Lin K, Lin KH, Lin PH, Lin T, Lin WW, Lin YS, Lin Y, Linden R, Lindholm D, Lindqvist LM, Lingor P, Linkermann A, Liotta LA, Lipinski MM, Lira VA, Lisanti MP, Liton PB, Liu B, Liu C, Liu CF, Liu F, Liu HJ, Liu J, Liu JJ, Liu JL, Liu K, Liu L, Liu L, Liu Q, Liu RY, Liu S, Liu S, Liu W, Liu X De, Liu X, Liu XH, Liu X, Liu X, Liu X, Liu Y, Liu Y, Liu Z, Liu Z, Liuzzi JP, Lizard G, Ljujic M, Lodhi IJ, Logue SE, Lokeshwar BL, Long YC, Lonial S, Loos B, López-Otín C, López-Vicario C, Lorente M, Lorenzi PL, Lõrincz P, Los M, Lotze MT, Lovat PE, Lu B, Lu B, Lu J, Lu Q, Lu SM, Lu S, Lu Y, Luciano F, Luckhart S, Lucocq JM, Ludovico P, Lugea A, Lukacs NW, Lum JJ, Lund AH, Luo H, Luo J, Luo S, Luparello C, Lyons T, Ma J, Ma Y, Ma Y, Ma Z, Machado J, Machado-Santelli GM, Macian F, MacIntosh GC, MacKeigan JP, Macleod KF, MacMicking JD, MacMillan-Crow LA, Madeo F, Madesh M, Madrigal-Matute J, Maeda A, Maeda T, Maegawa G, Maellaro E, Maes H, Magariños M, Maiese K, Maiti TK, Maiuri L, Maiuri MC, Maki CG, Malli R, Malorni W, Maloyan A, Mami-Chouaib F, Man N, Mancias JD, Mandelkow EM, Mandell MA, Manfredi AA, Manié SN, Manzoni C, Mao K, Mao Z, Mao ZW, Marambaud P, Marconi AM, Marelja Z, Marfe G, Margeta M, Margittai E, Mari M, Mariani F V., Marin C, Marinelli S, Mariño G, Markovic I, Marquez R, Martelli AM, Martens S, Martin KR, Martin SJ, Martin S, Martin-Acebes MA, Martín-Sanz P, Martinand-Mari C, Martinet W, Martinez J, Martinez-Lopez N, Martinez-Outschoorn U, Martínez-Velázquez M, Martinez-Vicente M, Martins WK, Mashima H, Mastrianni JA, Matarese G, Matarrese P, Mateo R, Matoba S, Matsumoto N, Matsushita T, Matsuura A, Matsuzawa T, Mattson MP, Matus S, Maugeri N, Mauvezin C, Mayer A, Maysinger D, Mazzolini GD, McBrayer MK, McCall K, McCormick C, McInerney GM, McIver SC, McKenna S, McMahon JJ, McNeish IA, Mechta-Grigoriou F, Medema JP, Medina DL, Megyeri K, Mehrpour M, Mehta JL, Mei Y, Meier UC, Meijer AJ, Meléndez A, Melino G, Melino S, de Melo EJT, Mena MA, Meneghini MD, Menendez JA, Menezes R, Meng L, Meng LH, Meng S, Menghini R, Menko AS, Menna-Barreto RFS, Menon MB, Meraz-Ríos MA, Merla G, Merlini L, Merlot AM, Meryk A, Meschini S, Meyer JN, Mi MT, Miao CY, Micale L, Michaeli S, Michiels C, Migliaccio AR, Mihailidou AS, Mijaljica D, Mikoshiba K, Milan E, Miller-Fleming L, Mills GB, Mills IG, Minakaki G, Minassian BA, Ming XF, Minibayeva F, Minina EA, Mintern JD, Minucci S, Miranda-Vizuete A, Mitchell CH, Miyamoto S, Miyazawa K, Mizushima N, Mnich K, Mograbi B, Mohseni S, Moita LF, Molinari M, Molinari M, Møller AB, Mollereau B, Mollinedo F, Mongillo M, Monick MM, Montagnaro S, Montell C, Moore DJ, Moore MN, Mora-Rodriguez R, Moreira PI, Morel E, Morelli MB, Moreno S, Morgan MJ, Moris A, Moriyasu Y, Morrison JL, Morrison LA, Morselli E, Moscat J, Moseley PL, Mostowy S, Motori E, Mottet D, Mottram JC, Moussa CEH, Mpakou VE, Mukhtar H, Levy JMM, Muller S, Muñoz-Moreno R, Muñoz-Pinedo C, Münz C, Murphy ME, Murray JT, Murthy A, Mysorekar IU, Nabi IR, Nabissi M, Nader GA, Nagahara Y, Nagai Y, Nagata K, Nagelkerke A, Nagy P, Naidu SR, Nair S, Nakano H, Nakatogawa H, Nanjundan M, Napolitano G, Naqvi NI, Nardacci R, Narendra DP, Narita M, Nascimbeni AC, Natarajan R, Navegantes LC, Nawrocki ST, Nazarko TY, Nazarko VY, Neill T, Neri LM, Netea MG, Netea-Maier RT, Neves BM, Ney PA, Nezis IP, Nguyen HTT, Nguyen HP, Nicot AS, Nilsen H, Nilsson P, Nishimura M, Nishino I, Niso-Santano M, Niu H, Nixon RA, Njar VCO, Noda T, Noegel AA, Nolte EM, Norberg E, Norga KK, Noureini SK, Notomi S, Notterpek L, Nowikovsky K, Nukina N, Nürnberger T, O’donnell VB, O’donovan T, O’dwyer PJ, Oehme I, Oeste CL, Ogawa M, Ogretmen B, Ogura Y, Oh YJ, Ohmuraya M, Ohshima T, Ojha R, Okamoto K, Okazaki T, Oliver FJ, Ollinger K, Olsson S, Orban DP, Ordonez P, Orhon I, Orosz L, O’rourke EJ, Orozco H, Ortega AL, Ortona E, Osellame LD, Oshima J, Oshima S, Osiewacz HD, Otomo T, Otsu K, Ou JHJ, Outeiro TF, Ouyang DY, Ouyang H, Overholtzer M, Ozbun MA, Ozdinler PH, Ozpolat B, Pacelli C, Paganetti P, Page G, Pages G, Pagnini U, Pajak B, Pak SC, Pakos-Zebrucka K, Pakpour N, Palková Z, Palladino F, Pallauf K, Pallet N, Palmieri M, Paludan SR, Palumbo C, Palumbo S, Pampliega O, Pan H, Pan W, Panaretakis T, Pandey A, Pantazopoulou A, Papackova Z, Papademetrio DL, Papassideri I, Papini A, Parajuli N, Pardo J, Parekh V V., Parenti G, Park JI, Park J, Park OK, Parker R, Parlato R, Parys JB, Parzych KR, Pasquet JM, Pasquier B, Pasumarthi KBS, Patschan D, Pattingre S, Pattison S, Pause A, Pavenstädt H, Pavone F, Pedrozo Z, Peña FJ, Peñalva MA, Pende M, Peng J, Penna F, Penninger JM, Pensalfini A, Pepe S, Pereira GJS, Pereira PC, de la Cruz VP, Pérez-Pérez ME, Pérez-Rodríguez D, Pérez-Sala D, Perier C, Perl A, Perlmutter DH, Perrotta I, Pervaiz S, Pesonen M, Pessin JE, Peters GJ, Petersen M, Petrache I, Petrof BJ, Petrovski G, Phang JM, Piacentini M, Pierdominici M, Pierre P, Pierrefite-Carle V, Pietrocola F, Pimentel-Muiños FX, Pinar M, Pineda B, Pinkas-Kramarski R, Pinti M, Pinton P, Piperdi B, Piret JM, Platanias LC, Platta HW, Plowey ED, Pöggeler S, Poirot M, Polčic P, Poletti A, Poon AH, Popelka H, Popova B, Poprawa I, Poulose SM, Poulton J, Powers SK, Powers T, Pozuelo-Rubio M, Prak K, Prange R, Prescott M, Priault M, Prince S, Proia RL, Proikas-Cezanne T, Prokisch H, Promponas VJ, Przyklenk K, Puertollano R, Pugazhenthi S, Puglielli L, Pujol A, Puyal J, Pyeon D, Qi X, Qian W Bin, Qin ZH, Qiu Y, Qu Z, Quadrilatero J, Quinn F, Raben N, Rabinowich H, Radogna F, Ragusa MJ, Rahmani M, Raina K, Ramanadham S, Ramesh R, Rami A, Randall-Demllo S, Randow F, Rao H, Rao VA, Rasmussen BB, Rasse TM, Ratovitski EA, Rautou PE, Ray SK, Razani B, Reed BH, Reggiori F, Rehm M, Reichert AS, Rein T, Reiner DJ, Reits E, Ren J, Ren X, Renna M, Reusch JEB, Revuelta JL, Reyes L, Rezaie AR, Richards RI, Richardson DR, Richetta C, Riehle MA, Rihn BH, Rikihisa Y, Riley BE, Rimbach G, Rippo MR, Ritis K, Rizzi F, Rizzo E, Roach PJ, Robbins J, Roberge M, Roca G, Roccheri MC, Rocha S, Rodrigues CMP, Rodríguez CI, de Cordoba SR, Rodriguez-Muela N, Roelofs J, Rogov V V., Rohn TT, Rohrer B, Romanelli D, Romani L, Romano PS, Roncero MIG, Rosa JL, Rosello A, Rosen K V., Rosenstiel P, Rost-Roszkowska M, Roth KA, Roué G, Rouis M, Rouschop KM, Ruan DT, Ruano D, Rubinsztein DC, Rucker EB, Rudich A, Rudolf E, Rudolf R, Ruegg MA, Ruiz-Roldan C, Ruparelia AA, Rusmini P, Russ DW, Russo GL, Russo G, Russo R, Rusten TE, Ryabovol V, Ryan KM, Ryter SW, Sabatini DM, Sacher M, Sachse C, Sack MN, Sadoshima J, Saftig P, Sagi-Eisenberg R, Sahni S, Saikumar P, Saito T, Saitoh T, Sakakura K, Sakoh-Nakatogawa M, Sakuraba Y, Salazar-Roa M, Salomoni P, Saluja AK, Salvaterra PM, Salvioli R, Samali A, Sanchez AMJ, Sánchez-Alcázar JA, Sanchez-Prieto R, Sandri M, Sanjuan MA, Santaguida S, Santambrogio L, Santoni G, Dos Santos CN, Saran S, Sardiello M, Sargent G, Sarkar P, Sarkar S, Sarrias MR, Sarwal MM, Sasakawa C, Sasaki M, Sass M, Sato K, Sato M, Satriano J, Savaraj N, Saveljeva S, Schaefer L, Schaible UE, Scharl M, Schatzl HM, Schekman R, Scheper W, Schiavi A, Schipper HM, Schmeisser H, Schmidt J, Schmitz I, Schneider BE, Schneider EM, Schneider JL, Schon EA, Schönenberger MJ, Schönthal AH, Schorderet DF, Schröder B, Schuck S, Schulze RJ, Schwarten M, Schwarz TL, Sciarretta S, Scotto K, Scovassi AI, Screaton RA, Screen M, Seca H, Sedej S, Segatori L, Segev N, Seglen PO, Seguí-Simarro JM, Segura-Aguilar J, Seiliez I, Seki E, Sell C, Semenkovich CF, Semenza GL, Sen U, Serra AL, Serrano-Puebla A, Sesaki H, Setoguchi T, Settembre C, Shacka JJ, Shajahan-Haq AN, Shapiro IM, Sharma S, She H, Shen CKJ, Shen CC, Shen HM, Shen S, Shen W, Sheng R, Sheng X, Sheng ZH, Shepherd TG, Shi J, Shi Q, Shi Q, Shi Y, Shibutani S, Shibuya K, Shidoji Y, Shieh JJ, Shih CM, Shimada Y, Shimizu S, Shin DW, Shinohara ML, Shintani M, Shintani T, Shioi T, Shirabe K, Shiri-Sverdlov R, Shirihai O, Shore GC, Shu CW, Shukla D, Sibirny AA, Sica V, Sigurdson CJ, Sigurdsson EM, Sijwali PS, Sikorska B, Silveira WA, Silvente-Poirot S, Silverman GA, Simak J, Simmet T, Simon AK, Simon HU, Simone C, Simons M, Simonsen A, Singh R, Singh S V., Singh SK, Sinha D, Sinha S, Sinicrope FA, Sirko A, Sirohi K, Sishi BJN, Sittler A, Siu PM, Sivridis E, Skwarska A, Slack R, Slaninová I, Slavov N, Smaili SS, Smalley KSM, Smith DR, Soenen SJ, Soleimanpour SA, Solhaug A, Somasundaram K, Son JH, Sonawane A, Song C, Song F, Song HK, Song JX, Song W, Soo KY, Sood AK, Soong TW, Soontornniyomkij V, Sorice M, Sotgia F, Soto-Pantoja DR, Sotthibundhu A, Sousa MJ, Spaink HP, Span PN, Spang A, Sparks JD, Speck PG, Spector SA, Spies CD, Springer W, Clair DS, Stacchiotti A, Staels B, Stang MT, Starczynowski DT, Starokadomskyy P, Steegborn C, Steele JW, Stefanis L, Steffan J, Stellrecht CM, Stenmark H, Stepkowski TM, Stern ST, Stevens C, Stockwell BR, Stoka V, Storchova Z, Stork B, Stratoulias V, Stravopodis DJ, Strnad P, Strohecker AM, Ström AL, Stromhaug P, Stulik J, Su YX, Su Z, Subauste CS, Subramaniam S, Sue CM, Suh SW, Sui X, Sukseree S, Sulzer D, Sun FL, Sun J, Sun J, Sun SY, Sun Y, Sun Y, Sun Y, Sundaramoorthy V, Sung J, Suzuki H, Suzuki K, Suzuki N, Suzuki T, Suzuki YJ, Swanson MS, Swanton C, Swärd K, Swarup G, Sweeney ST, Sylvester PW, Szatmari Z, Szegezdi E, Szlosarek PW, Taegtmeyer H, Tafani M, Taillebourg E, Tait SWG, Takacs-Vellai K, Takahashi Y, Takáts S, Takemura G, Takigawa N, Talbot NJ, Tamagno E, Tamburini J, Tan CP, Tan L, Tan ML, Tan M, Tan YJ, Tanaka K, Tanaka M, Tang D, Tang D, Tang G, Tanida I, Tanji K, Tannous BA, Tapia JA, Tasset-Cuevas I, Tatar M, Tavassoly I, Tavernarakis N, Taylor A, Taylor GS, Taylor GA, Taylor JP, Taylor MJ, Tchetina E V., Tee AR, Teixeira-Clerc F, Telang S, Tencomnao T, Teng BB, Teng RJ, Terro F, Tettamanti G, Theiss AL, Theron AE, Thomas KJ, Thomé MP, Thomes PG, Thorburn A, Thorner J, Thum T, Thumm M, Thurston TLM, Tian L, Till A, Ting JPY, Ting JPY, Titorenko VI, Toker L, Toldo S, Tooze SA, Topisirovic I, Torgersen ML, Torosantucci L, Torriglia A, Torrisi MR, Tournier C, Towns R, Trajkovic V, Travassos LH, Triola G, Tripathi DN, Trisciuoglio D, Troncoso R, Trougakos IP, Truttmann AC, Tsai KJ, Tschan MP, Tseng YH, Tsukuba T, Tsung A, Tsvetkov AS, Tu S, Tuan HY, Tucci M, Tumbarello DA, Turk B, Turk V, Turner RFB, Tveita AA, Tyagi SC, Ubukata M, Uchiyama Y, Udelnow A, Ueno T, Umekawa M, Umemiya-Shirafuji R, Underwood BR, Ungermann C, Ureshino RP, Ushioda R, Uversky VN, Uzcátegui NL, Vaccari T, Vaccaro MI, Váchová L, Vakifahmetoglu-Norberg H, Valdor R, Valente EM, Vallette F, Valverde AM, Van den Berghe G, Van Den Bosch L, van den Brink GR, van der Goot FG, van der Klei IJ, van der Laan LJW, van Doorn WG, van Egmond M, van Golen KL, Van Kaer L, Campagne M van L, Vandenabeele P, Vandenberghe W, Vanhorebeek I, Varela-Nieto I, Vasconcelos MH, Vasko R, Vavvas DG, Vega-Naredo I, Velasco G, Velentzas AD, Velentzas PD, Vellai T, Vellenga E, Vendelbo MH, Venkatachalam K, Ventura N, Ventura S, Veras PST, Verdier M, Vertessy BG, Viale A, Vidal M, Vieira HLA, Vierstra RD, Vigneswaran N, Vij N, Vila M, Villar M, Villar VH, Villarroya J, Vindis C, Viola G, Viscomi MT, Vitale G, Vogl DT, Voitsekhovskaja O V., von Haefen C, von Schwarzenberg K, Voth DE, Vouret-Craviari V, Vuori K, Vyas JM, Waeber C, Walker CL, Walker MJ, Walter J, Wan L, Wan X, Wang B, Wang C, Wang CY, Wang C, Wang C, Wang C, Wang D, Wang F, Wang F, Wang G, Wang HJ, Wang H, Wang HG, Wang H, Wang HD, Wang J, Wang J, Wang M, Wang MQ, Wang PY, Wang P, Wang RC, Wang S, Wang TF, Wang X, Wang XJ, Wang XW, Wang X, Wang X, Wang Y, Wang Y, Wang Y, Wang YJ, Wang Y, Wang Y, Wang YT, Wang Y, Wang ZN, Wappner P, Ward C, Ward DMV, Warnes G, Watada H, Watanabe Y, Watase K, Weaver TE, Weekes CD, Wei J, Weide T, Weihl CC, Weindl G, Weis SN, Wen L, Wen X, Wen Y, Westermann B, Weyand CM, White AR, White E, Whitton JL, Whitworth AJ, Wiels J, Wild F, Wildenberg ME, Wileman T, Wilkinson DS, Wilkinson S, Willbold D, Williams C, Williams K, Williamson PR, Winklhofer KF, Witkin SS, Wohlgemuth SE, Wollert T, Wolvetang EJ, Wong E, Wong GW, Wong RW, Wong VKW, Woodcock EA, Wright KL, Wu C, Wu D, Wu GS, Wu J, Wu J, Wu M, Wu M, Wu S, Wu WKK, Wu Y, Wu Z, Xavier CPR, Xavier RJ, Xia GX, Xia T, Xia W, Xia Y, Xiao H, Xiao J, Xiao S, Xiao W, Xie CM, Xie Z, Xie Z, Xilouri M, Xiong Y, Xu C, Xu C, Xu F, Xu H, Xu H, Xu J, Xu J, Xu J, Xu L, Xu X, Xu Y, Xu Y, Xu ZX, Xu Z, Xue Y, Yamada T, Yamamoto A, Yamanaka K, Yamashina S, Yamashiro S, Yan B, Yan B, Yan X, Yan Z, Yanagi Y, Yang DS, Yang JM, Yang L, Yang M, Yang PM, Yang P, Yang Q, Yang W, Yang WY, Yang X, Yang Y, Yang Y, Yang Z, Yang Z, Yao MC, Yao PJ, Yao X, Yao Z, Yao Z, Yasui LS, Ye M, Yedvobnick B, Yeganeh B, Yeh ES, Yeyati PL, Yi F, Yi L, Yin XM, Yip CK, Yoo YM, Yoo YH, Yoon SY, Yoshida KI, Yoshimori T, Young KH, Yu H, Yu JJ, Yu JT, Yu J, Yu L, Yu WH, Yu XF, Yu Z, Yuan J, Yuan ZM, Yue BYJT, Yue J, Yue Z, Zacks DN, Zacksenhaus E, Zaffaroni N, Zaglia T, Zakeri Z, Zecchini V, Zeng J, Zeng M, Zeng Q, Zervos AS, Zhang DD, Zhang F, Zhang G, Zhang GC, Zhang H, Zhang H, Zhang H, Zhang J, Zhang J, Zhang J, Zhang JP, Zhang L, Zhang L, Zhang L, Zhang MY, Zhang X, Zhang XD, Zhang Y, Zhang Y, Zhang Y, Zhang Y, Zhang Y, Zhao M, Zhao WL, Zhao X, Zhao YG, Zhao Y, Zhao Y, Zhao YX, Zhao Z, Zhao ZJ, Zheng D, Zheng XL, Zheng X, Zhivotovsky B, Zhong Q, Zhou GZ, Zhou G, Zhou H, Zhou SF, Zhou XJ, Zhu H, Zhu H, Zhu WG, Zhu W, Zhu XF, Zhu Y, Zhuang SM, Zhuang X, Ziparo E, Zois CE, Zoladek T, Zong WX, Zorzano A, Zughaier SM. 2016. Guidelines for the use and interpretation of assays for monitoring autophagy. Autophagy 12:1–222. doi:10.1080/15548627.2015.1100356

Klionsky DJ, Schulman BA. 2014. Dynamic regulation of macroautophagy by distinctive ubiquitin-like proteins. Nat Struct Mol Biol 21:336–345. doi:10.1038/nsmb.2787

Kobayashi Y, Yang S, Nykamp K, Garcia J, Lincoln SE, Topper SE. 2017. Pathogenic variant burden in the ExAC database: An empirical approach to evaluating population data for clinical variant interpretation. Genome Med 9:1–14. doi:10.1186/s13073-017-0403-7

Kohlhoff KJ, Robustelli P, Cavalli A, Salvatella X, Vendruscolo M. 2009. Fast and accurate predictions of protein NMR chemical shifts from interatomic distances. J Am Chem Soc 131:13894–5. doi:10.1021/ja903772t

Kotlyar M, Pastrello C, Malik Z, Jurisica I. 2019. IID 2018 update: Context-specific physical protein-protein interactions in human, model organisms and domesticated species. Nucleic Acids Res 47:D581–D589. doi:10.1093/nar/gky1037

Kouno T, Mizuguchi M, Tanidal I, Uenol T, Kanematsu T, Mori Y, Shinoda H, Hirata M, Kominami E, Kawano K. 2005. Solution structure of microtubule-associated protein light chain 3 and identification of its functional subdomains. J Biol Chem 280:24610–24617. doi:10.1074/jbc.M413565200

Kraft LJ, Nguyen TA, Vogel SS, Kenworthy AK. 2014. Size, stoichiometry, and organization of soluble LC3-associated complexes. Autophagy 10:861–877. doi:10.4161/auto.28175

Kuang Y, Ma K, Zhou C, Ding P, Zhu Y, Chen Q, Xia B. 2016a. Structural Basis for the Phosphorylation of FUNDC1 LIR as a Molecular Switch of Mitophagy. Autophagy 1–26.

Kuang Y, Ma K, Zhou C, Ding P, Zhu Y, Chen Q, Xia B. 2016b. Structural basis for the phosphorylation of FUNDC1 LIR as a molecular switch of mitophagy. Autophagy 12:2363–2373. doi:10.1080/15548627.2016.1238552

Kubli DA, Gustafsson ÅB. 2012. Mitochondria and mitophagy: The yin and yang of cell death control. Circ Res 111:1208–1221. doi:10.1161/CIRCRESAHA.112.265819

Kwon DH, Kim L, Kim BW, Kim JH, Roh KH, Choi EJ, Song HK. 2017. A novel conformation of the LC3-interacting region motif revealed by the structure of a complex between LC3B and RavZ. Biochem Biophys Res Commun 490:1093–1099. doi:10.1016/j.bbrc.2017.06.173

Lambrughi M, De Gioia L, Gervasio FL, Lindorff-Larsen K, Nussinov R, Urani C, Bruschi M, Papaleo E. 2016. DNA-binding protects p53 from interactions with cofactors involved in transcription-independent functions. Nucleic Acids Res 44:9096–9109. doi:10.1093/nar/gkw770

Lange OF, van der Spoel D, de Groot BL. 2010. Scrutinizing molecular mechanics force fields on the submicrosecond timescale with NMR data. Biophys J 99:647–55. doi:10.1016/j.bpj.2010.04.062

Lazova R, Camp RL, Klump V, Siddiqui SF, Amaravadi RK, Pawelek JM. 2012. Punctate LC3B expression is a common feature of solid tumors and associated with proliferation, metastasis, and poor outcome. Clin Cancer Res 18:370–379. doi:10.1158/1078-0432.CCR-11-1282

Lee YK, Lee JA. 2016. Role of the mammalian ATG8/LC3 family in autophagy: Differential and compensatory roles in the spatiotemporal regulation of autophagy. BMB Rep 49:424–430. doi:10.5483/BMBRep.2016.49.8.081

Li D, Brüschweiler R. 2015. PPM-One: A static protein structure based chemical shift predictor. J Biomol NMR 62:403–409. doi:10.1007/s10858-015-9958-z

Li DW, Brüschweiler R. 2012. PPM: A side-chain and backbone chemical shift predictor for the assessment of protein conformational ensembles. J Biomol NMR 54:257–265. doi:10.1007/s10858-012-9668-8

Li DW, Brüschweiler R. 2010a. Certification of molecular dynamics trajectories with NMR chemical shifts. J Phys Chem Lett 1:246–248. doi:10.1021/jz9001345

Li DW, Brüschweiler R. 2010b. NMR-based protein potentials. Angew Chemie - Int Ed 49:6778–6780. doi:10.1002/anie.201001898

Li J, Zhao T, Zhang Y, Zhang K, Shi L, Chen Y, Wang X, Sun Z. 2018. Performance evaluation of pathogenicity-computation methods for missense variants. Nucleic Acids Res 46:7793–7804. doi:10.1093/nar/gky678

Li S, Elcock AH. 2015. Residue-Specific Force Field (RSFF2) improves the modeling of conformational behavior of peptides and proteins. J Phys Chem Lett 6:2127–2133. doi:10.1021/acs.jpclett.5b00654

Lindorff-Larsen K, Maragakis P, Piana S, Eastwood MP, Dror RO, Shaw DE. 2012. Systematic validation of protein force fields against experimental data. PLoS One 7:e32131. doi:10.1371/journal.pone.0032131

Lindorff-Larsen K, Maragakis P, Piana S, Shaw DE. 2016. Picosecond to Millisecond Structural Dynamics in Human Ubiquitin. J Phys Chem B acs.jpcb.6b02024. doi:10.1021/acs.jpcb.6b02024

Lindorff-Larsen K, Piana S, Palmo K, Maragakis P, Klepeis JL, Dror RO, Shaw DE. 2010. Improved side-chain torsion potentials for the Amber ff99SB protein force field. Proteins 78:1950–8. doi:10.1002/prot.22711

Liu C, Ma H, Wu J, Huang Q, Liu JO, Yu L. 2013. Arginine68 is an essential residue for the C-terminal cleavage of human Atg8 family proteins. BMC Cell Biol 14:27. doi:10.1186/1471-2121-14-27

Long D, Li DW, Walter KFA, Griesinger C, Brüschweiler R. 2011. Toward a predictive understanding of slow methyl group dynamics in proteins. Biophys J 101:910–915. doi:10.1016/j.bpj.2011.06.053

Lovell SC, Word JM, Richardson JS, Richardson DC. 2000. The penultimate rotamer library. Proteins 40:389–408.

Lv M, Wang C, Li F, Peng J, Wen B, Gong Q, Shi Y, Tang Y. 2016. Structural insights into the recognition of phosphorylated FUNDC1 by LC3B in mitophagy. Protein Cell. doi:10.1007/s13238-016-0328-8

MacKerell, AD, Bashford D, Dunbrack, RL, Evanseck JD, Field MJ, Fischer S, Gao J, Guo H, Ha S, Joseph-McCarthy D, Kuchnir L, Kuczera K, Lau FTK, Mattos C, Michnick S, Ngo T, Nguyen DT, Prodhom B, Reiher WE, Roux B, Schlenkrich M, Smith JC, Stote R, Straub J, Watanabe M, Wiórkiewicz-Kuczera J, Yin D, Karplus M. 1998. All-Atom Empirical Potential for Molecular Modeling and Dynamics Studies of Proteins. J Phys Chem B 102:3586–3616. doi:10.1021/jp973084f

Mah LY, Ryan KM. 2012. Autophagy and cancer. Cold Spring Harb Perspect Biol 4:a008821.

Maier JA, Martinez C, Kasavajhala K, Wickstrom L, Hauser KE, Simmerling C. 2015. ff14SB: Improving the accuracy of protein side chain and backbone parameters from ff99SB. J Chem Theory Comput 11:3696–3713. doi:10.4172/2157-7633.1000305.Improved

Mancias JD, Kimmelman AC. 2016. Mechanisms of Selective Autophagy in Normal Physiology and Cancer. J Mol Biol 428:1659–1680. doi:10.1016/j.jmb.2016.02.027

Maragakis P, Lindorff-Larsen K, Eastwood MP, Dror RO, Klepeis JL, Arkin IT, Jensen MØ, Xu H, Trbovic N, Friesner RA, Palmer AG, Shaw DE. 2008. Microsecond Molecular Dynamics Simulation Shows Effect of Slow Loop Dynamics on Backbone Amide Order Parameters of Proteins. J Phys Chem B 112:6155–6158. doi:10.1021/jp077018h

Mariani S, Dell’Orco D, Felline A, Raimondi F, Fanelli F. 2013. Network and Atomistic Simulations Unveil the Structural Determinants of Mutations Linked to Retinal Diseases. PLoS Comput Biol 9. doi:10.1371/journal.pcbi.1003207

Marshall RS, Hua Z, Mali S, McLoughlin F, Vierstra RD. 2019. ATG8-Binding UIM Proteins Define a New Class of Autophagy Adaptors and Receptors. Cell 177:766–781.e24. doi:10.1016/j.cell.2019.02.009

Martín-García F, Papaleo E, Gomez-Puertas P, Boomsma W, Lindorff-Larsen K. 2015. Comparing Molecular Dynamics Force Fields in the Essential Subspace. PLoS One 10:e0121114. doi:10.1371/journal.pone.0121114

Maruyama Y, Sou YS, Kageyama S, Takahashi T, Ueno T, Tanaka K, Komatsu M, Ichimura Y. 2014. LC3B is indispensable for selective autophagy of p62 but not basal autophagy. Biochem Biophys Res Commun 446:309–315. doi:10.1016/j.bbrc.2014.02.093

McEwan DG, Popovic D, Gubas A, Terawaki S, Suzuki H, Stadel D, Coxon FP, MirandadeStegmann D, Bhogaraju S, Maddi K, Kirchof A, Gatti E, Helfrich MH, Wakatsuki S, Behrends C, Pierre P, Dikic I. 2015. PLEKHM1 regulates autophagosome-lysosome fusion through HOPS complex and LC3/GABARAP proteins. Mol Cell 57:39–54. doi:10.1016/j.molcel.2014.11.006

Mercadante D, Gräter F, Daday C. 2018. CONAN: A Tool to Decode Dynamical Information from Molecular Interaction Maps. Biophys J 114:1267–1273. doi:10.1016/j.bpj.2018.01.033

Michaud-Agrawal N, Denning EJ, Woolf TB, Beckstein O. 2011. MDAnalysis: a toolkit for the analysis of molecular dynamics simulations. J Comput Chem 32:2319–2327. doi:10.1016/j.pestbp.2011.02.012.Investigations

Mikhaylova O, Stratton Y, Hall D, Kellner E, Ehmer B, Drew AF, Gallo CA, Plas DR, Biesiada J, Meller J, Czyzyk-Krzeska MF. 2012. VHL-Regulated MiR-204 Suppresses Tumor Growth through Inhibition of LC3B-Mediated Autophagy in Renal Clear Cell Carcinoma. Cancer Cell 21:532– 546. doi:10.1016/j.ccr.2012.02.019

Mizushima N. 2018. A brief history of autophagy from cell biology to physiology and disease. Nat Cell Biol 20:521–527. doi:10.1038/s41556-018-0092-5

Mizushima N, Yoshimori T, Levine B. 2010. Methods in mammalian autophagy research. Cell 140:313–26. doi:10.1016/j.cell.2010.01.028

Mohan J, Wollert T. 2018. Human ubiquitin-like proteins as central coordinators in autophagy. Interface Focus 8:20180025. doi:10.1098/rsfs.2018.0025

Nakatogawa H, Ichimura Y, Ohsumi Y. 2007. Atg8, a Ubiquitin-like Protein Required for Autophagosome Formation, Mediates Membrane Tethering and Hemifusion. Cell 130:165–178. doi:10.1016/j.cell.2007.05.021

Nielsen S V., Stein A, Dinitzen AB, Papaleo E, Tatham MH, Poulsen EG, Kassem MM, Rasmussen LJ, Lindorff-Larsen K, Hartmann-Petersen R. 2017. Predicting the impact of Lynch syndrome-causing missense mutations from structural calculations. PLOS Genet 13:e1006739. doi:10.1371/journal.pgen.1006739

Noda NN, Kumeta H, Nakatogawa H, Satoo K, Adachi W, Ishii J, Fujioka Y, Ohsumi Y, Inagaki F. 2008. Structural basis of target recognition by Atg8/LC3 during selective autophagy. Genes to Cells 13:1211–1218. doi:10.1111/j.1365-2443.2008.01238.x

Noda NN, Ohsumi Y, Inagaki F. 2010. Atg8-family interacting motif crucial for selective autophagy. FEBS Lett 584:1379–1385. doi:10.1016/j.febslet.2010.01.018

Nuta GC, Gilad Y, Gershoni M, Sznajderman A, Schlesinger T, Bialik S, Eisenstein M, Pietrokovski S, Kimchi A. 2018. A cancer associated somatic mutation in LC3B attenuates its binding to E1-like ATG7 protein and subsequent lipidation. Autophagy 0:15548627.2018.1525476. doi:10.1080/15548627.2018.1525476

Nygaard M, Terkelsen T, Olsen AV, Sora V, Salamanca J, Rizza F, Bergstrand S, Marco M Di, Vistesen M, Lambrughi M, Jaattela M, Kallunki T, Papaleo E. 2016. The mutational landscape of the oncogenic MZF1 SCAN domain in cancer. Front Mol Biosci 3:1–18. doi:10.3389/fmolb.2016.00078

Olsvik HL, Lamark T, Takagi K, Larsen KB, Evjen G, Øvervatn A, Mizushima T, Johansen XT. 2015. FYCO1 contains a C-terminally extended, LC3A/B-preferring LC3-interacting region (LIR) motif required for efficient maturation of autophagosomes during basal autophagy. J Biol Chem 290:29361–29374. doi:10.1074/jbc.M115.686915

Ovchinnikov S, Kamisetty H, Baker D. 2014. Robust and accurate prediction of residue–residue interactions across protein interfaces using evolutionary information. Elife 3:1–21. doi:10.7554/eLife.02030

Palmer AG. 2015. Enzyme dynamics from NMR spectroscopy. Acc Chem Res 48:457–65. doi:10.1021/ar500340a

Pandini A, Fornili A. 2016. Using Local States to Drive the Sampling of Global Conformations in Proteins. J Chem Theory Comput 12:1368–1379. doi:10.1021/acs.jctc.5b00992

Pandini A, Fornili A, Fraternali F, Kleinjung J. 2013. GSATools: Analysis of allosteric communication and functional local motions using a structural alphabet. Bioinformatics 29:2053–2055. doi:10.1093/bioinformatics/btt326

Pandini A, Fornili A, Kleinjung J, Corey R, Pauling L, Jones T, Thirup S, Ramachandran G, Ramakrishnan C, Sasisekharan V, Walther D, Cohen F, Rooman M, Rodriguez J, Wodak S, Park B, Levitt M, Bystroff C, Baker D, Micheletti C, Seno F, Maritan A, Brevern A de, Etchebest C, Hazout S, Kolodny R, Koehl P, Guibas L, Levitt M, Camproux A, Gautier R, Tufféry P, Tung C, Huang J, Yang J, Offmann B, Tyagi M, Brevern A de, Hunter C, Subramaniam S, Camproux A, Tuffery P, Chevrolat J, Boisvieux J, Hazout S, Kolodny R, Levitt M, Fourrier L, Benros C, Brevern A de, Etchebest C, Benros C, Hazout S, Brevern A, Friedberg I, Harder T, Kolodny R, Sitbon E, Li Z, Godzik A, Schenk G, Margraf T, Torda A, Guyon F, Camproux A, Hochez J, Tufféry P, Yang J, Tung C, Tung C, Yang J, Tyagi M, Brevern A de, Srinivasan N, Offmann B, Pandini A, Bonati L, Fraternali F, Kleinjung J, Maupetit J, Gautier R, Tufféry P, Le Q, Pollastri G, Koehl P, Deschavanne P, Tufféry P, Tuffery P, Derreumaux P, Maupetit J, Derreumaux P, Tuffery P, Maupetit J, Derreumaux P, Tufféry P, MacDonald J, Maksimiak K, Sadowski M, Taylor W, Chandonia J, Hon G, Walker N, Conte L, Koehl P, Levitt M, Brenner S, Brenner S, Koehl P, Levitt M, Ankerst M, Breunig M, Kriegel H, Sander J, Daszykowski M, Walczak B, Massart D, Kriegel H, Brecheisen S, Januzaj E, Kröger P, Venables W, Ripley B, Theobald D, Akaike H, Konishi S, Kitagawa G, Mitchell M, Bahar I, Atilgan A, Erman B, Haliloglu T, Bahar I, Erman B, Bahar I, Rader A, Hess B, Groot B de, Aalten D van, Scheek R, Amadei A, Vriend G, Berendsen H, Seeliger D, Haas J, Groot B de, Seeliger D, Groot B De, Fernández A, Berry R, Barrett C, Hall B, Noble M, Eyrisch S, Helms V, Kabsch W, Sander C, Spoel Dv Der, Lindahl E, Hess B, Groenhof G, Mark A, Berendsen H, Kaminski G, Friesner R, Tirado-Rives J, Jorgensen W, Shannon C, Frishman D, Argos P, Kundu S, Melton J, Sorensen D, Phillips G, Martin J, Regad L, Etchebest C, Camproux A, Chen Y, Reilly K, Sprague A, Guan Z, Ligges U, Mächler M, Delano W. 2010. Structural alphabets derived from attractors in conformational space. BMC Bioinformatics 11:97. doi:10.1186/1471-2105-11-97

Pankiv S, Clausen TH, Lamark T, Brech A, Bruun JA, Outzen H, Øvervatn A, Bjørkøy G, Johansen T. 2007. p62/SQSTM1 binds directly to Atg8/LC3 to facilitate degradation of ubiquitinated protein aggregates by autophagy*[S]. J Biol Chem 282:24131–24145. doi:10.1074/jbc.M702824200

Papaleo E. 2015. Integrating atomistic molecular dynamics simulations, experiments, and network analysis to study protein dynamics: strength in unity. Front Mol Biosci 2:28. doi:10.3389/fmolb.2015.00028

Papaleo E, Camilloni C, Teilum K, Vendruscolo M, Lindorff-Larsen K. 2018. Molecular dynamics ensemble refinement of the heterogeneous native state of NCBD using chemical shifts and NOEs. PeerJ 6:e5125. doi:10.7717/peerj.5125

Papaleo E, Lindorff-larsen K, Gioia L De. 2012a. Paths of long-range communication in the E2 enzymes of family 3 : a molecular dynamics investigation w. Phys Chem Chem Phys 12515–12525. doi:10.1039/c2cp41224a

Papaleo E, Parravicini F, Grandori R, De Gioia L, Brocca S. 2014a. Structural investigation of the cold-adapted acylaminoacyl peptidase from Sporosarcina psychrophila by atomistic simulations and biophysical methods. Biochim Biophys Acta - Proteins Proteomics 1844:2203–2213. doi:10.1016/j.bbapap.2014.09.018

Papaleo E, Renzetti G, Tiberti M. 2012b. Mechanisms of Intramolecular Communication in a Hyperthermophilic Acylaminoacyl Peptidase: A Molecular Dynamics Investigation. PLoS One 7:e35686. doi:10.1371/journal.pone.0035686

Papaleo E, Saladino G, Lambrughi M, Lindorff-Larsen K, Gervasio FL, Nussinov R. 2016. The role of protein loops and linkers in conformational dynamics and allostery. Chem Rev 116:6391–6423. doi:10.1021/acs.chemrev.5b00623

Papaleo E, Sutto L, Gervasio FL, Lindorff-Larsen K. 2014b. Conformational changes and free energies in a proline isomerase. J Chem Theory Comput 10:4169–4174. doi:10.1021/ct500536r

Parzych KR, Klionsky DJ. 2013. An Overview of Autophagy: Morphology, Mechanism, and Regulation. Antioxid Redox Signal 20:460–473. doi:10.1089/ars.2013.5371

Pasi M, Tiberti M, Arrigoni A, Papaleo E. 2012. xPyder: a PyMOL plugin to analyze coupled residues and their networks in protein structures. J Chem Inf Model 279:1–6. doi:10.1021/ci300213c

Piana S, Klepeis JL, Shaw DE. 2014. Assessing the accuracy of physical models used in protein-folding simulations: quantitative evidence from long molecular dynamics simulations. Curr Opin Struct Biol 24:98–105. doi:10.1016/j.sbi.2013.12.006

Piana S, Lindorff-Larsen K, Shaw DE. 2011. How robust are protein folding simulations with respect to force field parameterization? Biophys J 100:L47-9. doi:10.1016/j.bpj.2011.03.051

Privalov PL. 1979. Stability of Proteins Small Globular Proteins. Adv Protein Chem 33:167–241. doi:10.1016/S0065-3233(08)60460-X

Qiu Y, Zheng Y, Wu KP, Schulman BA. 2017. Insights into links between autophagy and the ubiquitin system from the structure of LC3B bound to the LIR motif from the E3 ligase NEDD4. Protein Sci 26:1674–1680. doi:10.1002/pro.3186

Riera C, Lois S, de la Cruz X. 2014. Prediction of pathological mutations in proteins: the challenge of integrating sequence conservation and structure stability principles. Wiley Interdiscip Rev Comput Mol Sci 4:249–268. doi:10.1002/wcms.1170

Robustelli P, Piana S, Shaw DE. 2018. Developing a molecular dynamics force field for both folded and disordered protein states. Proc Natl Acad Sci 115:E4758–E4766. doi:10.1073/pnas.1800690115

Robustelli P, Stafford KA, Palmer AG. 2012. Interpreting Protein Structural Dynamics from NMR Chemical Shifts. J Am Chem Soc 134:6365–6374. doi:10.1021/ja300265w

Rogov V V., Suzuki H, Marinković M, Lang V, Kato R, Kawasaki M, Buljubašić M, Šprung M, Rogova N, Wakatsuki S, Hamacher-Brady A, Dötsch V, Dikic I, Brady NR, Novak I. 2017. Phosphorylation of the mitochondrial autophagy receptor Nix enhances its interaction with LC3 proteins. Sci Rep 7:1–12. doi:10.1038/s41598-017-01258-6

Rogov V, Dötsch V, Johansen T, Kirkin V. 2014. Interactions between Autophagy Receptors and Ubiquitin-like Proteins Form the Molecular Basis for Selective Autophagy. Mol Cell 53:167–178. doi:10.1016/j.molcel.2013.12.014

Rogov V V, Suzuki H, Fiskin E, Wild P, Kniss A, Rozenknop A, Kato R, Kawasaki M, McEwan DG, Löhr F, Güntert P, Dikic I, Wakatsuki S, Dötsch V. 2013. Structural basis for phosphorylation-triggered autophagic clearance of Salmonella. Biochem J 454:459–466. doi:10.1042/BJ20121907

Rybstein MD, Bravo-San Pedro JM, Kroemer G, Galluzzi L. 2018. The autophagic network and cancer. Nat Cell Biol 20:243–251. doi:10.1038/s41556-018-0042-2

Satoo K, Noda NN, Kumeta H, Fujioka Y, Mizushima N, Ohsumi Y, Inagaki F. 2009. The structure of Atg4B-LC3 complex reveals the mechanism of LC3 processing and delipidation during autophagy. EMBO J 28:1341–50. doi:10.1038/emboj.2009.80

Schaaf MBE, Keulers TG, Vooijs MA, Rouschop KMA. 2016. LC3/GABARAP family proteins: Autophagy-(un)related functions. FASEB J 30:3961–3978. doi:10.1096/fj.201600698R

Scheller R, Stein A, Nielsen S V., Marin FI, Gerdes A-M, Di Marco M, Papaleo E, Lindorff-Larsen K, Hartmann-Petersen R. 2019. Toward mechanistic models for genotype-phenotype correlations in phenylketonuria using protein stability calculations. Hum Mutat. doi:10.1002/humu.23707

Schymkowitz J, Borg J, Stricher F, Nys R, Rousseau F, Serrano L. 2005. The FoldX web server: an online force field. Nucleic Acids Res 33:W382-8. doi:10.1093/nar/gki387

Shieh AD, Hashimoto TB, Airoldi EM. 2011. Tree preserving embedding. Proc Natl Acad Sci 108:16916–16921. doi:10.1073/pnas.1018393108

Showalter SA, Brüschweiler R. 2007. Validation of Molecular Dynamics Simulations of Biomolecules Using NMR Spin Relaxation as Benchmarks: Application to the AMBER99SB Force Field. J Chem Theory Comput 3:961–75. doi:10.1021/ct7000045

Singh SS, Vats S, Chia AYQ, Tan TZ, Deng S, Ong MS, Arfuso F, Yap CT, Goh BC, Sethi G, Huang RYJ, Shen HM, Manjithaya R, Kumar AP. 2018. Dual role of autophagy in hallmarks of cancer. Oncogene 37:1142–1158. doi:10.1038/s41388-017-0046-6

Slobodkin MR, Elazar Z. 2013. The Atg8 family: multifunctional ubiquitin-like key regulators of autophagy. Essays Biochem 55:51–64. doi:10.1042/bse0550051

Sora V, Papaleo E. 2019. Bcl-xL dynamics and cancer-associated mutations under the lens of protein structure network and biomolecular simulations. bioarXiv. doi:10.1101/574699

Stadel D, Millarte V, Tillmann KD, Huber J, Tamin-Yecheskel BC, Akutsu M, Demishtein A, Ben-Zeev B, Anikster Y, Perez F, Dötsch V, Elazar Z, Rogov V, Farhan H, Behrends C. 2015. TECPR2 Cooperates with LC3C to Regulate COPII-Dependent ER Export. Mol Cell 60:89–104. doi:10.1016/j.molcel.2015.09.010

Stein A, Fowler DM, Hartmann-Petersen R, Lindorff-Larsen K. 2019. Biophysical and Mechanistic Models for Disease-Causing Protein Variants. Trends Biochem Sci 1–14. doi:10.1016/j.tibs.2019.01.003

Stolz A, Ernst A, Dikic I. 2014. Cargo recognition and trafficking in selective autophagy. Nat Cell Biol 16:495–501. doi:10.1038/ncb2979

Sugawara K, Suzuki NN, Fujioka Y, Mizushima N, Ohsumi Y, Inagaki F. 2004. The crystal structure of microtubule-associated protein light chain 3, a mammalian homologue of Saccharomyces cerevisiae Atg8. Genes to Cells 9:611–618. doi:10.1111/j.1356-9597.2004.00750.x

Sultan MM, Wayment-Steele HK, Pande VS. 2018. Transferable neural networks for enhanced sampling of protein dynamics.

Suzuki H, Tabata K, Morita E, Kawasaki M, Kato R, Dobson RCJ, Yoshimori T, Wakatsuki S. 2014. Structural basis of the autophagy-related LC3/Atg13 LIR complex: Recognition and interaction mechanism. Structure 22:47–58. doi:10.1016/j.str.2013.09.023

Tang JY, Hsi E, Huang YC, Hsu NCH, Chu PY, Chai CY. 2013. High LC3 expression correlates with poor survival in patients with oral squamous cell carcinoma. Hum Pathol 44:2558–2562. doi:10.1016/j.humpath.2013.06.017

Tanida I, Ueno T, Kominami E. 2008. LC3 and autophagy - Methods in Molecular Biology. Methods Mol Biol 445:77–88. doi:10.1007/978-1-59745-157-4-4

Tanida I, Ueno T, Kominami E. 2004. Human light chain 3/MAP1LC3B Is cleaved at its carboxyl-terminal Met 121 to expose Gly120 for lipidation and targeting to autophagosomal membranes. J Biol Chem 279:47704–47710. doi:10.1074/jbc.M407016200

Tate JG, Bamford S, Jubb HC, Sondka Z, Beare DM, Bindal N, Boutselakis H, Cole CG, Creatore C, Dawson E, Fish P, Harsha B, Hathaway C, Jupe SC, Kok CY, Noble K, Ponting L, Ramshaw CC, Rye CE, Speedy HE, Stefancsik R, Thompson SL, Wang S, Ward S, Campbell PJ, Forbes SA. 2019. COSMIC: the Catalogue Of Somatic Mutations In Cancer. Nucleic Acids Res 47:D941–D947. doi:10.1093/nar/gky1015

Thorburn A. 2014. Autophagy and Its Effects: Making Sense of Double-Edged Swords. PLoS Biol 12:10–13. doi:10.1371/journal.pbio.1001967

Thorburn A, Thamm DH, Gustafson DL. 2014. Autophagy and cancer therapy. Mol Pharmacol 85:830–838. doi:10.4161/auto.2.2.2463

Thukral L, Sengupta D, Ramkumar A, Murthy D, Agrawal N, Gokhale RS. 2015. The Molecular Mechanism Underlying Recruitment and Insertion of Lipid-Anchored LC3 Protein into Membranes. Biophys J 109:2067–2078. doi:10.1016/j.bpj.2015.09.022

Tiberti M, Invernizzi G, Lambrughi M, Inbar Y, Schreiber G, Papaleo E. 2014. PyInteraph : a framework for the analysis of interaction networks in structural ensembles of proteins. J Chem Inf Model 54:1537–1551.

Tiberti M, Papaleo E, Bengtsen T, Boomsma W, Lindorff-Larsen K. 2015. ENCORE: software for quantitative ensemble comparison. PLoS Comput Biol 11:e1004415.

Tokuriki N, Stricher F, Schymkowitz J, Serrano L, Tawfik DS. 2007. The Stability Effects of Protein Mutations Appear to be Universally Distributed. J Mol Biol 369:1318–1332. doi:10.1016/j.jmb.2007.03.069

Towers CG, Thorburn A. 2016. Therapeutic Targeting of Autophagy. EBioMedicine 14:15–23. doi:10.1016/j.ebiom.2016.10.034

Unan H, Yildirim A, Tekpinar M. 2015. Opening mechanism of adenylate kinase can vary according to selected molecular dynamics force field. J Comput Aided Mol Des 29:655–65. doi:10.1007/s10822-015-9849-0

Van Roey K, Dinkel H, Weatheritt RJ, Gibson TJ, Davey NE. 2013. The switches.ELM resource: a compendium of conditional regulatory interaction interfaces. Sci Signal 6:rs7. doi:10.1126/scisignal.2003345

Verkhivker GM. 2019. Biophysical simulations and structure-based modeling of residue interaction networks in the tumor suppressor proteins reveal functional role of cancer mutation hotspots in molecular communication. Biochim Biophys Acta - Gen Subj 1863:210–225. doi:10.1016/j.bbagen.2018.10.009

Viloria JS, Allega MF, Lambrughi M, Papaleo E. 2017. An optimal distance cutoff for contact-based Protein Structure Networks using side-chain centers of mass. Sci Rep 7:1–11. doi:10.1038/s41598-017-01498-6

von Muhlinen N, Akutsu M, Ravenhill BJ, Foeglein Á, Bloor S, Rutherford TJ, Freund SM V, Komander D, Randow F. 2012. LC3C, Bound Selectively by a Noncanonical LIR Motif in NDP52, Is Required for Antibacterial Autophagy. Mol Cell 48:329–342. doi:10.1016/j.molcel.2012.08.024

Vuillon L, Lesieur C. 2015. From local to global changes in proteins: a network view. Curr Opin Struct Biol 31:1–8. doi:10.1016/j.sbi.2015.02.015

Wagner SA, Beli P, Weinert BT, Nielsen ML, Cox J, Mann M, Choudhary C. 2011. A Proteome-wide, Quantitative Survey of In Vivo Ubiquitylation Sites Reveals Widespread Regulatory Roles. Mol Cell Proteomics 10:M111.013284. doi:10.1074/mcp.M111.013284

White E. 2012. Deconvoluting the context-dependent role for autophagy in cancer. Nat Rev Cancer. doi:10.1038/nrc3262

White E, DiPaola RS. 2009. The double-edged sword of autophagy modulation in cancer. Clin Cancer Res 15:5308–5316. doi:10.1158/1078-0432.CCR-07-5023

Wild P, McEwan DG, Dikic I. 2014. The LC3 interactome at a glance. J Cell Sci 127:3–9. doi:10.1242/jcs.140426

Wilkinson DS, Hansen M. 2015. LC3 is a novel substrate for the mammalian Hippo kinases, STK3/STK4. Autophagy 11:856–857. doi:10.1080/15548627.2015.1017197

Wilkinson DS, Jariwala JS, Anderson E, Mitra K, Meisenhelder J, Chang JT, Ideker T, Hunter T, Nizet V, Dillin A, Hansen M. 2015. Phosphorylation of LC3 by the hippo kinases STK3/STK4 is essential for autophagy. Mol Cell 57:55–68. doi:10.1016/j.molcel.2014.11.019

Wu DH, Jia CC, Chen J, Lin ZX, Ruan DY, Li X, Lin Q, Min-Dong, Ma XK, Wan XB, Cheng N, Chen ZH, Xing YF, Wu XY, Wen JY. 2014. Autophagic LC3B overexpression correlates with malignant progression and predicts a poor prognosis in hepatocellular carcinoma. Tumor Biol 35:12225– 12233. doi:10.1007/s13277-014-2531-7

Wu S, Sun C, Tian D, Li Y, Gao X, He S, Li T. 2015. Expression and clinical significances of Beclin1, LC3 and mTOR in colorectal cancer. Int J Clin Exp Pathol 8:3882–3891.

Xin J, Mark A, Afrasiabi C, Tsueng G, Juchler M, Gopal N, Stupp GS, Putman TE, Ainscough BJ, Griffith OL, Torkamani A, Whetzel PL, Mungall CJ, Mooney SD, Su AI, Wu C. 2016. High-performance web services for querying gene and variant annotation. Genome Biol 17:1–7. doi:10.1186/s13059-016-0953-9

Xue Y, Ren J, Gao X, Jin C, Wen L, Yao X. 2008. GPS 2.0, a Tool to Predict Kinase-specific Phosphorylation Sites in Hierarchy. Mol Cell Proteomics 7:1598–1608. doi:10.1074/mcp.m700574-mcp200

Yang A, Pantoom S, Wu Y-W. 2017. Elucidation of the anti-autophagy mechanism of the Legionella effector RavZ using semisynthetic LC3 proteins. Elife 6:1–23. doi:10.7554/elife.23905

Yao D, Wang P, Zhang J, Fu L, Ouyang L. 2016. Deconvoluting the relationships between autophagy and metastasis for potential cancer therapy. Apoptosis. doi:10.1007/s10495-016-1237-2

Zaffagnini G, Martens S. 2016. Mechanisms of Selective Autophagy. J Mol Biol 428:1714–1724. doi:10.1016/j.jmb.2016.02.004

Zhao H, Yang M, Zhao B. 2017. Beclin 1 and LC3 as predictive biomarkers for metastatic colorectal carcinoma. Oncotarget 8:59058–59067. doi:10.18632/oncotarget.19939

Zhao H, Yang M, Zhao J, Wang J, Zhang Y, Zhang Q. 2013. High expression of LC3B is associated with progression and poor outcome in triple-negative breast cancer. Med Oncol 30. doi:10.1007/s12032-013-0475-1

Zhao L, Liu S, Xu J, Li W, Duan G, Wang H, Yang H, Yang Z, Zhou R. 2017. A new molecular mechanism underlying the EGCG-mediated autophagic modulation of AFP in HepG2 cells. Cell Death Dis 8. doi:10.1038/cddis.2017.563

Zhou CY, Jiang F, Wu YD. 2015. Residue-specific force field based on protein coil library. RSFF2: Modification of AMBER ff99SB. J Phys Chem B 119:1035–1047. doi:10.1021/jp5064676

