## Supplementary Table S1 for "The conformational and mutational landscape of the ubiquitin-like marker for the autophagosome formation in cancer"

**Table S1. Pairwise Wilcox test (MWU) for the distribution of the resolution R values** for each MD ensemble of LC3B_7-116_ collected using ten different state-of-the-art force fields. The (adjusted) p-values of the force field pairs that have p-values higher than 0.05 are not significantly different from each other under the Holm-Bonferroni correction.

|  | CHARMM22* | CHARMM27 | CHARMM36 | ff14SB | ff99SB*-ILDN | ff99SB*-ILDN-Q | ff99SBnmr1 | RSFF2 | a99SB-disp | RSFF1 |
| --- | --- | --- | --- | --- | --- | --- | --- | --- | --- | --- |
| CHARMM22* | - | - | - | - | - | - | - | - | - | - |
| CHARMM27 | 0,13 | - | - | - | - | - | - | - | - | - |
| CHARMM36 | 0,94 | p<0.05 | - | - | - | - | - | - | - | - |
| ff14SB | p<0.05 | 0,49 | p<0.05 | - | - | - | - | - | - | - |
| ff99SB*-ILDN | 1,00 | p<0.05 | 1,00 | p<0.05 | - | - | - | - | - | - |
| ff99SB*-ILDN-Q | p<0.05 | p<0.05 | p<0.05 | 0,49 | p<0.05 | - | - | - | - | - |
| ff99SBnmr1 | p<0.05 | p<0.05 | p<0.05 | p<0.05 | p<0.05 | 0,49 | - | - | - | - |
| RSFF2 | 0,37 | 1,00 | p<0.05 | 0,49 | 0,08 | p<0.05 | p<0.05 | - | - | - |
| a99SB-disp | p<0.05 | p<0.05 | p<0.05 | p<0.05 | p<0.05 | 0,49 | 1,00 | p<0.05 | - | - |
| RSFF1 | p<0.05 | p<0.05 | p<0.05 | p<0.05 | p<0.05 | p<0.05 | 0,08 | p<0.05 | p<0.05 | - |
