## Supplementary TableS2 for "The conformational and mutational landscape of the ubiquitin-like marker for the autophagosome formation in cancer"

**Table S2. Root-Mean-Square-Error (RMSE) between the predicted chemical shift values from the MD simulations and the experimentally measured chemical shifts**

| **RMSE (in units of ppm)** |  |  |  |  |  |
| --- | --- | --- | --- | --- | --- |
| **Force Field** | **C** | **CA** | **N** | **H** | **HA** |
| **PPM_One Error** | **1.44** | **0.90** | **2.31** | **0.43** | **0.24** |
| **CHARMM22*** | 0.98 | 1.03 | 2.35 | 0.51 | 0.26 |
| **CHARMM27** | 0.94 | 0.95 | 2.55 | 0.49 | 0.26 |
| **CHARMM36** | 0.97 | 0.93 | 2.60 | 0.57 | 0.27 |
| **ff14SB** | 1.07 | 0.99 | 2.50 | 0.53 | 0.26 |
| **ff99SB*-ILDN** | 1.04 | 1.04 | 2.53 | 0.57 | 0.26 |
| **ff99SB*-ILDN-Q** | 0.99 | 1.02 | 2.45 | 0.56 | 0.26 |
| **ff99SBnmr1** | 1.00 | 0.87 | 2.45 | 0.49 | 0.25 |
| **RSFF2** | 1.08 | 1.13 | 2.82 | 0.57 | 0.33 |
| **a99SB-disp** | 1.01 | 0.93 | 2.65 | 0.54 | 0.27 |
| **RSFF1** | 1.15 | 1.23 | 2.97 | 0.62 | 0.33 |
