## Supplementary figures and images for "The conformational and mutational landscape of the ubiquitin-like marker for the autophagosome formation in cancer"

### Supplementary File S1

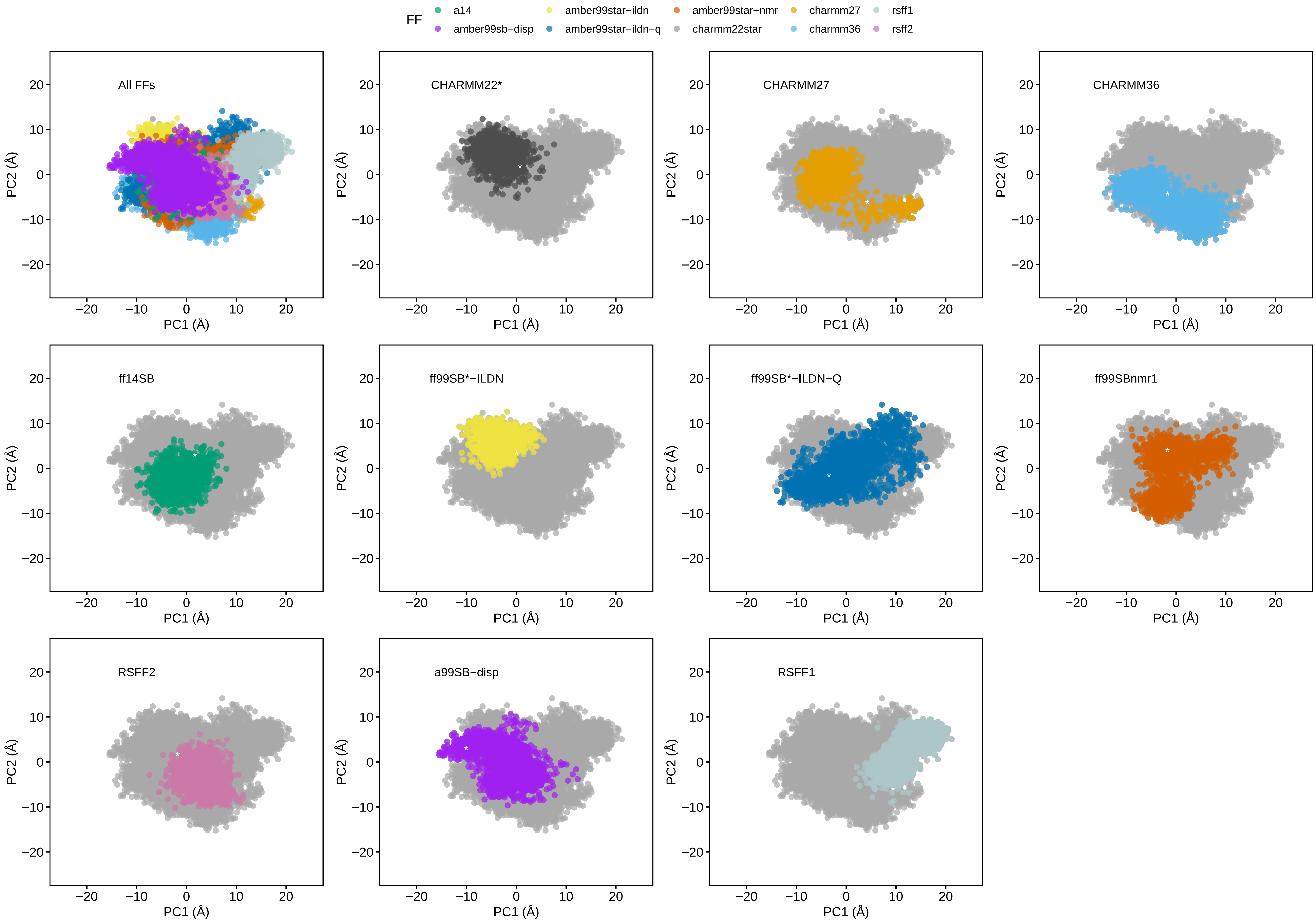

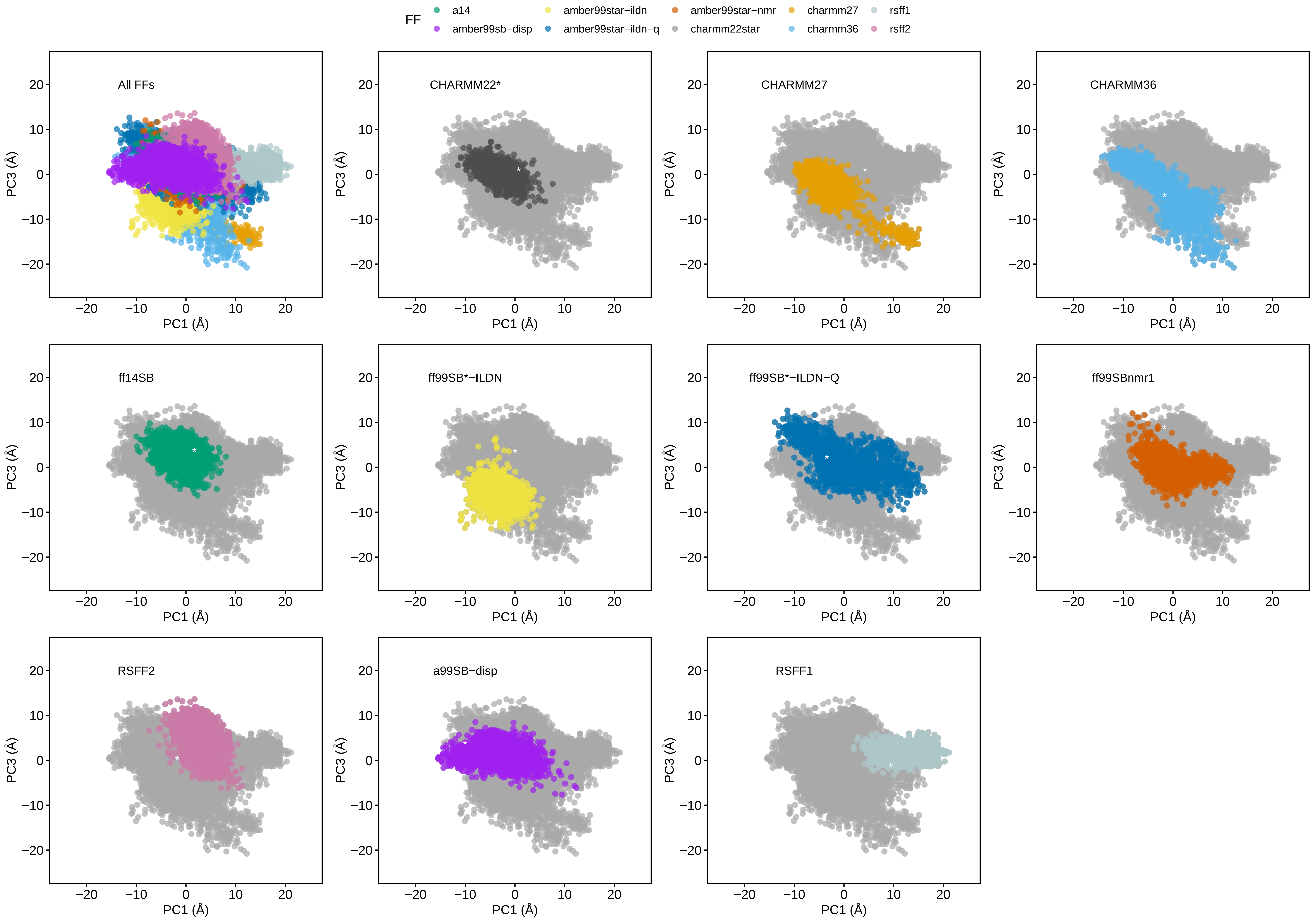

### Supplementary File S2

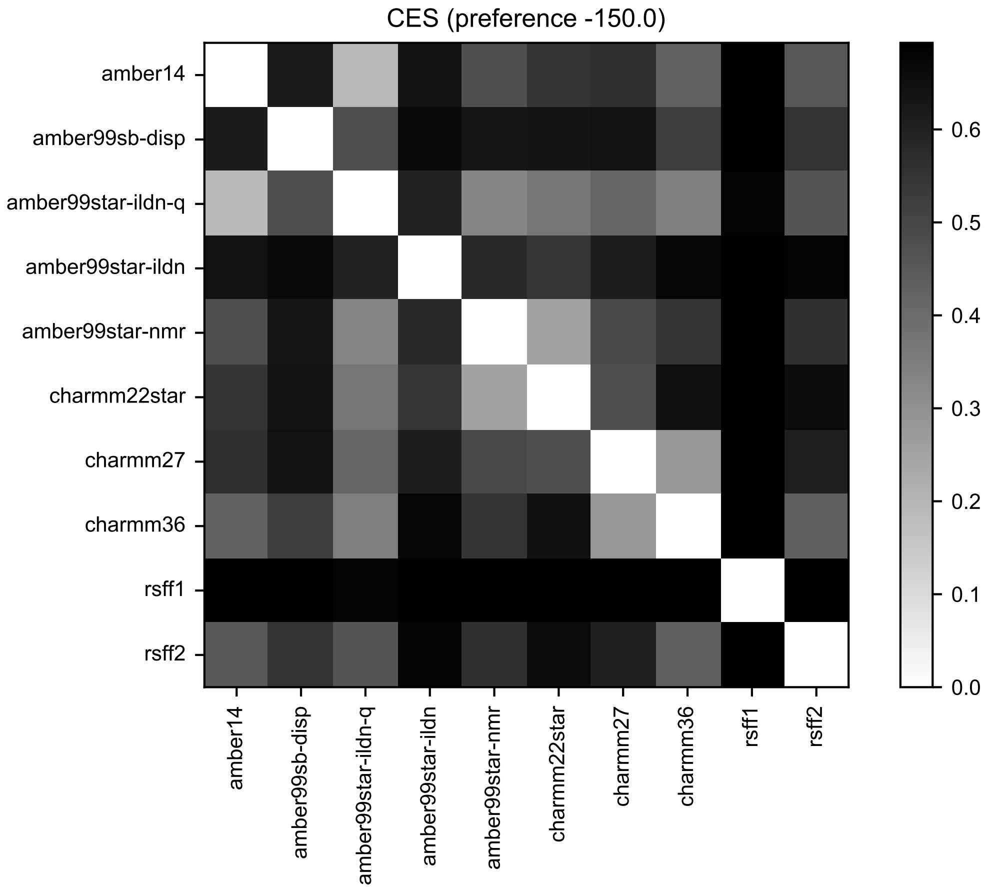

### Supplementary File S8

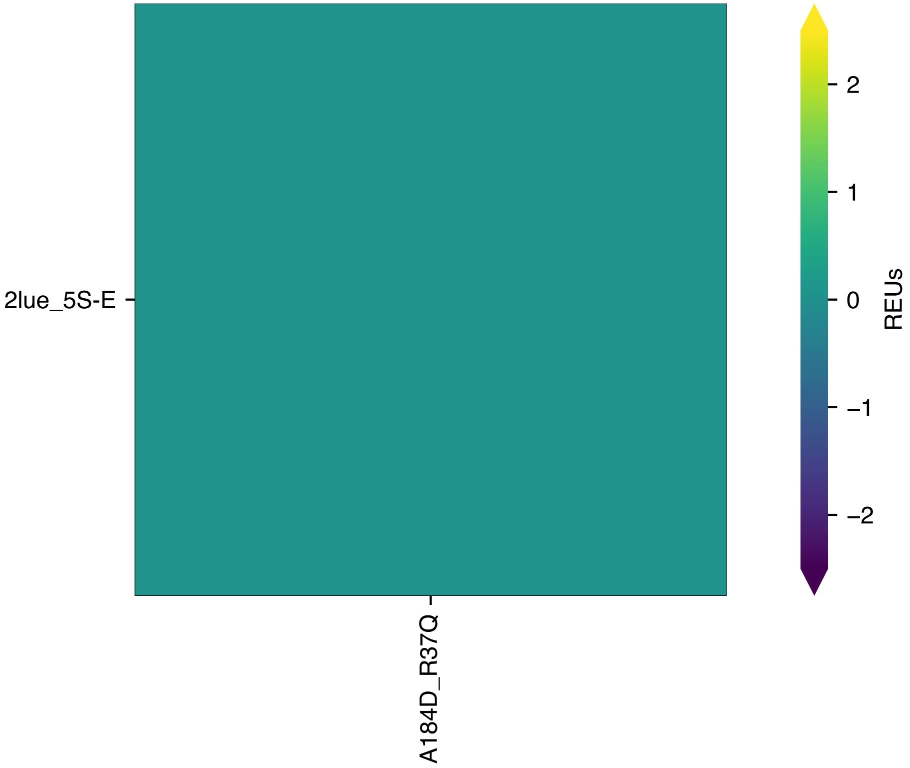

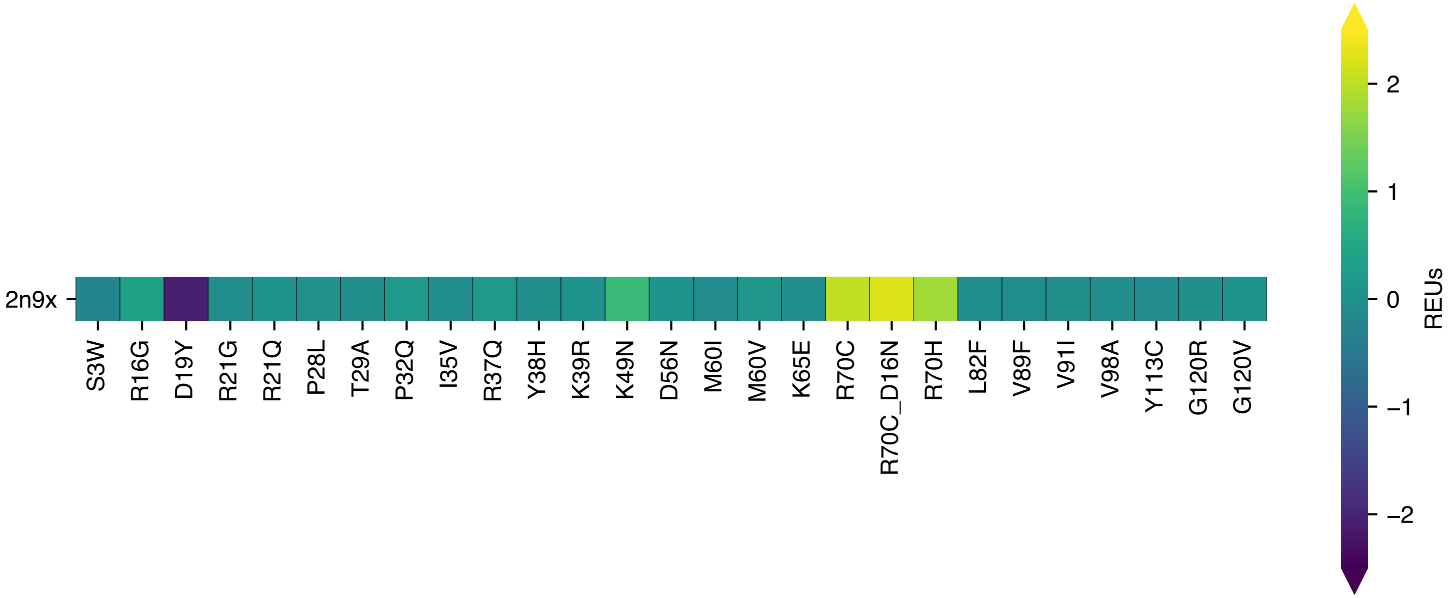

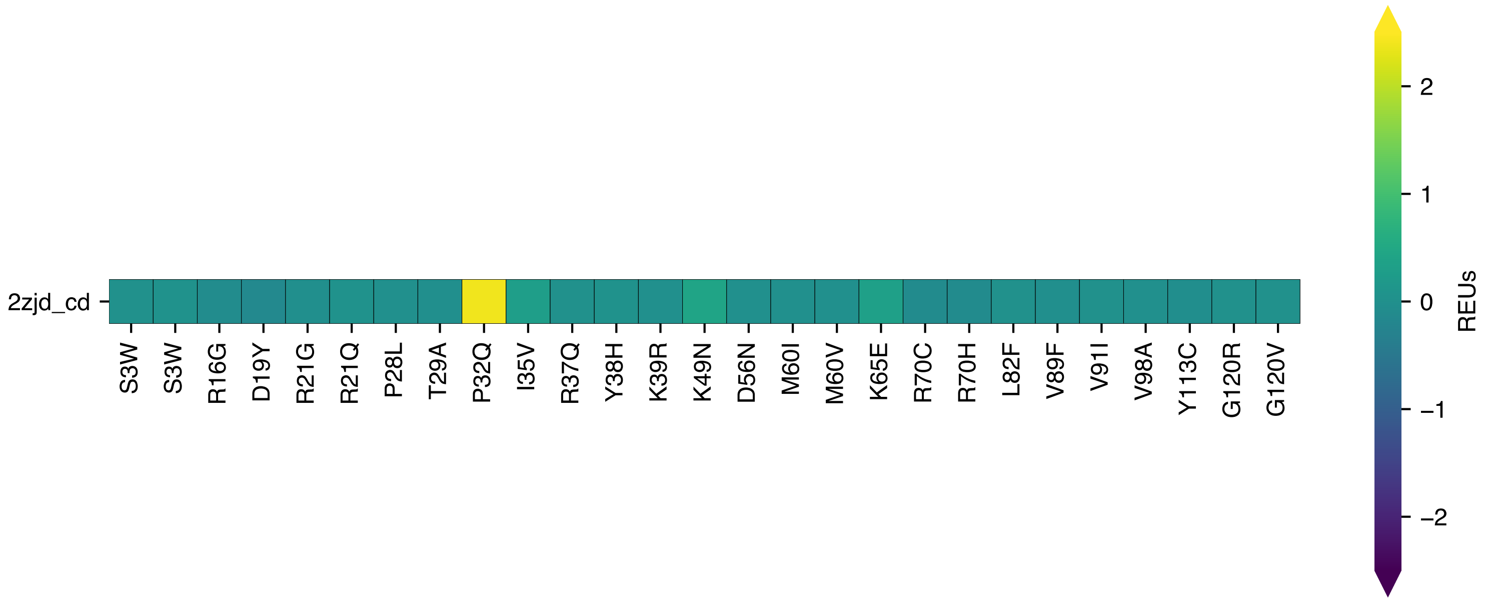

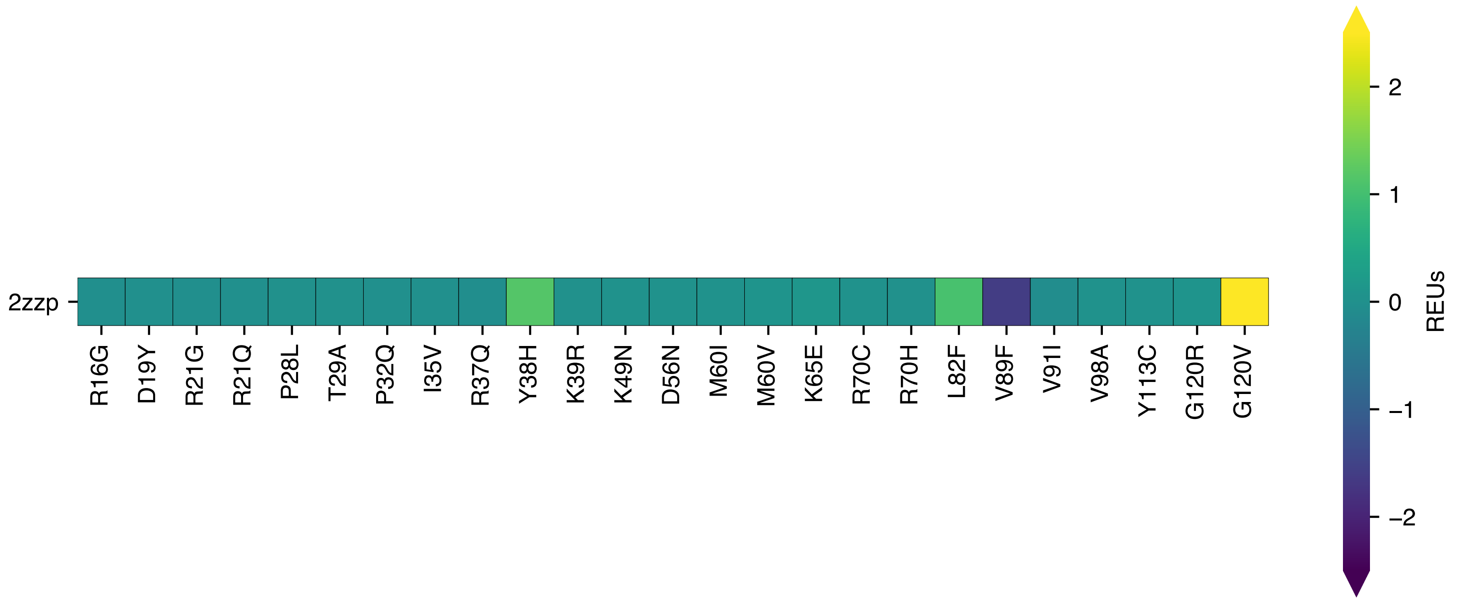

### Supplementary File S9

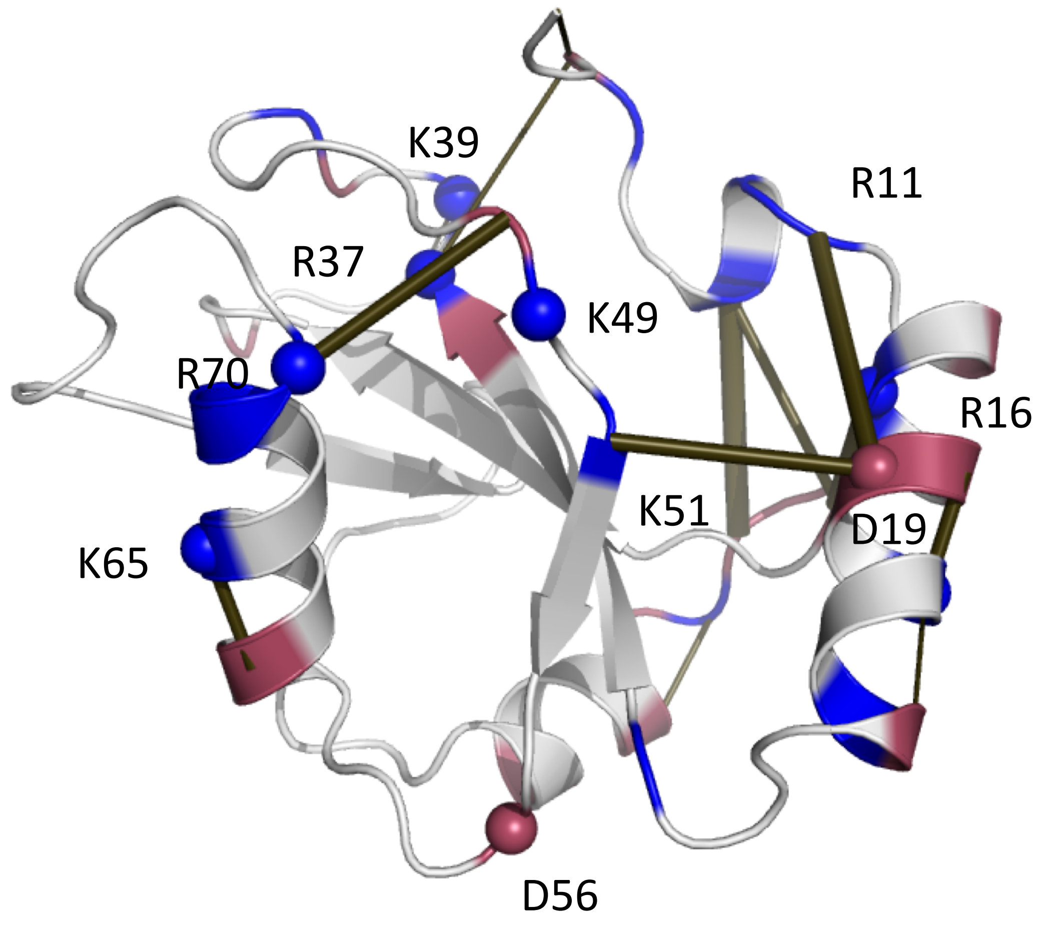
