## Supplementary File S4 for "The conformational and mutational landscape of the ubiquitin-like marker for the autophagosome formation in cancer"

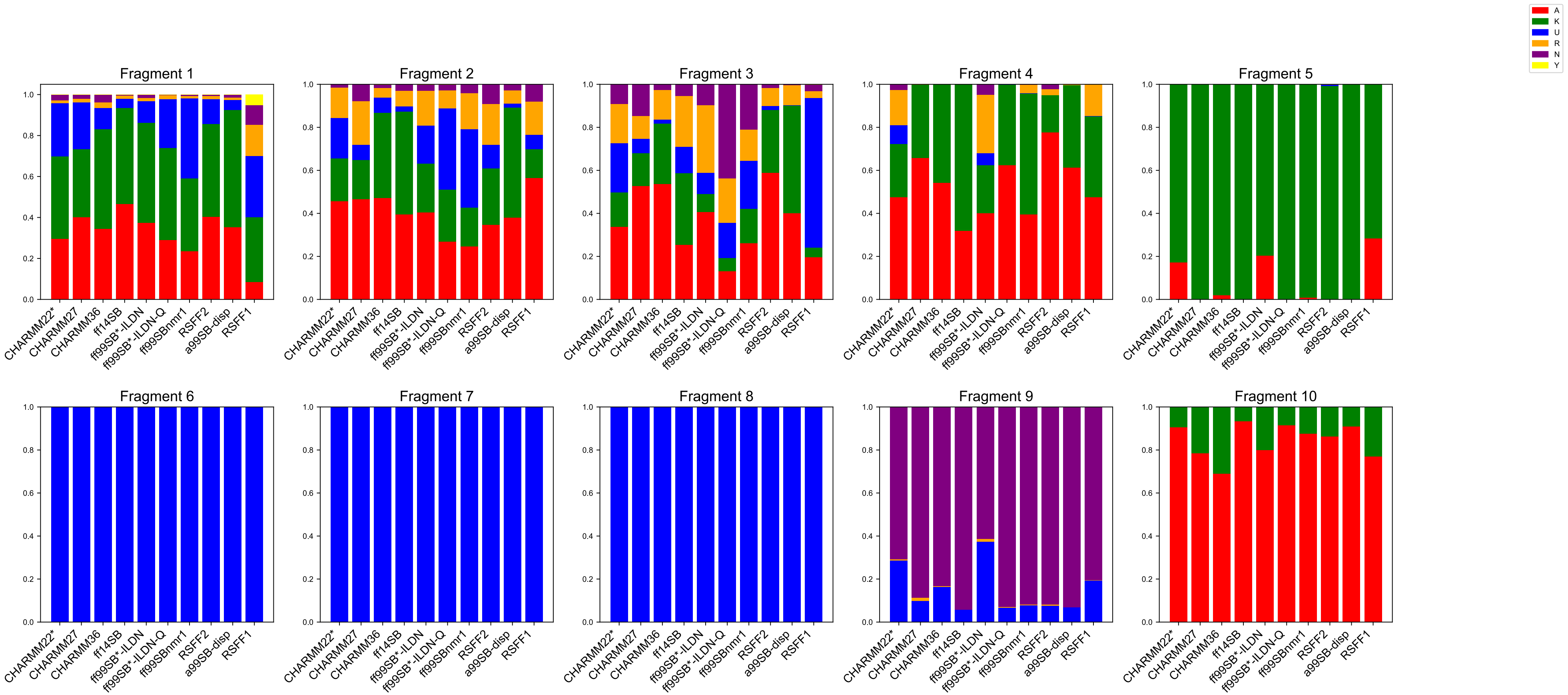
Fragment 1
Fragment 2
Fragment 3
Fragment 4
Fragment 5
Fragment 6
Fragment 7
Fragment 8
Fragment 9
Fragment 10
A
K
U
R
N
Y
CHARMM22*
CHARMM27
CHARMM36
ff14SB
ff99SB*-ILDN
ff99SB*-ILDN-Q
ff99SBnmr1
RSFF2
a99SB-disp
RSFF1
0.0
0.2
0.4
0.6
0.8
1.0

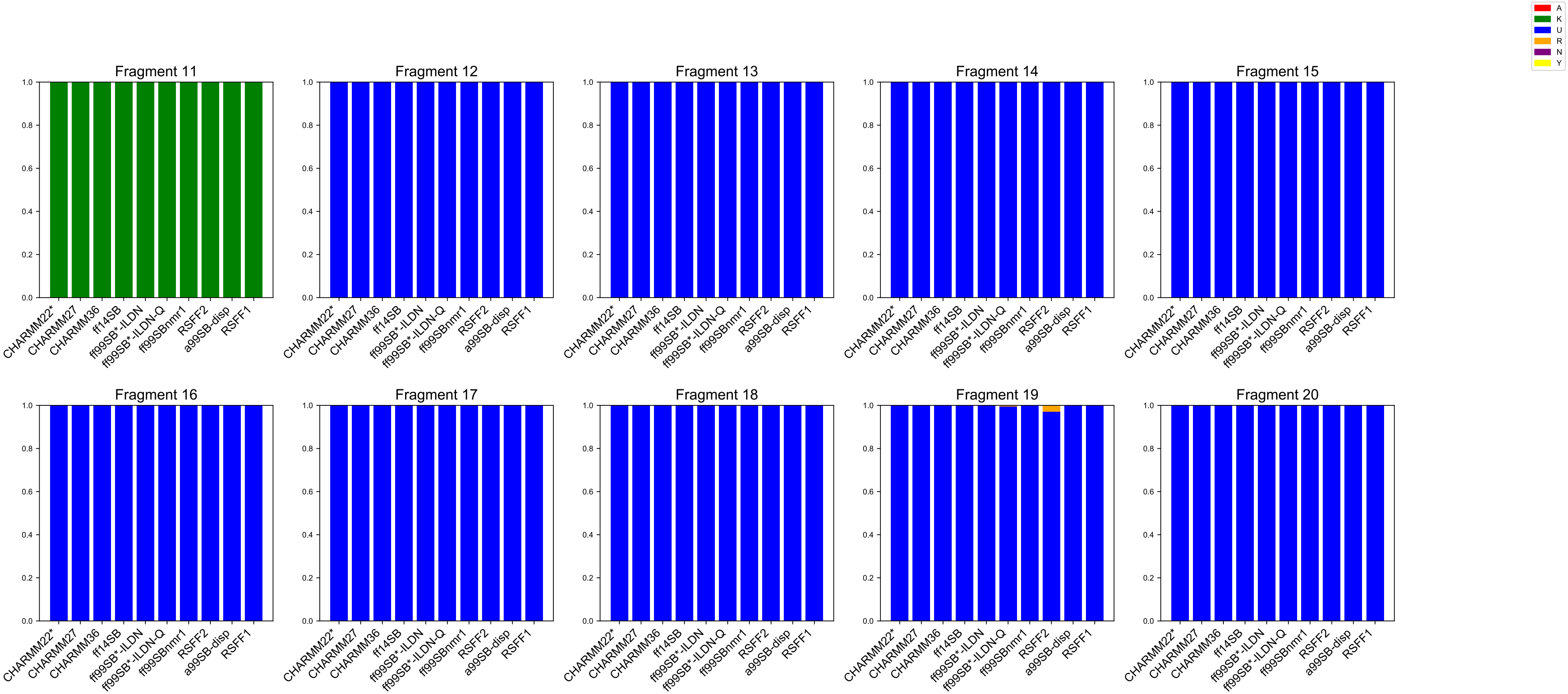

A
K
U
R
N
Y
Fragment 11
Fragment 12
Fragment 13
Fragment 14
Fragment 15
Fragment 16
Fragment 17
Fragment 18
Fragment 19
Fragment 20
0.0
0.2
0.4
0.6
0.8
1.0
CHARMM22*
CHARMM27
CHARMM36
ff14SB
ff99SB*-ILDN
ff99SB*-ILDN-Q
ff99SBnmr1
RSFF2
a99SB-disp
RSFF1

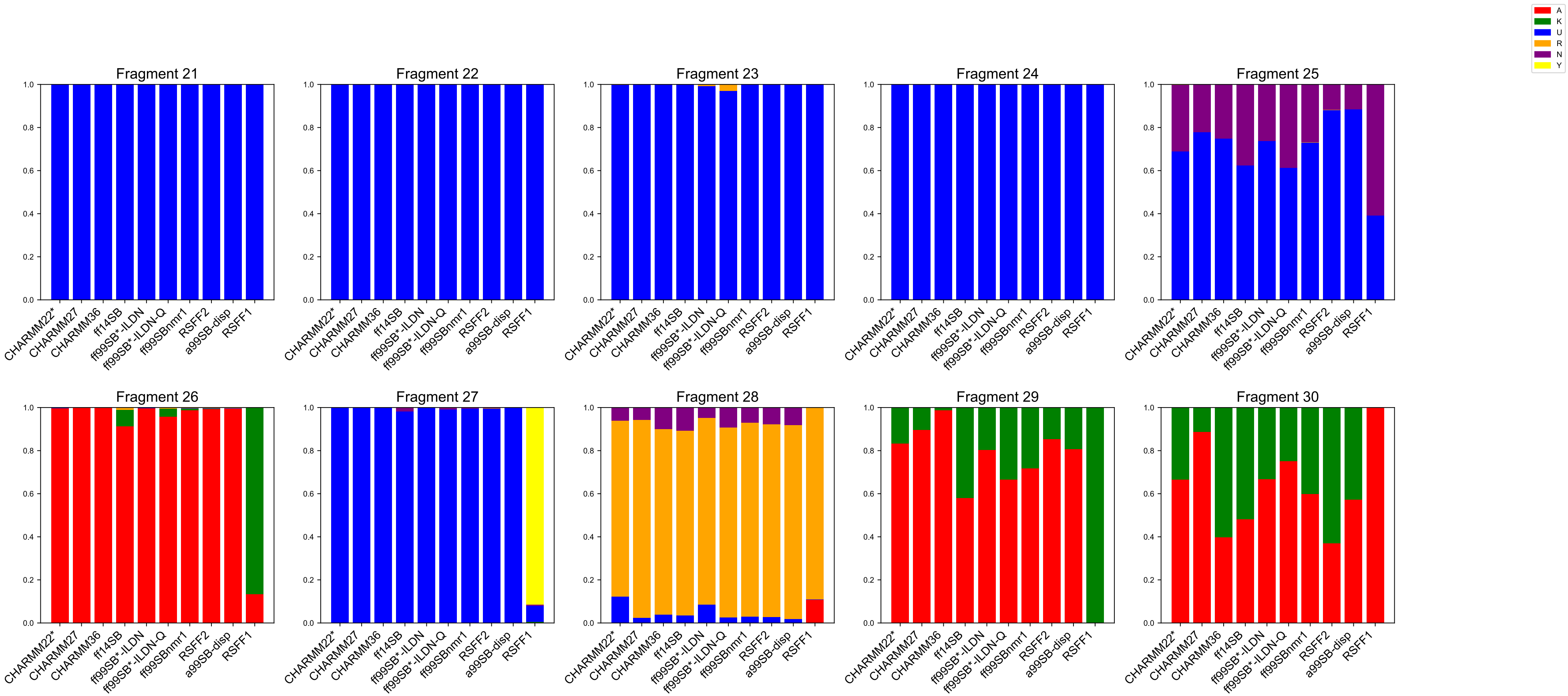
A
K
U
R
N
Y
Fragment 21
Fragment 22
Fragment 23
Fragment 24
Fragment 25
Fragment 26
Fragment 27
Fragment 28
Fragment 29
Fragment 30
1.0
0.8
0.6
0.4
0.2
0.0
CHARMM22*
CHARMM27
CHARMM36
ff14SB
ff99SB*-ILDN
ff99SB*-ILDN-Q
ff99SBnmr1
RSFF2
a99SB-disp
RSFF1

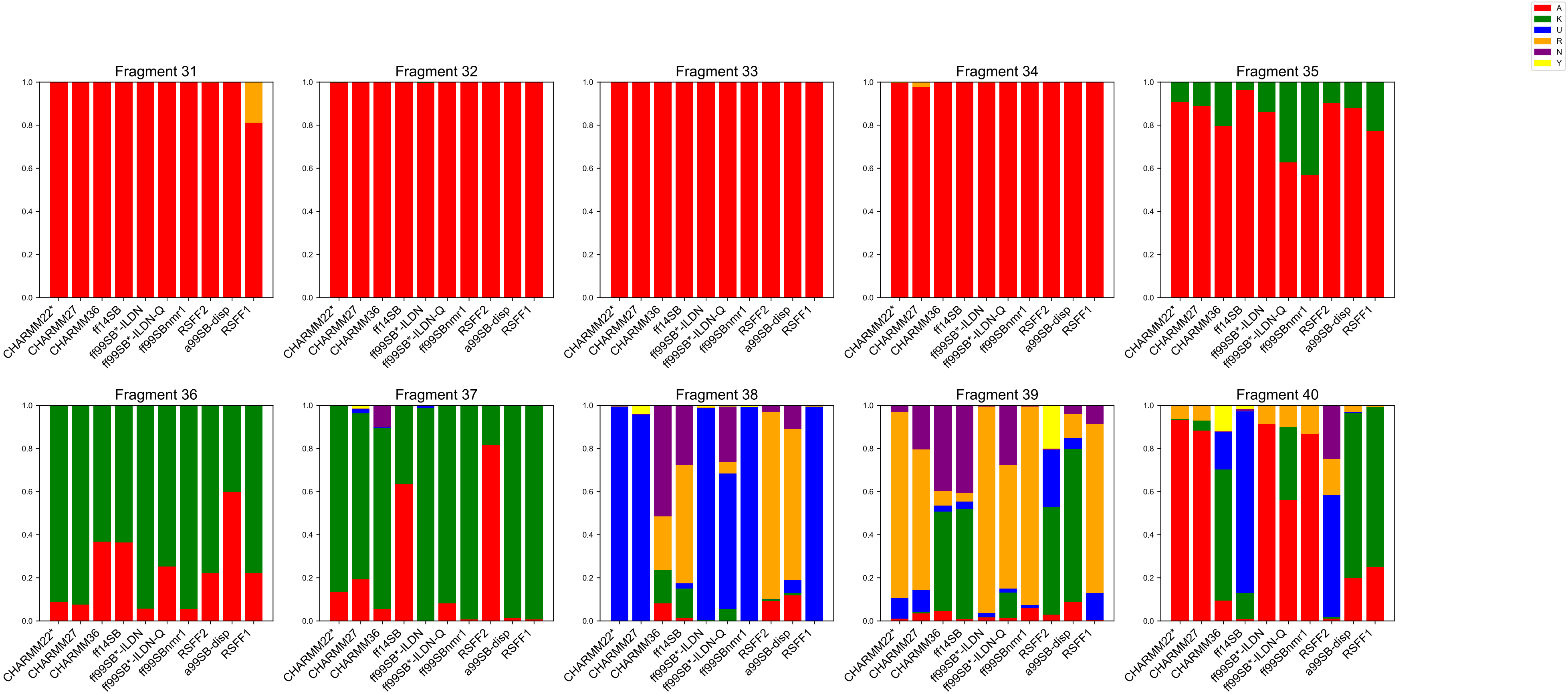
Fragment 31
Fragment 32
Fragment 33
Fragment 34
Fragment 35
Fragment 36
Fragment 37
Fragment 38
Fragment 39
Fragment 40
A
K
U
R
N
Y
0.0
0.2
0.4
0.6
0.8
1.0
CHARMM22*
CHARMM27
CHARMM36
ff14SB
ff99SB*-ILDN
ff99SB*-ILDN-Q
ff99SBnmr1
RSFF2
a99SB-disp
RSFF1

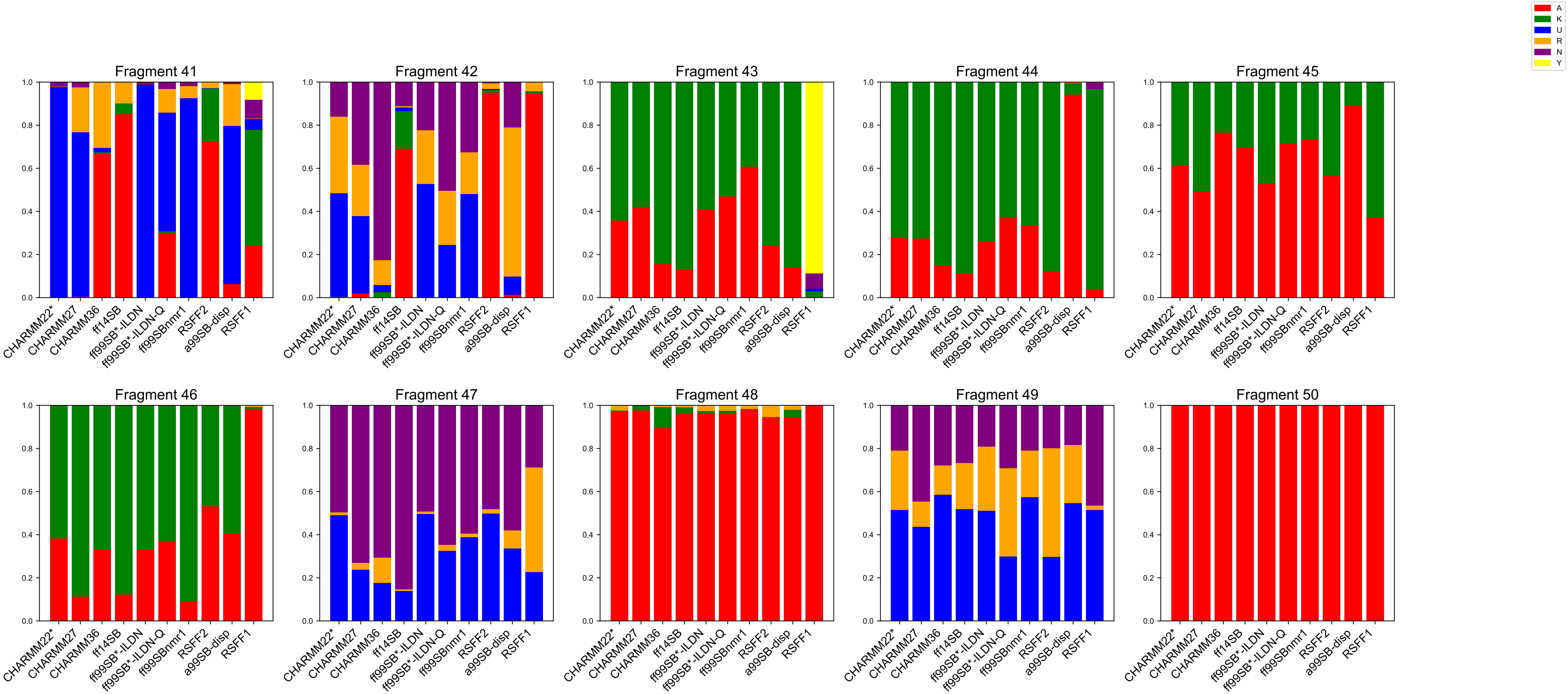

Fragment 41
Fragment 42
Fragment 43
Fragment 44
Fragment 45
Fragment 46
Fragment 47
Fragment 48
Fragment 49
Fragment 50
A
K
U
R
N
Y
0.0
0.2
0.4
0.6
0.8
1.0
CHARMM22*
CHARMM27
CHARMM36
ff14SB
ff99SB*-ILDN
ff99SB*-ILDN-Q
ff99SBnmr1
RSFF2
a99SB-disp
RSFF1

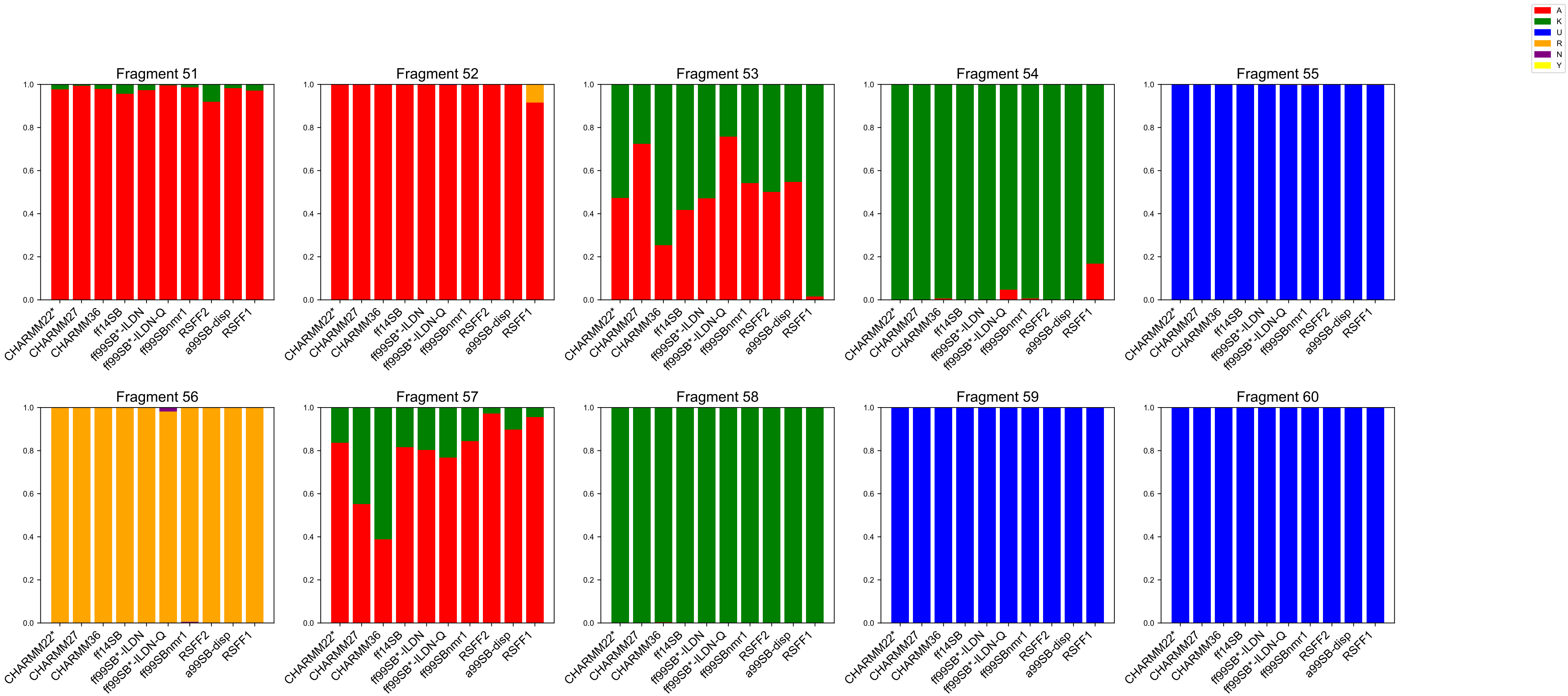

Fragment 51
Fragment 52
Fragment 53
Fragment 54
Fragment 55
Fragment 56
Fragment 57
Fragment 58
Fragment 59
Fragment 60
0.0
0.2
0.4
0.6
0.8
1.0
CHARMM22*
CHARMM27
CHARMM36
ff14SB
ff99SB*-ILDN
ff99SB*-ILDN-Q
ff99SBnmr1
RSFF2
a99SB-disp
RSFF1
A
K
U
R
N
Y

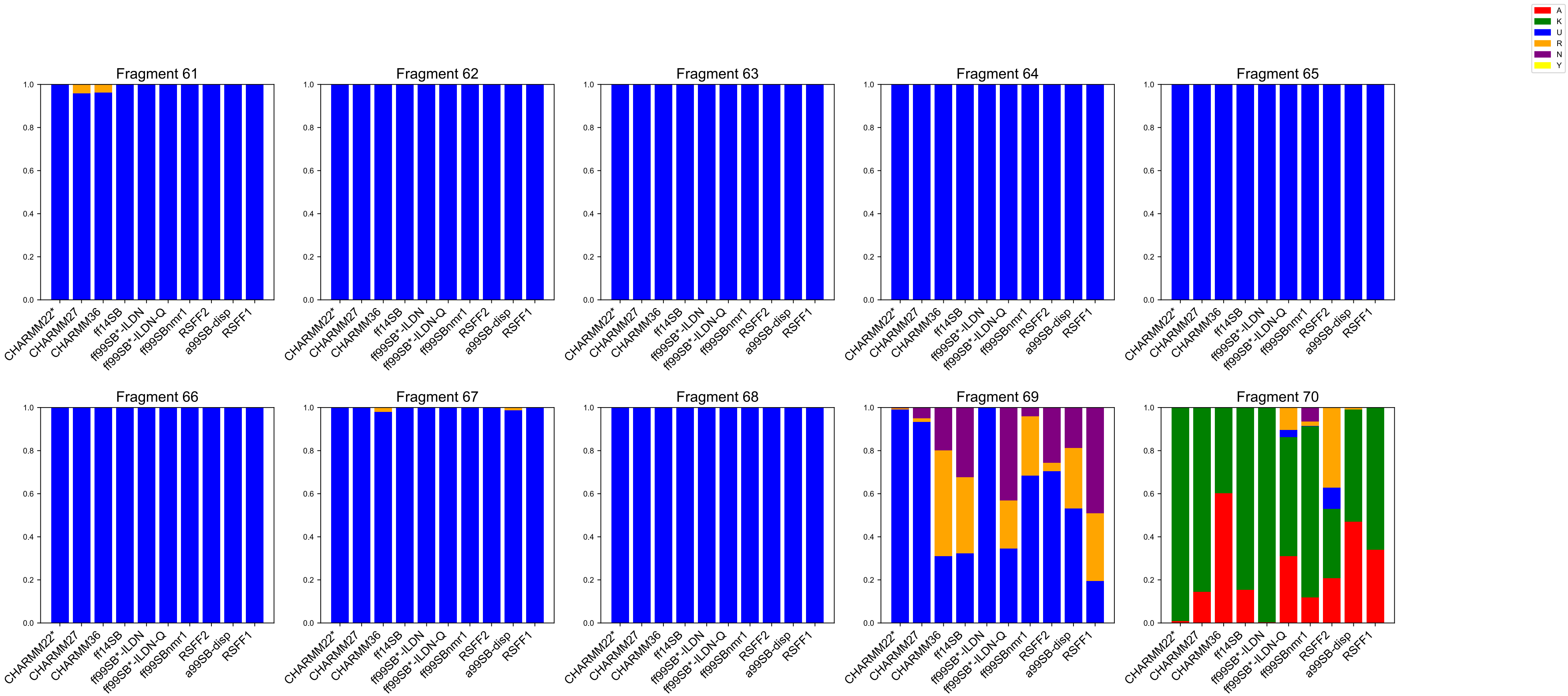
A
K
U
R
N
Y
Fragment 61
Fragment 62
Fragment 63
Fragment 64
Fragment 65
Fragment 66
Fragment 67
Fragment 68
Fragment 69
Fragment 70
1.0
0.8
0.6
0.4
0.2
0.0
CHARMM22*
CHARMM27
CHARMM36
ff14SB
ff99SB*-ILDN
ff99SB*-ILDN-Q
ff99SBnmr1
RSFF2
a99SB-disp
RSFF1

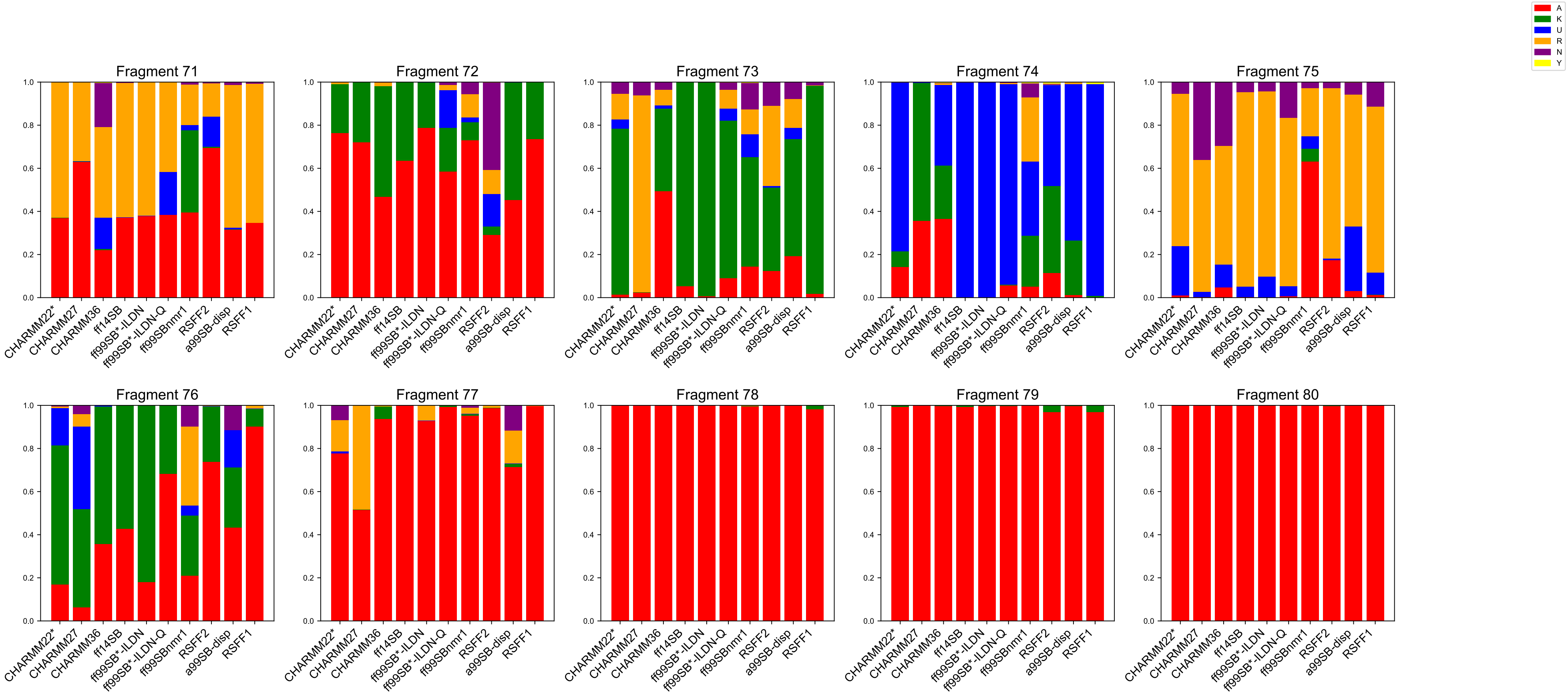
A
K
U
R
N
Y
Fragment 71
Fragment 72
Fragment 73
Fragment 74
Fragment 75
Fragment 76
Fragment 77
Fragment 78
Fragment 79
Fragment 80
0.0
0.2
0.4
0.6
0.8
1.0
CHARMM22*
CHARMM27
CHARMM36
ff14SB
ff99SB*-ILDN
ff99SB*-ILDN-Q
ff99SBnmr1
RSFF2
a99SB-disp
RSFF1

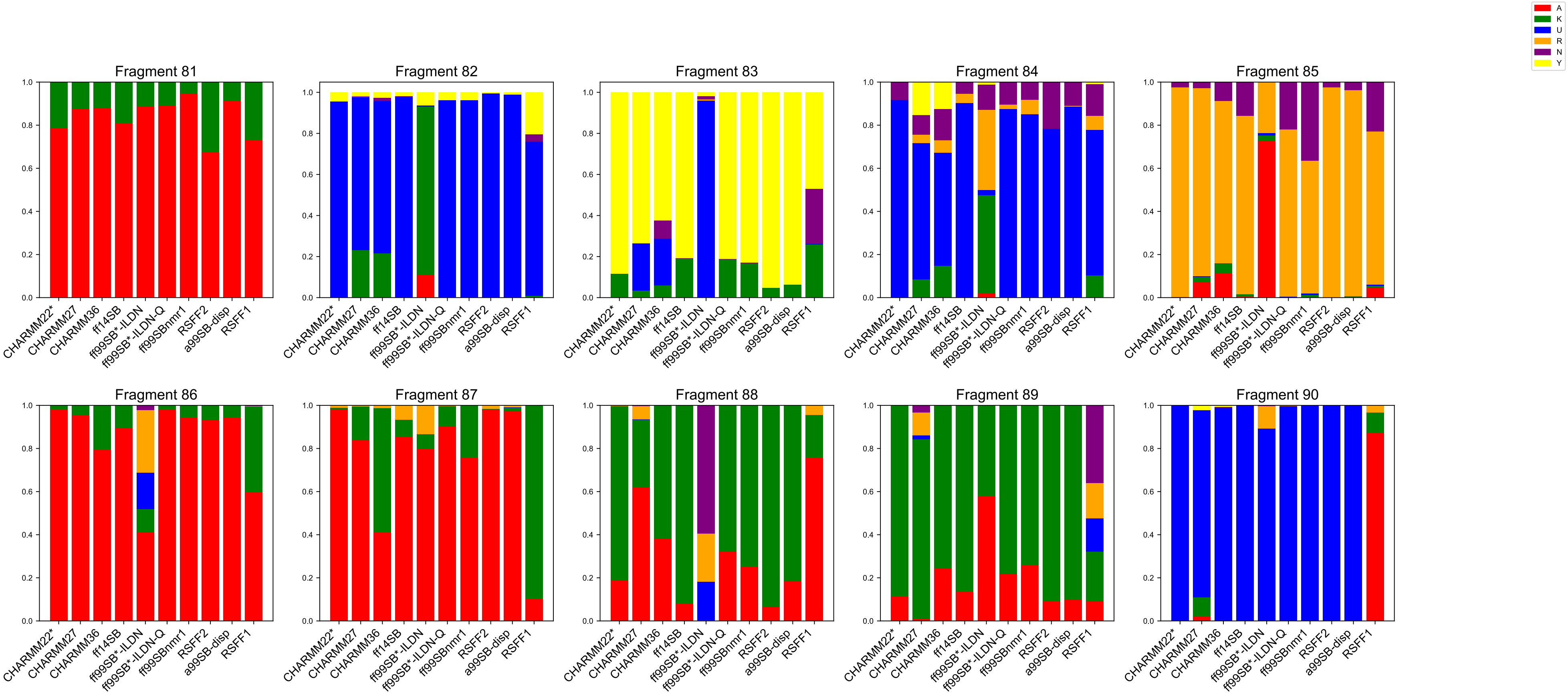
A
K
U
R
N
Y
Fragment 81
Fragment 82
Fragment 83
Fragment 84
Fragment 85
Fragment 86
Fragment 87
Fragment 88
Fragment 89
Fragment 90
0.0
0.2
0.4
0.6
0.8
1.0
CHARMM22*
CHARMM27
CHARMM36
ff14SB
ff99SB*-ILDN
ff99SB*-ILDN-Q
ff99SBnmr1
RSFF2
a99SB-disp
RSFF1

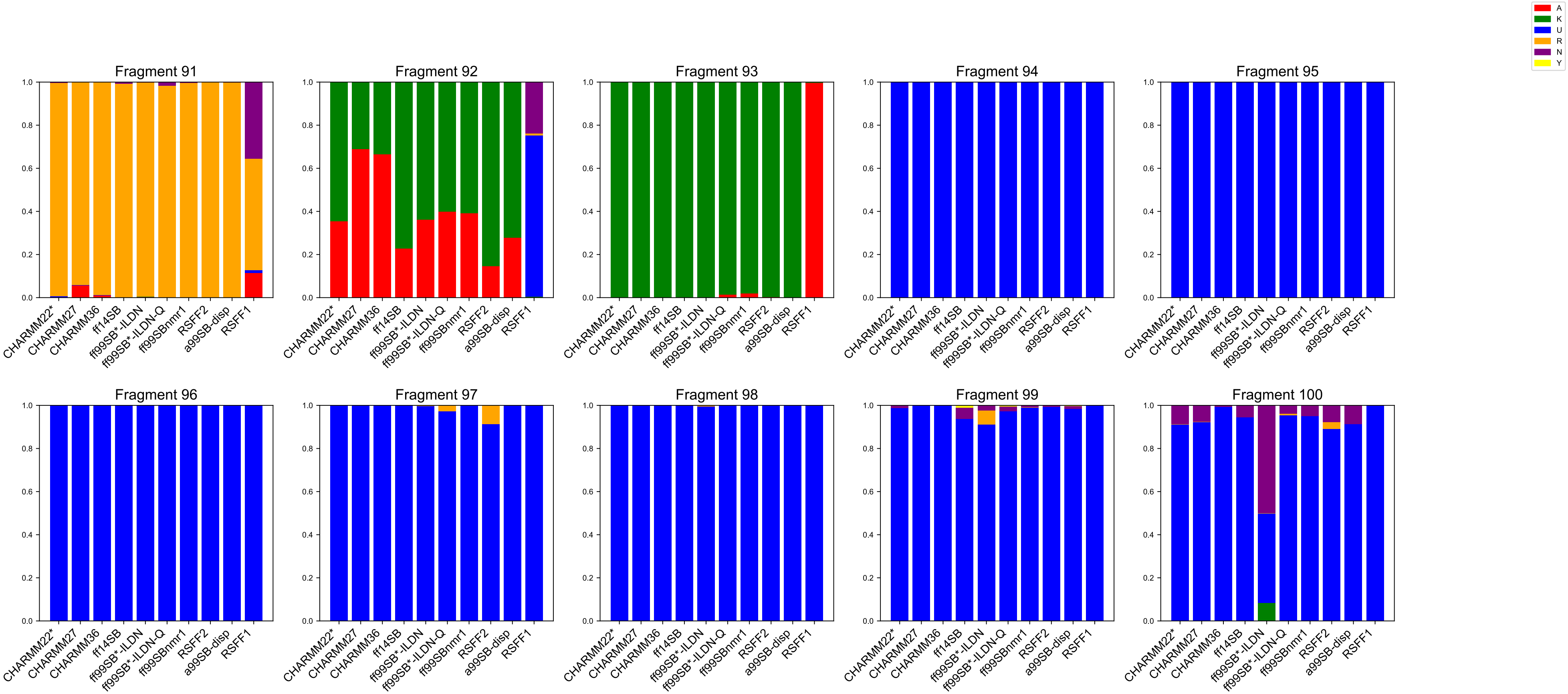
Fragment 91
Fragment 92
Fragment 93
Fragment 94
Fragment 95
Fragment 96
Fragment 97
Fragment 98
Fragment 99
Fragment 100
CHARMM22*
CHARMM27
CHARMM36
ff14SB
ff99SB*-ILDN
ff99SB*-ILDN-Q
ff99SBnmr1
RSFF2
a99SB-disp
RSFF1
A
K
U
R
N
Y

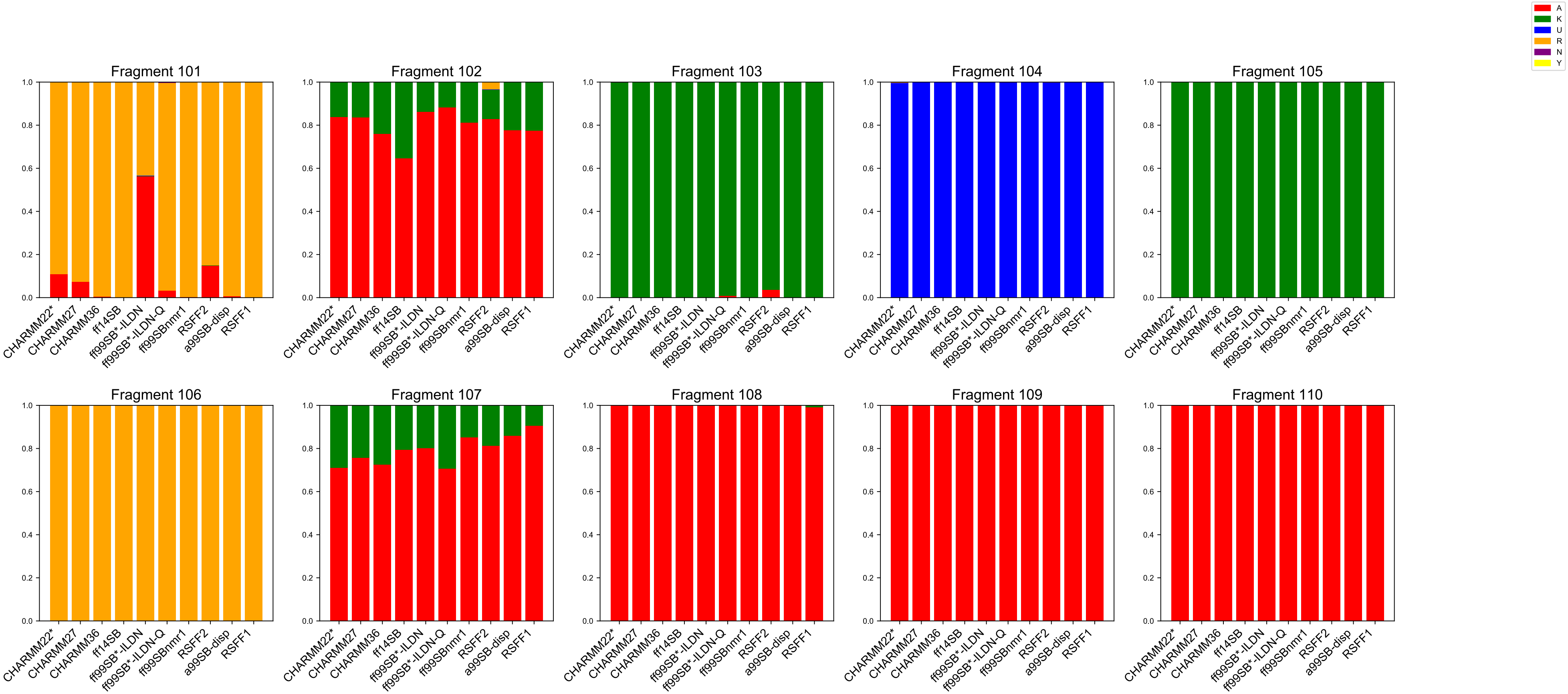
Fragment 101
Fragment 102
Fragment 103
Fragment 104
Fragment 105
Fragment 106
Fragment 107
Fragment 108
Fragment 109
Fragment 110
A
K
U
R
N
Y
0.0
0.2
0.4
0.6
0.8
1.0
CHARMM22*
CHARMM27
CHARMM36
ff14SB
ff99SB*-ILDN
ff99SB*-ILDN-Q
ff99SBnmr1
RSFF2
a99SB-disp
RSFF1

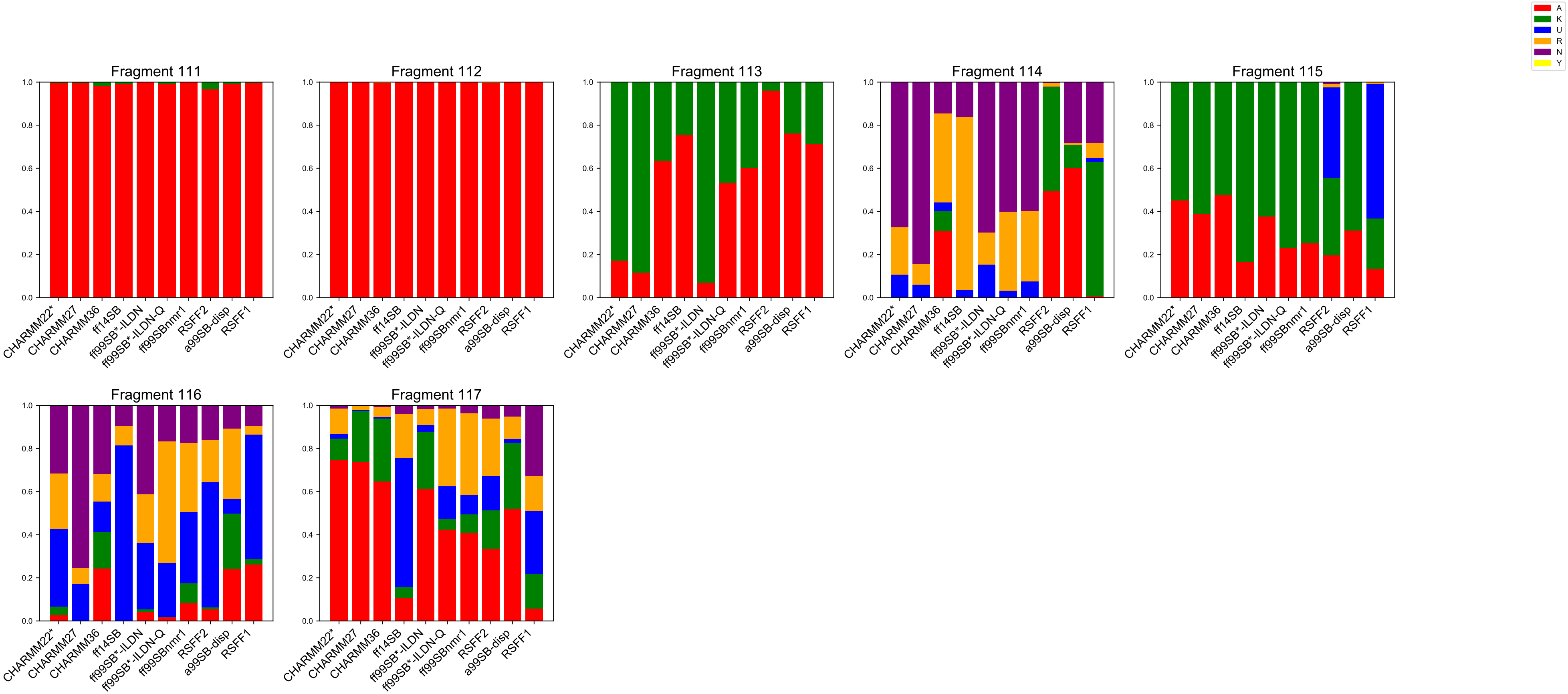

Fragment 111
Fragment 112
Fragment 113
Fragment 114
Fragment 115
Fragment 116
Fragment 117
CHARMM22*
CHARMM27
CHARMM36
ff14SB
ff99SB*-ILDN
ff99SB*-ILDN-Q
ff99SBnmr1
RSFF2
a99SB-disp
RSFF1
A
K
U
R
N
Y
