## Supplementary File S7 for "The conformational and mutational landscape of the ubiquitin-like marker for the autophagosome formation in cancer"

```
CLUSTAL O(1.2.4) multiple sequence alignment
sp|O95166|GBRAP_HUMAN      --------MKFVYKEEHPFEKRRSEGEKIRKKYPDRVPVIVEKAPKA-RIGDLDKKKYLV 51
sp|Q9BXW4|MLP3C_HUMAN      MPPPQKIPSVRPFKQRKSLAIRQEEVAGIRAKFPNKIPVVVERYPRETFLPPLDKTKFLV 60
sp|Q9GZQ8|MLP3B_HUMAN      ------MPSEKTFKQRRTFEQRVEDVRLIREQHPTKIPVIIERYKGEKQLPVLDKTKFLV 54
sp|Q9H492|MLP3A_HUMAN      ------MPSDRPFKQRRSFADRCKEVQQIRDQHPSKIPVIIERYKGEKQLPVLDKTKFLV 54
                                       :*:.: :  * .:   ** :.* ::**::*:      :  ***.*:**

sp|O95166|GBRAP_HUMAN      PSDLTVGQFYFLIRKRIHLRAEDALFFFVNNV-IPPTSATMGQLYQEHHEEDFFLYIAYS 110
sp|Q9BXW4|MLP3C_HUMAN      PQELTMTQFLSIIRSRMVLRATEAFYLLVNNKSLVSMSATMAEIYRDYKDEDGFVYMTYA 120
sp|Q9GZQ8|MLP3B_HUMAN      PDHVNMSELIKIIRRRLQLNANQAFFLLVNGHSMVSVSTPISEVYESEKDEDGFLYMVYA 114
sp|Q9H492|MLP3A_HUMAN      PDHVNMSELVKIIRRRLQLNPTQAFFLLVNQHSMVSVSTPIADIYEQEKDEDGFLYMVYA 114
                           *..:.: ::  :** *: *.  :*::::**   :   *: :.::*.. ::** *:*:.*:

sp|O95166|GBRAP_HUMAN      DESVYGL--------------------  117
sp|Q9BXW4|MLP3C_HUMAN      SQETFGCLESAAPRDGSSLEDRPCNPL  147
sp|Q9GZQ8|MLP3B_HUMAN      SQETFGMKLSV----------------  125
sp|Q9H492|MLP3A_HUMAN      SQETFGF--------------------  121
                           .:..:*
```
